# Engineering yeast for *de-novo* synthesis of the insect repellent - nepetalactone

**DOI:** 10.1101/2021.08.30.458239

**Authors:** Meghan E Davies, Daniel Tsyplenkov, Vincent J. J. Martin

## Abstract

While nepetalactone, the active ingredient in catnip, is a potent insect repellent, its low *in planta* accumulation limits its commercial viability as an alternative repellent. Here we describe a platform for *de novo* nepetalactone production in *Saccharomyces cerevisiae*, enabling sustainable and scalable production. Nepetalactone production required introduction of eight exogenous genes including the cytochrome P450 geraniol-8-hydroxylase, which represented the bottleneck of the heterologous pathway. Combinatorial assessment of geraniol-8-hydroxylase and cytochrome P450 reductase variants, as well as copy-number variations were used to overcome this bottleneck. We found that several reductases improved hydroxylation activity, with a higher geraniol-8-hydroxylase ratio further increasing 8-hydroxygeraniol titers. Another roadblock was the accumulation of an unwanted metabolite that implied inefficient channeling of carbon through the pathway. With the native yeast old yellow enzymes previously shown to use monoterpene intermediates as substrates, both homologs were deleted. These deletions increased 8-hydroxygeraniol yield, resulting in a final *de novo* accumulation of 3.10 mg/L/OD_600_ of nepetalactone from simple sugar in microtiter plates. Our pathway optimization will aid in the development of high yielding monoterpene *S. cerevisiae* strains.

---

Monoterpenes and their indole alkaloid derivatives (MIAs) are plant secondary metabolites known for their pharmaceutical activities^1^. Synthesis of the central MIA scaffold, strictosidine, proceeds via nepetalactol whose oxidized derivative, nepetalactone, is a potent insect repellent with higher repellency than N,N-diethyl-m-toluamide (DEET)^2^. With the growing population of DEET resistant mosquitoes^3^, nepetalactone could be an alternative source of insect repellant if a commercial scale production process is developed. Nepetalactone has higher *in planta* yields than MIAs, yet the mass market demand would preclude plant-sourced nepetalactone for commercial scale production due to prohibitive costs. Elucidation of the strictosidine biosynthetic pathway^4^ has enabled *de novo* production of nepetalactol^5^ and strictosidine^6^ in the budding yeast *S. cerevisiae*. Nepetalactone production was reported in yeast, however this was achieved using an incomplete pathway and supplying pathway intermediates^7^. Based on these initial studies, poor geraniol-8-hydroxylase (G8H) activity and undesired reactions consuming pathway intermediates, by both native and exogenous enzymes, were identified as limiting titers.

The G8H monooxygenase belongs to the cytochrome P450 (CYP) superfamily of hemoproteins. This cytosol-facing, N-terminally bound, endoplasmic reticulum (ER)-localized protein catalyzes the C8-hydroxylation of geraniol, requiring the transfer of two electrons catalyzed by the ER-bound, NADPH-dependent cytochrome P450 reductase (CPR)^8^. Optimization of heterologous CYP activity in yeast can be achieved by selecting or engineering for improved enzyme activity and by optimising CYP and CPR ratios or expression levels^9,10^.The complex relationship between CYP and CPR proteins is further demonstrated by the evolutionary conservation of their interaction domains, where CYP-CPR pairs from different kingdoms can functionally complement, alter or influence enzyme activity^10,11^. Although *S. cerevisiae* expresses a native CPR, heterologous co-expression of a plant CPR frequently results in higher CYP activity^12^.

The synthesis of nepetalactone in *S. cerevisiae* branches from the mevalonate pathway following geranyl pyrophosphate (GPP) dephosphorylation to geraniol by geraniol synthase (GES)^13^ (Figure 1). G8H and CPR hydroxylate geraniol to 8-hydroxygeraniol (8HG) followed by oxidation to 8-oxogeraniol, by 8HG oxidoreductase^14^. Iridoid synthase, an NADPH-dependant reductive cyclase, and three nepetalactol-related short-chain reductases (NEPS), subsequently cyclize and oxidize 8-oxogeraniol to nepetalactol and nepetalactone (Figure 1)^15^. In this current study, we report on the optimization of 8HG synthesis in *S. cerevisiae* while reducing off-target reactions in a geraniol producing *S. cerevisiae* strain, leading to *in vivo* production of the insect repellent nepetalactone. We demonstrate that non-cognate CYP-CPR combinations and the ratio of such combinations can modulate 8HG production for increased nepetalactone biosynthesis. Optimization of G8H activity not only leads to increased nepetalactone titers but is also vital in the optimization of an efficient strictosidine pathway in yeast.

**Figure 1.**
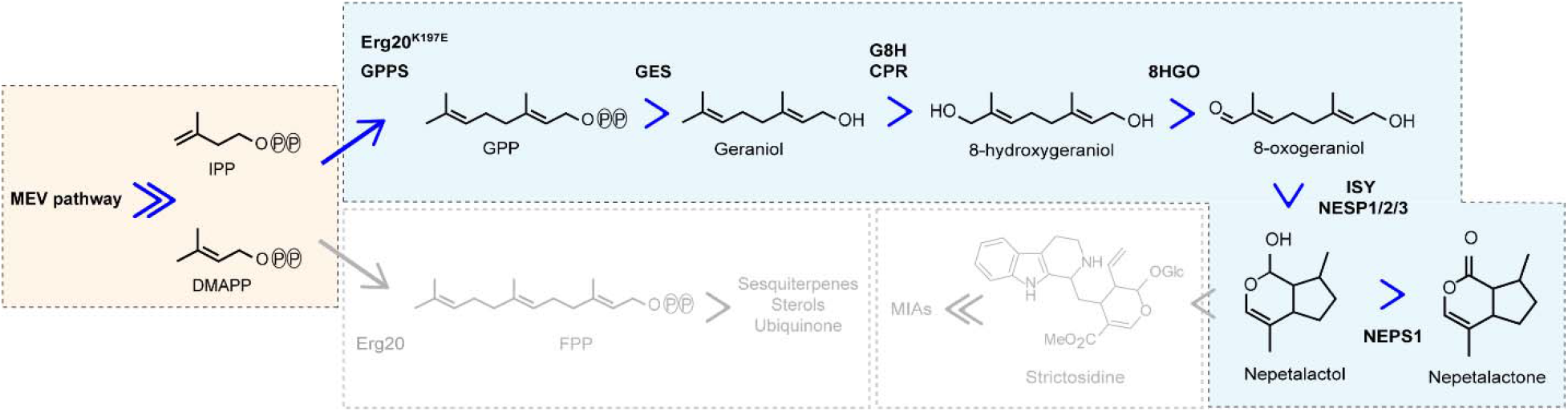
Overview of nepetalactone biosynthesis in engineered *S. cerevisiae*. Mevalonate pathway isopentenyl pyrophosphate (IPP) and dimethylallyl pyrophosphate (DMAPP) are condensed by farnesyl pyrophosphate synthase (ERG20) and geranyl pyrophosphate synthase (GPPS) to form geranyl pyrophosphate (GPP). Geraniol synthase (GES) then converts GPP to geraniol which is then hydroxylated to 8-hydroxygeraniol (8HG). The conversion of 8HG to nepetalactone occurs following oxidation by 8-hydroxygeraniol oxidase (8HGO), cyclisation by iridoid synthase (ISY) and a combination of three nepetalactol-related short-chain reductases (NEPS), and oxidation by NEPS1.

## RESULTS AND DISCUSSION

### *In vivo* synthesis of geraniol

*S. cerevisiae* GPP levels are not sufficient for production of detectable amounts of geraniol, hence the modification of the native mevalonate pathway and the expression of exogenous plant derived genes is essential. To increase GPP synthesis, an additional copy of two rate-limiting mevalonate pathway genes were integrated in the genome, a truncated *ScHMGR* and *SclDII*^16^ along with a mutated *ERG20 (ERG20^K197E^* ) expressed from its native locus^5^. The expression of a geraniol synthase (*ObGES*) and the subsequent removal of the N-terminal plastid targeting peptide^17^ resulted in accumulated titers of 7.22 ± 0.06 mg/L/OD_600_ of geraniol (Figure S1). The integration of a dedicated GPP synthase from *Abies grandis* (*AgGPPS*) further increased geraniol titers by over 2-fold, reaching 16.18 ± 0.03 mg/L/OD_600_. This geraniol-producing strain was used as the starting point for optimizing 8HG and nepetalactone synthesis.

### Screening of G8H/CPR combinations for optimized 8HG synthesis

Synthesis of 8HG in *S. cerevisiae* requires optimizing the rate-limiting hydroxylation of geraniol by G8H. Using the *Cr*G8H amino acid sequence as the search query, a preliminary list of predicted G8Hs was established based on sequence identity and functional domains (Figure 2A)^18–20^. This library was paired with a CPR collection (Figure 2B) similarly generated using the CPR amino acid sequence from *Papaver somniferum* as the search query (Table S1). The CYPs and CPRs from these libraries were paired combinatorially and analyzed for improved geraniol hydroxylation activity in yeast. The *Catharanthus roseus* G8H remained the most efficient enzyme for C8 geraniol hydroxylation (Figure 2C, Table S2). When paired with *Cr*G8H, several CPRs increased 8HG titers, with *Ns*CPR accumulating 7.36 ± 0.97 mg/L/OD_600_, a 3.3-fold increase in comparison to *Cr*CPR. Furthermore, *Pk*CPR and *Ns*CPR increased the molar conversion of geraniol to 8HG by 48.2 and 48.5%, respectively (Table S3). The range of monoterpene profiles across CYP-CPR pairs in Figure 2C-D may be explained by variations in coupling efficiencies, potentially influenced by the transmembrane domain and membrane lipid composition^21^. When paired with the *Cr*G8H, several non-cognate CPRs may have increased coupling efficiencies thereby increasing substrate turnover and 8HG titers.

**Figure 2.**
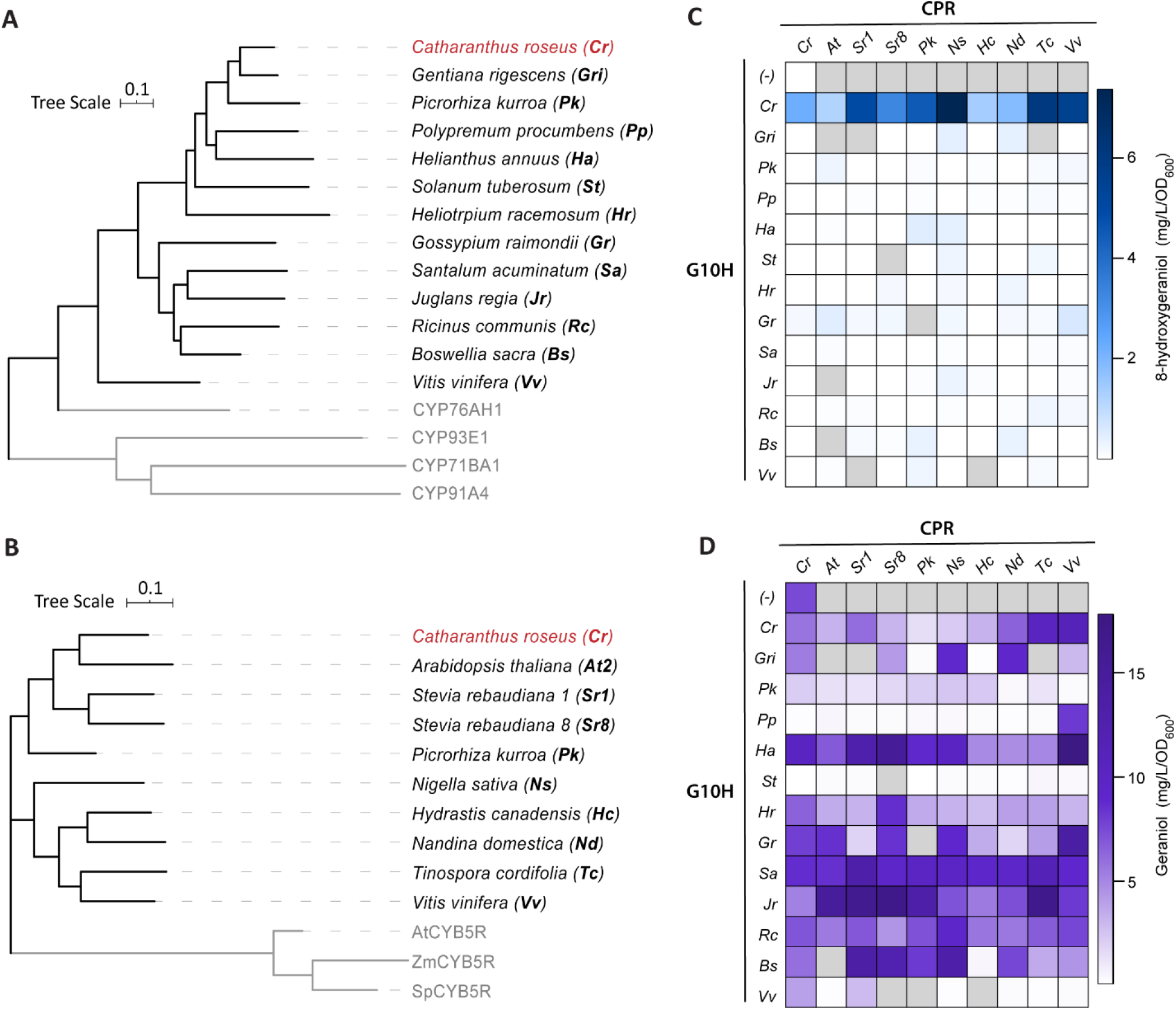
Analysis of CYP-CPR pairs on monoterpene production. **A.** Bootstrapped maximum likelihood phylogenetic tree of predicted G8Hs, rooted using outgroups (grey). **B.** Bootstrapped maximum likelihood phylogenetic tree of the CPR library rooted by cytochrome b5 reductase (CYB5R) outgroups (grey). **C.** 8HG and **D.** Geraniol synthesis from CYP-CPR combinations measured by GC-MS after 48 hours (Table S2 and S4). *n* = 3. Grey squares represent CYP-CPR pairs that could not be successfully constructed.

Seven out of the 122 CYP-CPR combinations tested showed >2-fold increases in geraniol titers (Figure 2D, Table S4). *Ha*G8H, for example, increased geraniol accumulation to 17.38 ± 1.80 mg/L/OD_600_ when paired with *Vv*CPR, with no observed 8HG accumulation, which could indicate geraniol synthase activity^22^. Analysis using the ExPASy ScanProsite tool^23^ identified the highly conserved terpene synthase Mg^2+^ binding motif (DDxx(D/E))^24^ in the *Ha*G8H variant, possibly explaining the increase in geraniol accumulation. When compared to the parent geraniol-producing strain, several CYP-CPR combinations resulted in decreased geraniol titers, as expected, but without a corresponding increase in 8HG (Figure 2C and 2D) potentially suggesting the conversion of geraniol to unidentified, non-productive products^25,26^. Nonetheless, the identification of several CPRs that improve CYP-catalyzed 8HG synthesis provides a baseline for further optimizations.

### Optimization of CrG8H-CPR expression ratio

As CYP:CPR ratios can influence substrate turnover and therefore metabolite accumulation^27^, we iteratively integrated multiple copies of *CrG8H* into a strain with *CrCPR* to determine the optimal ratio for 8HG production. As the CYP:CPR ratio was increased so were 8HG levels, up to 10.46 ± 2.27 mg/L/OD_600_ with three copies of *CrG8H*, an 88% molar conversion of geraniol (Figure 3A, Table S5). Addition of a single *CrG8H* copy increased cell growth, potentially by reducing levels of geraniol, as it is known to permeabilize the cell membrane^28^. Yet, expressing two or three *CrG8H* copies reduced growth, which may have been a result of ER disruption by the overexpression of heterologous membrane-bound proteins (Figure 3A)^29^.

**Figure 3.**
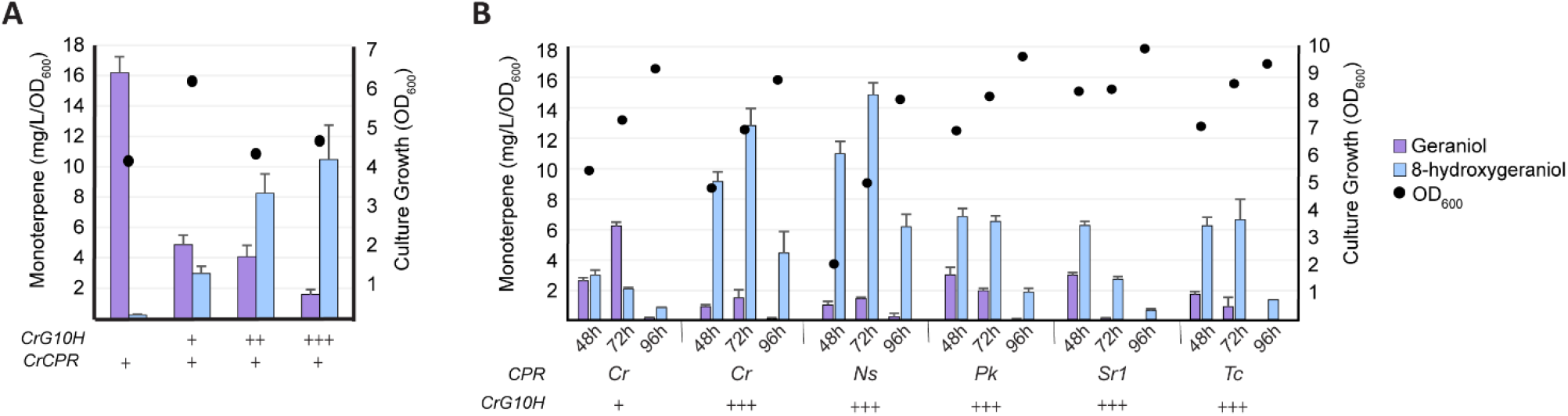
Monoterpene production from CYP-CPR optimization. **A.** Monoterpene concentrations and cell growth measured by GC-MS after 48 hours in a geraniol-producing strain expressing a single copy of *Cr*CPR with one, two or three copies of *Cr*G8H. *n* = 3. **B.** Monoterpene production and growth of geraniol-producing strains expressing three copies of *Cr*G8H together with various CPRs measured by GC-MS after 48, 72 and 96 hours. *n* = 3. Error bars indicate standard deviation.

We next tested the translatability of our higher 8HG titers, resulting from a 3:1 *CrG8H* to *CrCPR* ratio, using four of the CPRs that were identified to increase 8HG production or molar conversion (Figure 2C). When expressed along with 3x-*CrG8H*,*Ns*CPR accumulated a slightly higher 8HG titer (p>0.05) of 14.75 ± 0.77 mg/L/OD_600_ when compared to 12.71 ± 1.11 mg/L/OD_600_ for *Cr*CPR (Figure 3B). However, this trend was not consistent with the remaining three CPRs, reinforcing the requirement for optimal CYP:CPR ratios for enzyme pairs, as several CPRs outperformed *Cr*CPR in the 1x-*Cr*G8H strain but not in the 3x-*CrG8H* strain. *Ns*CPR could be a potential candidate for improving 8HG biosynthesis, however the low growth observed at both 48 and 72h influences total accumulation of 8HG and therefore requires further optimisation for its use as an alternative to *Cr*CPR. High-throughput combinatorial multiplexed integration of multiple G8H and CPR could serve as a tool to optimise CYP:CPR ratios, as previously described for butanediol synthesis in *S. cerevisiae* from xylose^30^.

### Eliminating non-productive pathway intermediates

Optimal 8HG production in *S. cerevisiae* requires mitigating off-target products that sequester carbon away from its biosynthesis. Increasing 8HG titers correlated with an increase in an off-target metabolite predicted to be isopulegol by the NIST08 MS database (Figure 4A, peak 3)^31^. Subsequent comparison of the shunt product spectra to that of an authentic isopulegol standard revealed different retention times and mass spectra between isopulegol and the unidentified peak (Figure S2) indicating that the shunt product was not isopulegol. Further experimentation, including NMR, would be needed to determine the identity of this product. To mitigate shunt product formation, the yeast NADPH oxidoreductases, old yellow enzymes Oye2 and Oye3, were deleted from the *Cr*G8H expressing strain^32,33^. Deletion of *OYE3* increased both 8HG and the shunt product peak by 2- and 4- fold, respectively (Figure 4B), suggesting Oye3 catalyzes the synthesis of non-productive terpene metabolites, but not the identified shunt product. Deletion of both *OYE2* and *OYE3* eliminated the production of the shunt product resulting in 3- and 8.5-fold increase in 8HG in the strains expressing one or three copies of *Cr*G8H, respectively.

**Figure 4.**
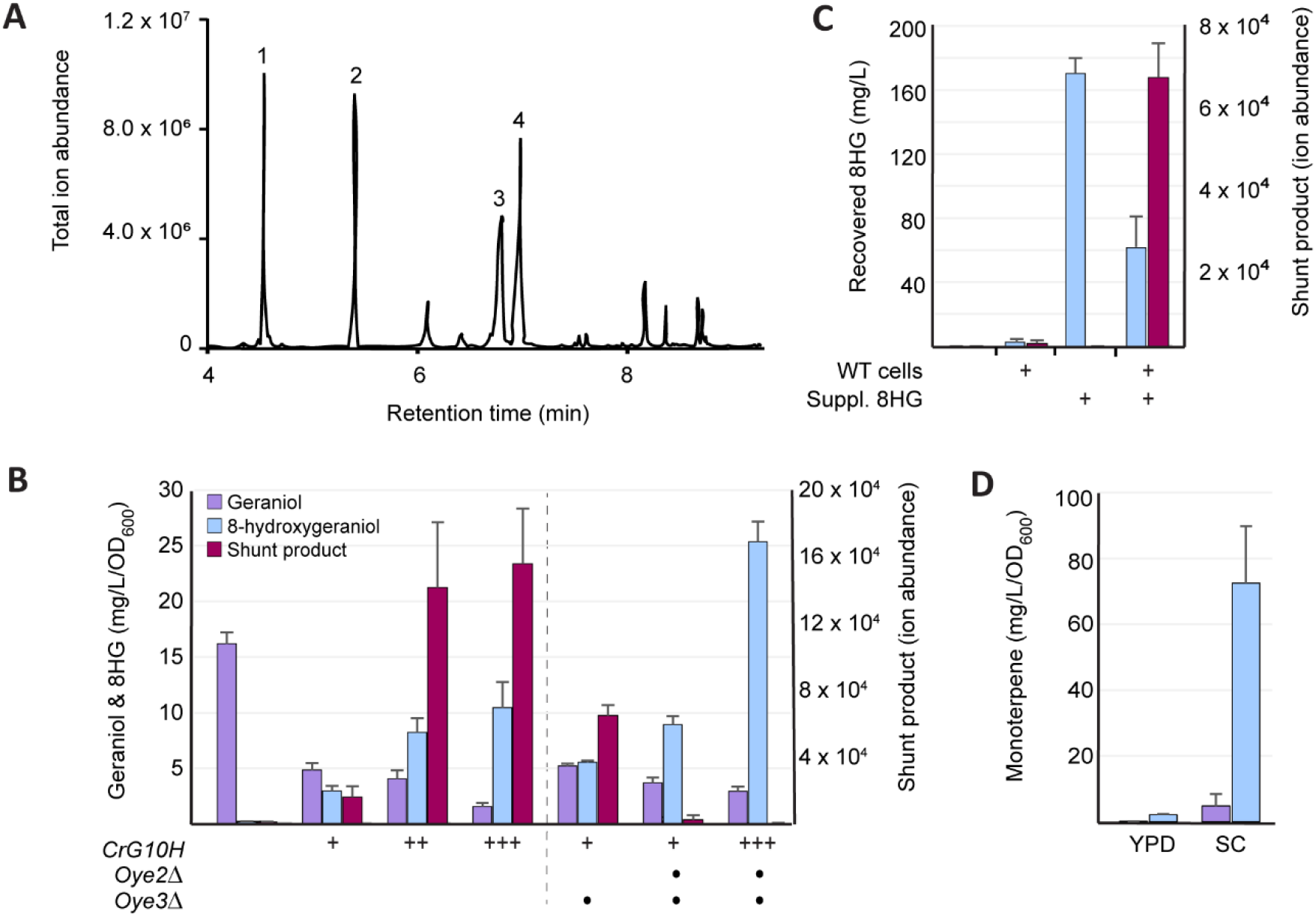
Elimination of unknown monoterpene shunt product. **A.** GC-MS total ion abundance chromatograms for extracts of the 3x-*Cr*G8H strain culture broths. Peak identification using analytic standards are as follows: **1.** Geraniol; **2.** Eugenol (internal standard); **3.** Unknown **4.** 8HG. Remaining peaks are yeast metabolites present in the wild-type control. **B.** Monoterpene titers in the *CrG8H* background strains with *OYE2* and *OYE3* deletions. Due to the lack of an appropriate standard, the shunt product is reported as area under the curve. *n = 3*. **C.** Monoterpene titers following supplementation of 8HG to wild-type *S. cerevisiae* culture broth. *n = 3*. **D.** Monoterpene titers from the strain expressing 3x-*Cr*G8H grown in YPD or SC media. *n = 4*. Error bars indicate standard deviation.

To test if 8HG is the precursor in the synthesis of the shunt product, cultures of wild-type *S. cerevisiae* were supplemented with 8HG and analyzed for the appearance of the off-target peak (Figure 4C). The observed accumulation of the shunt product suggests 8HG is the precursor metabolite and that the reaction(s) required for its conversion is catalyzed by a native yeast enzyme(s). In addition, this off-target peak was only observed from cultures in which the geraniol-producing strain was expressing the *Cr*G8H (Figure S3), providing further evidence for 8HG as the precursor.

Following the mitigation of non-productive reactions, we tested whether changes to media composition could increase cell productivity. The strain expressing three copies of *Cr*G8H was grown in YPD, 1xSC, 2xSC and 2xSC with 2% glucose and 8HG accumulation was assessed. When grown in 1xSC, 8HG titers increased by 6.9-fold when compared to YPD, reaching 72.52 ± 17.27 mg/L/OD_600_ (Figure 4D), the remaining media compositions did not affect titers (data not shown). The difference in 8HG accumulation between YPD and SC may be due to the presence of yeast extract in YPD, potentially providing end-product metabolites that could regulate biochemical pathways. With the mitigation of non-productive reductions and the identification of a more optimal growth media for increasing productivity, the remainder of the pathway for nepetalactone biosynthesis was integrated into this 8HG-producing strain.

### Production of nepetalactone

Several additional enzymes are required to achieve *de novo* biosynthesis of nepetalactone in *S. cerevisiae*. First, 8HG is oxidized by 8HGO to 8-oxogeraniol, which is subsequently cyclized by ISY, to produce nepetalactol (Figure 5A). In the *Nepeta* plant species, ISY works with three NEPS for the cyclisation of 8-oxogeraniol to nepetalactol, determining the stereochemistry of the bridgehead 4a-7a-carbons^34^ (Figure 5A). ISY catalyses the initial reduction of 8-oxogeraniol to the activated oxocitronellyl enol intermediate enabling spontaneous cyclisation to (4aS,7S,7aR)-nepetalactol. NEPS1 and NEPS2 stabilize the enol intermediate thereby reducing off-target iridodial products^34^. Subsequent oxidation by NEPS1 is responsible for the conversion of nepetalactol to their cognate nepetalactone isomers^34^. The genes for these remaining enzymes were sequentially integrated into the strain harboring 3x-*Cr*G8H. The *OYE2* and *OYE3* genes were then iteratively deleted from each strain to assess their effect on nepetalactone synthesis (Figure 5B).

**Figure 5.**
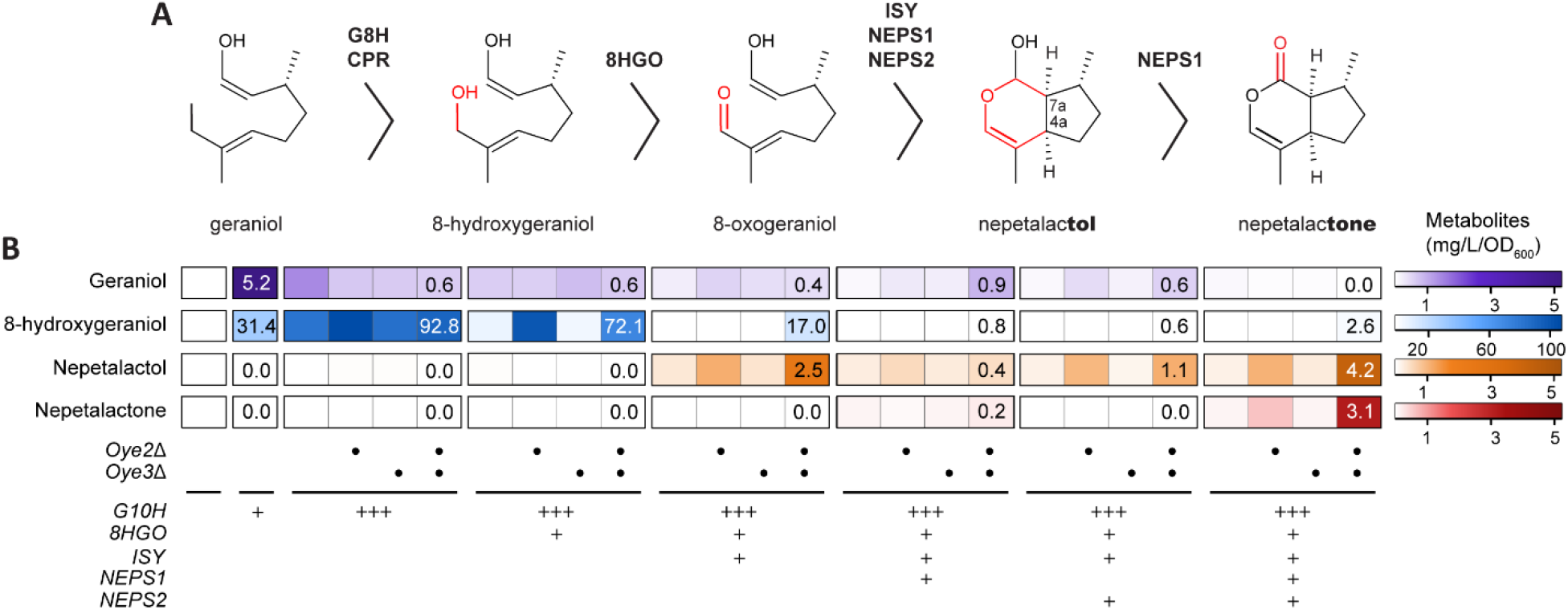
*De novo* synthesis of nepetalactone in *S. cerevisiae*. **A.** Enzymatic conversion of geraniol to nepetalactone. **B.** Monoterpene and iridoid synthesis from glucose following the integration of *Cr8HGO*, *ISY*, *NEPS1*, *NEPS2* and deletion of *OYE2* and *OYE3*. Titers were analyzed by Q-TOF LC-MS after 72 hours. *n = 4*.

Following the introduction of *Cr8HGO* and *ISY* into the 3x-*CrG8H* strain, nepetalactol accumulation reached 0.33 ± 0.05 mg/L/OD_600_ with titers increasing to 2.54 ± 0.06 mg/L/OD_600_ following the deletion of both *OYE2* and *OYE3* (Figure 5B). Subsequent integration of *NEPS1*, resulted in trace amounts of nepetalactone (0.19 ± 0.01 mg/L/OD_600_) while the integration of *NEPS2* alone, following *ISY* integration, did not produce any measurable nepetalactone (Figure 5B). Since NEPS2 is not directly involved in nepetalactone synthesis, this result is in agreement with its reported activity^15^. However, nepetalactol titers decreased to 1.11 ± 0.1 mg/L/OD_600_ upon *NEPS2* integration compared to the strain with only *ISY* (Figure 5C). This may suggest that without *NEPS1*, some reactive enol intermediate is lost to non-productive reactions. The integration of both *NEPS1* and *NEPS2* into the *ISY* strain resulted in the highest titers of both nepetalactol and nepetalactone, accumulating 4.22 ± 0.79 mg/L/OD_600_ and 3.19 ± 0.66 mg/L/OD_600_, respectively. This is a notable increase in titers compared to the same strain with *OYE2* and *OYE3*, where accumulation of nepetalactol and nepetalactone only reached 0.19 ± 0.02 mg/L/OD_600_ and 0.12 ± 0.01 mg/L/OD_600_, respectively. These results suggest that both NEPS1 and NEPS2 are required for efficient nepetalactone production and deletion of the *OYE* homologs are necessary for increasing titers.

This system presents the first *de novo* nepetalactone production in *S. cerevisiae* and lays the foundation for future optimization strategies aimed at increasing total biosynthesis of nepetalactone from glucose, thereby providing an opportunity to produce a plant-derived alternative insect repellent at industrial scale.

## MATERIALS AND METHODS

### Strains, media and DNA manipulation

*S. cerevisiae* strains used in this study are listed in Table S6. Yeast cultures were grown in YPD (10 g/L yeast extract, 20 g/L tryptone, 20 g/L dextrose) or synthetic complete (SC) medium (6.8 g/L Yeast Nitrogen Base (YNB), amino acids, 2% (w/v) glucose) and supplemented with 100μg/ml geneticin (G418) to select for the ERG20^K197E^ background. When appropriate, 200 μg/ml of hygromycin was added for the selection of the pCas plasmid. Plasmids were maintained and propagated in *Escherichia coli* DH5α. *E. coli* cultures were incubated at 37°C with shaking at 200 rpm in Lysogeny Broth (LB) supplemented with either 100 μg/mL of ampicillin or 100 μg/mL of hygromycin.

Genes and primers used in this study are described in Tables S1 and S7, respectively. Promoters, terminators, and the yeast mevalonate pathway genes were amplified by PCR from *S. cerevisiae* CEN.PK genomic DNA. Genes originating from plant species, apart from the G8H variant library, were amplified from pYES2-GerOH^5^. The pCas plasmid was purchased from (Addgene plasmid #60847) and modified to express the hphNT1 cassette to confer hygromycin resistance. Genes encoding G8H enzymes were codon optimized and synthesized by GeneArt or Twist Bioscience. The *G8H* genes were further sub-cloned individually into pJet2.1 using the CloneJET PCR cloning kit. The integrative plasmids used to introduce the genes for nepetalactone synthesis - *8HGO*, *ISY*, *NEPS1* and *NEPS2* - were constructed using Golden Gate assembly^35^ with components from the Yeast Toolkit^36^. Plasmids were purified from *E. coli* stocks using the GeneJET plasmid miniprep kit where the DNA parts were amplified by PCR using Phusion High-Fidelity DNA polymerase, resolved using 0.8% (w/v) agarose gel electrophoresis and individually purified using Qiagen Gel Purification kit.

### Library construction and phylogenetic analysis

The G8H sequence library was assembled using the *Cr*G8H amino acid sequence as the search query for three separate databases: Plant Metabolic Network^18^, 1k Plant Collection^20^ and NCBI using DeltaBLAST^19^. From each database, sequences with over 65% homology were selected. This list was consolidated using CD-HIT^37^ with a sequence identity cut-off of 90%. Bootstrapped (500 replicates) maximum likelihood phylogenetic trees for the G8H and CPR libraries were built following MUSCLE alignment using MEGA7^38^. The phylogenetic trees were rooted using the various outgroups; amino acid sequences can be found in Table S1.

### Yeast transformation and CRISPR/Cas9-mediated genome integration

*S. cerevisiae* was transformed using a method modified from the Gietz PEG/LiAc protocol^39^. Strains were initially grown overnight in YPD G418 medium to maintain selection of the *Erg20^K197E^* mutant. After cells were harvested and washed, the cell pellet was suspended in 3M lithium acetate (5.6 μL per transformation) with DNA and water to a total volume of 40 μL. The competent cells were incubated at room temperature for 10 minutes before adding the transformation mix and completing the rest of the protocol. All genomic integrations (Table S8) were performed using the CRISPR-Cas9 delivery system^40^. Between 1 and 3 μg of donor DNA were used for CRISPR Cas9-mediated integration.

### Analysis of monoterpenes and iridoids

Monoterpenes were analyzed by either GC-MS or Q-TOF LC-MS. Strains were inoculated in a 96-well deep well plate containing 500 μL of YPD or SC with G418 and incubated at 30°C and 200 rpm for either 48, 72 or 96 hours. For GC-MS sample preparation, 400 μL culture aliquots were extracted into 200 μL of ethyl acetate containing eugenol as an internal standard. Extracts were analyzed on an Agilent 6890N GC coupled to an Agilent 5875C mass selective detector. An HP-5ms column (30 m ×0.25 mm × 0.25 μm film thickness) was used with a 1 μL splitless injection, hydrogen gas as the carrier with a constant flow of 1.3 mL/min and an inlet temperature of 240°C. The initial oven temperature was set to 60°C for 2 minutes, followed by an increase to 150°C at 30°C/min to, 10°C/min to 220°C and 30°C/min to 325°C (5 min hold). Total ion chromatograms were recorded between *m/z* 50–220. For LC-MS, 400 μL culture aliquots were extracted into 400 μL of 100% methanol. Samples were analyzed using the 1290 Infinity II LC system coupled to a 6560 Ion Mobility Q-TOF (Agilent Technologies). Samples were analyzed with a Zorbax Eclipse Plus C18 50 × 4.6 mm column (Agilent Technologies) using a gradient of solvent A (100% water, 0.1% formic acid) and solvent B (100% acetonitrile, 0.1% formic acid) to separate metabolites. Metabolites were separated using a linear gradient: 0-5 min 95% A/5% B, 5- 6 min 40% A/60% B at a flow rate of 0.3 mL/min, 6-7 min 5% A/95% B at a flow rate of 0.45 mL/min followed by a 2 min equilibration at 95% A/5% B at a flow rate of 0.4 mL/min. The system was operated in positive electrospray mode using a capillary voltage 4000 V, fragmentor voltage 400 V, source temperature 325 °C, nebulizer pressure 55 psi and, gas flow 10 L/ min. Citronellol, geraniol, eugenol (Sigma Aldrich), 8HG, nepetalactol and nepetalactone (Toronto Research Chemicals Ins.) standards were used to determine retention times and calibration curves for comparison with samples (Figure S4). Isopulegol was identified using the NIST 08 standard reference database^31^.

## ACKNOWLEDGEMENTS

We thank Yves Gelinas for the use of his GC-MS. This study was financially supported by a NSERC Discovery grant. M.E.D. was supported by the Faculty of Arts and Science Graduate fellowship from Concordia University and D.T. was supported by the SynBioApps NSERC-CREATE training program and V.J.J.M. is supported by a Concordia University Research Chair.

## AUTHOR CONTRIBUTIONS

M.E.D and V.J.J.M. designed the research. M.E.D., and D.T. performed the experiments. V.J.J.M supervised the research. M.E.D. wrote the manuscript with editing help from D.T. and V.J.J.M.

## ADDITIONAL INFORMATION

Supplementary information is available in the online version of the paper. Correspondence and requests for materials should be addressed to V.J.J.M.

## DATA AVAILABILITY

The data that support the findings of this study are available from the authors upon reasonable request.

