## Supporting Information for "Engineering yeast for *de-novo* synthesis of the insect repellent - nepetalactone"

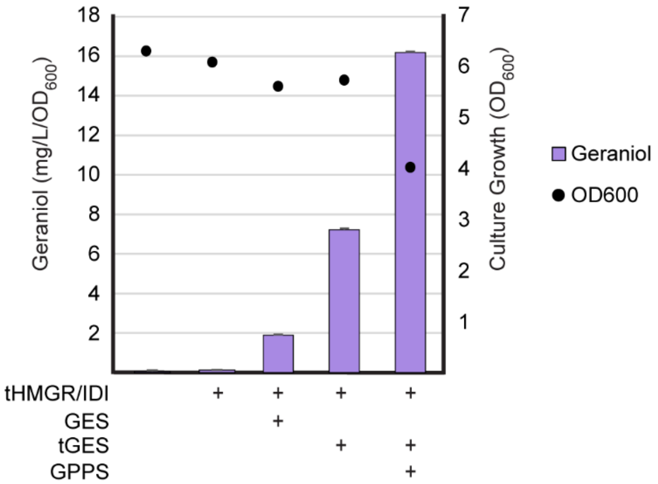


**Figure S1. Increasing endogenous titers of geraniol.** Geraniol accumulation in strains with genomic integrations aimed at increasing geraniol titers. Geraniol (purple) concentrations and $OD_{600}$ values were measures after 48 hours of growth. Error bars represent the standard deviation from triplicate cultures.

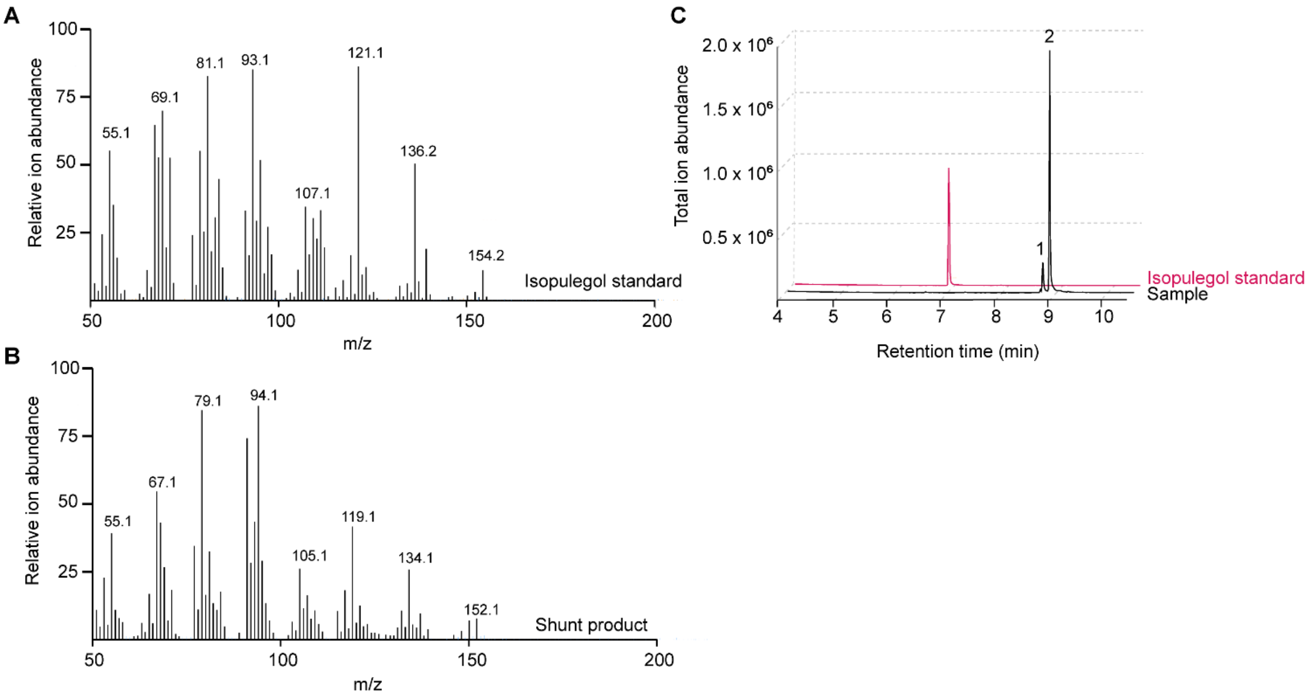


**Figure S2. Comparison of isopulegol and 'shunt product' mass spectra and chromatograms from GC-MS.** The relative ion abundance for A) isopulegol and B) the 'shunt product'. C) Total ion abundance chromatograms for the isopulegol standard (pink) and extract from the wild-type strain supplemented with 8HG. Peak identification as follows: **1.** 'shunt product' and **2.** 8HG.

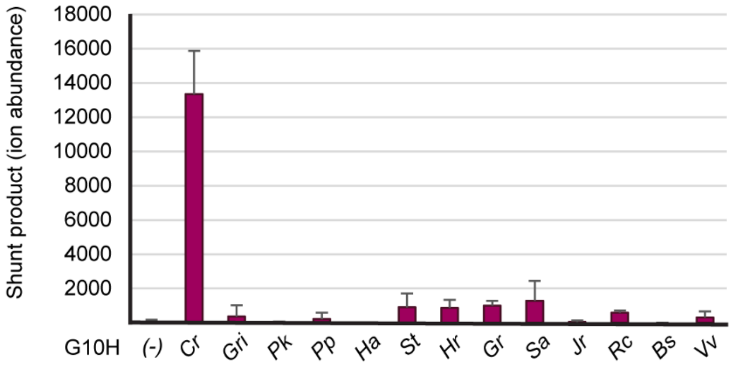


**Figure S3. Analysis of shunt product accumulation in strains expressing G10H variants.** Shunt product (maroon) accumulation was measures after 48 hours of growth. Error bars represent the standard deviation triplicate cultures.

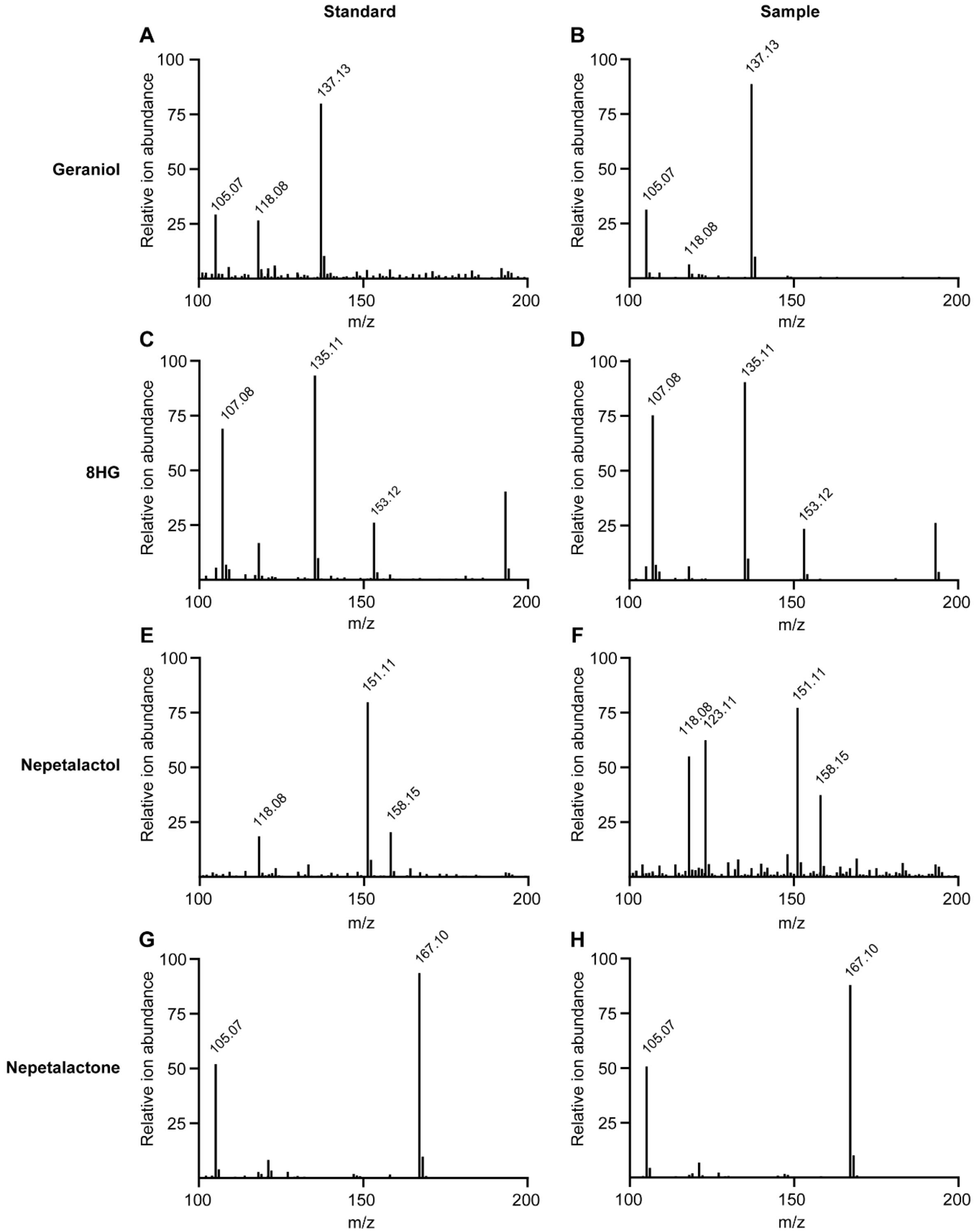


**Figure S4. Comparison of mass spectra from standards and samples from QTOF-LC-MS.** The mass spectra from the standards and samples for A-B) geraniol, C-D) 8HG, E-F) nepetalactol and G-H) nepetalactone were compared.

### Table S1. Codon-optimized genes and synthetic sequences employed in this study

| Gene/Insert | Sequence (5' > 3') |
|---|---|
| *SctHMGR* | ATGGTTTTAACCAATAAAACAGTCATTTCTGGATCGAAAGTCAAAAGTTTATCATCTGCGCAATCGAGCTCATCAGGACCTTCAT<br>CATCTAGTGAGGAAGATGATTCCCGCGATATTGAAAGCTTGGATAAGAAAATACGTCCTTTAGAAGAATTAGAAGCATTATTAA<br>GTAGTGGAAATACAAAACAATTGAAGAACAAAGAGGTCGCTGCCTTGGTTATTCACGGTAAGTTACCTTTGTACGCTTTGGAGA<br>AAAAATTAGGTGATACTACGAGAGCGGTTGCGGTACGTAGGAAGGCTCTTTCAATTTTGGCAGAAGCTCCTGTATTAGCATCTGA<br>TCGTTTACCATATAAAAATTATGACTACGACCGCGTATTTGGCGCTTGTTGTGAAAATGTTATAGGTTACATGCCTTTGCCCGTTG<br>GTGTTATAGGCCCCTTGGTTATCGATGGTACATCTTATCATATACCAATGGCAACTACAGAGGGTTGTTTGGTAGCTTCTGCCATG<br>CGTGGCTGTAAGGCAATCAATGCTGGCGGTGGTGCAACAACTGTTTTAACTAAGGATGGTATGACAAGAGGCCCAGTAGTCCGT<br>TTCCCAACTTTGAAAAGATCTGGTGCCTGTAAGATATGGTTAGACTCAGAAGAGGGACAAAACGCAATTAAAAAAGCTTTTAAC<br>TCTACATCAAGATTTGCACGTCTGCAACATATTCAAACTTGTCTAGCAGGAGATTTACTCTTCATGAGATTTAGAACAACTACTG<br>GTGACGCAATGGGTATGAATATGATTTCTAAAGGTGTCGAATACTCATTAAAGCAAATGGTAGAAGAGTATGGCTGGGAAGATA<br>TGGAGGTTGTCTCCGTTTCTGGTAACTACTGTACCGACAAAAAACCAGCTGCCATCAACTGGATCGAAGGTCGTGGTAAGAGTGT<br>CGTCGCAGAAGCTACTATTCCTGGTGATGTTGTCAGAAAAGTGTTAAAAAGTGATGTTTCCGCATTGGTTGAGTTGAACATTGCT<br>AAGAATTTGGTTGGATCTGCAATGGCTGGGTCTGTTGGTGGATTTAACGCACATGCAGCTAATTTAGTGACAGCTGTTTTCTTGG<br>CATTAGGACAAGATCCTGCACAAAATGTTGAAAGTTCCAACTGTATAACATTGATGAAAGAAGTGGACGGTGATTTGAGAATTT<br>CCGTATCCATGCCATCCATCGAAGTAGGTACCATCGGTGGTGGTACTGTTCTAGAACCACAAGGTGCCATGTTGGACTTATTAGG<br>TGTAAGAGGCCCGCATGCTACCGCTCCTGGTACCAACGCACGTCAATTAGCAAGAATAGTTGCCTGTGCCGTCTTGGCAGGTGA<br>ATTATCCTTATGTGCTGCCCTAGCAGCCGGCCATTTGGTTCAAAGTCATATGACCCACAACAGGAAACCTGCTGAACCAACAAAA<br>CCTAACAATTTGGACGCCACTGATATAAATCGTTTGAAAGATGGGTCCGTCACCTGCATTAAATCCTAA |
| *ScIDI* | ATGACTGCCGACAACAATAGTATGCCCCATGGTGCAGTATCTAGTTACGCCAAATTAGTGCAAAACCAAACACCTGAAGACATT<br>TTGGAAGAGTTTCCTGAAATTATTCCATTACAACAAAGACCTAATACCCGATCTAGTGAGACGTCAAATGACGAAAGCGGAGAA<br>ACATGTTTTTCTGGTCATGATGAGGAGCAAATTAAGTTAATGAATGAAAATTGTATTGTTTTGGATTGGGACGATAATGCTATTG<br>GTGCCGGTACCAAGAAAGTTTGTCATTTAATGGAAAATATTGAAAAGGGTTTACTACATCGTGCATTCTCCGTCTTTATTTTCAAT<br>GAACAAGGTGAATTACTTTTACAACAAAGAGCCACTGAAAAAATAACTTTCCCTGATCTTTGGACTAACACATGCTGCTCTCATC<br>CACTATGTATTGATGACGAATTAGGTTTGAAGGGTAAGCTAGACGATAAGATTAAGGGCGCTATTACTGCGGCGGTGAGAAAAC<br>TAGATCATGAATTAGGTATTCCAGAAGATGAAACTAAGACAAGGGGTAAGTTTCACTTTTTAAACAGAATCCATTACATGGCAC<br>CAAGCAATGAACCATGGGGTGAACATGAAATTGATTACATCCTATTTTATAAGATCAACGCTAAAGAAAACTTGACTGTCAACC<br>CAAACGTCAATGAAGTTAGAGACTTCAAATGGGTTTCACCAAATGATTTGAAAACTATGTTTGCTGACCCAAGTTACAAGTTTAC<br>GCCTTGGTTTAAGATTATTTGCGAGAATTACTTATTCAACTGGTGGGAGCAATTAGATGACCTTTCTGAAGTGGAAAATGACAGG<br>CAAATTCATAGAATGCTATAA |
| *ObGES* | ATGTCTTGTGCACGGATCACCGTAACATTGCCGTATCGCTCCGCAAAAACATCAATTCAACGGGGAATTACGCATTACCCCGCCC<br>TTATACGCCCACGCTTCTCTGCTTGCACGCCTTTGGCATCGGCGATGCCTCTAAGTTCAACTCCTCTCATCAACGGGGATAACTCT<br>CAGCGTAAAAACACACGTCAACACATGGAGGAGAGCAGCAGCAAGAGGAGAGAATATCTGCTGGAGGAAACGACGCGAAAAC |

| | |
|---|---|
| | TGCAGAGAAACGACACCGAATCGGTGGAGAAACTCAAGCTTATCGACAACATCCAACAGTTGGGAATCGGCTACTATTTTGAGG<br>ACGCCATCAACGCCGTACTCCGCTCGCCTTTCTCCACCGGAGAAGAAGACCTCTTCACCGCTGCTCTGCGCTTCCGCTTGCTCCGC<br>CACAACGGCATCGAAATCAGCCCTGAAATATTCCTAAAATTCAAGGACGAGAGGGGAAAATTCGACGAATCGGACACGCTAGG<br>GTTACTGAGCTTGTACGAAGCGTCAAATTTGGGGGTTGCAGGAGAAGAAATATTGGAGGAGGCTATGGAGTTTGCGGAGGCTCG<br>CCTGAGACGGTCGCTGTCAGAGCCGGCGGCGCCGCTTCATGGTGAGGTGGCGCAAGCGCTAGATGTGCCGAGGCATCTGAGAAT<br>GGCGAGGTTGGAAGCGAGACGATTCATCGAGCAGTATGGTAAACAGAGCGATCATGATGGAGATCTTTTGGAGCTGGCAATTTT<br>GGATTATAATCAAGTTCAGGCTCAACACCAATCCGAACTCACTGAAATAATCAGGTGGTGGAAGGAGCTCGGTTTGGTGGATAA<br>GTTGAGTTTTGGGCGAGACAGACCATTGGAGTGCTTTTTGTGGACCGTGGGGCTCCTCCCAGAGCCCAAGTATTCGAGCGTTAGA<br>ATAGAGTTGGCGAAAGCCATCTCTATTCTCTTAGTGATCGATGATATTTTCGATACCTATGGAGAGATGGATGACCTCATCCTCTT<br>CACCGATGCAATTCGAAGATGGGATCTTGAAGCAATGGAGGGGCTCCCTGAGTACATGAAAATATGCTACATGGCGTTGTACAA<br>TACCACCAATGAAGTATGCTACAAAGTGCTCAGGGATACTGGACGGATTGTCCTCCTTAACCTCAAATCTACGTGGATAGACATG<br>ATTGAAGGTTTCATGGAGGAAGCAAAATGGTTCAATGGTGGAAGTGCACCAAAATTGGAAGAGTATATAGAGAATGGAGTGTC<br>CACGGCAGGAGCATACATGGCTTTTGCACACATCTTCTTTCTCATAGGAGAAGGTGTTACACACCAAAATTCCCAACTCTTCACC<br>CAAAAACCCTACCCCAAGGTCTTCTCCGCCGCCGGCCGCATTCTTCGCCTCTGGGATGATCTCGGAACCGCCAAGGAAGAGCAA<br>GAGCGAGGAGATCTGGCTTCGTGCGTGCAGTTATTTATGAAAGAGAAGTCGTTGACGGAAGAGGAGGCAAGAAGTCGCATTTTG<br>GAAGAGATAAAAGGATTATGGAGGGATCTGAATGGGGAACTGGTCTACAACAAGAATTTGCCGTTATCCATAATCAAAGTCGCA<br>CTTAACATGGCGAGAGCTTCTCAAGTTGTGTACAAGCACGATCAAGACACTTATTTTTCAAGCGTAGACAATTATGTGGATGCCC<br>TCTTCTTCACTCAATAA |
| *t63OBGES* | ATGCAACACATGGAGGAGAGCAGCAGCAAGAGGAGAGAATATCTGCTGGAGGAAACGACGCGAAAACTGCAGAGAAACGACA<br>CCGAATCGGTGGAGAAACTCAAGCTTATCGACAACATCCAACAGTTGGGAATCGGCTACTATTTTGAGGACGCCATCAACGCCG<br>TACTCCGCTCGCCTTTCTCCACCGGAGAAGAAGACCTCTTCACCGCTGCTCTGCGCTTCCGCTTGCTCCGCCACAACGGCATCGA<br>AATCAGCCCTGAAATATTCCTAAAATTCAAGGACGAGAGGGGAAAATTCGACGAATCGGACACGCTAGGGTTACTGAGCTTGTA<br>CGAAGCGTCAAATTTGGGGGTTGCAGGAGAAGAAATATTGGAGGAGGCTATGGAGTTTGCGGAGGCTCGCCTGAGACGGTCGCT<br>GTCAGAGCCGGCGGCGCCGCTTCATGGTGAGGTGGCGCAAGCGCTAGATGTGCCGAGGCATCTGAGAATGGCGAGGTTGGAAG<br>CGAGACGATTCATCGAGCAGTATGGTAAACAGAGCGATCATGATGGAGATCTTTTGGAGCTGGCAATTTTGGATTATAATCAAG<br>TTCAGGCTCAACACCAATCCGAACTCACTGAAATAATCAGGTGGTGGAAGGAGCTCGGTTTGGTGGATAAGTTGAGTTTTGGGC<br>GAGACAGACCATTGGAGTGCTTTTTGTGGACCGTGGGGCTCCTCCCAGAGCCCAAGTATTCGAGCGTTAGAATAGAGTTGGCGA<br>AAGCCATCTCTATTCTCTTAGTGATCGATGATATTTTCGATACCTATGGAGAGATGGATGACCTCATCCTCTTCACCGATGCAATT<br>CGAAGATGGGATCTTGAAGCAATGGAGGGGCTCCCTGAGTACATGAAAATATGCTACATGGCGTTGTACAATACCACCAATGAA<br>GTATGCTACAAAGTGCTCAGGGATACTGGACGGATTGTCCTCCTTAACCTCAAATCTACGTGGATAGACATGATTGAAGGTTTCA<br>TGGAGGAAGCAAAATGGTTCAATGGTGGAAGTGCACCAAAATTGGAAGAGTATATAGAGAATGGAGTGTCCACGGCAGGAGCA<br>TACATGGCTTTTGCACACATCTTCTTTCTCATAGGAGAAGGTGTTACACACCAAAATTCCCAACTCTTCACCCAAAAACCCTACC<br>CCAAGGTCTTCTCCGCCGCCGGCCGCATTCTTCGCCTCTGGGATGATCTCGGAACCGCCAAGGAAGAGCAAGAGCGAGGAGATC<br>TGGCTTCGTGCGTGCAGTTATTTATGAAAGAGAAGTCGTTGACGGAAGAGGAGGCAAGAAGTCGCATTTTGGAAGAGATAAAAG<br>GATTATGGAGGGATCTGAATGGGGAACTGGTCTACAACAAGAATTTGCCGTTATCCATAATCAAAGTCGCACTTAACATGGCGA<br>GAGCTTCTCAAGTTGTGTACAAGCACGATCAAGACACTTATTTTTCAAGCGTAGACAATTATGTGGATGCCCTCTTCTTCACTCA<br>ATAA |

| | |
|---|---|
| *AgGPPS* | ATGGCTTACTCTGCTATGGCTACTATGGGCTATAACGGGATGGCAGCATCATGTCACACATTACATCCTACTTCACCACTTAAAC<br>CATTCCATGGCGCTTCTACAAGCCTAGAGGCATTCAATGGTGAGCACATGGGTCTGTTGAGAGGATACTCTAAGAGAAAGCTAT<br>CTTCATACAAAAACCCAGCTTCTCGTTCATCAAATGCTACCGTCGCGCAGTTACTGAACCCACCACAAAAGGGCAAAAAGGCAG<br>TAGAATTTGATTTCAACAAATACATGGACTCTAAAGCTATGACAGTTAATGAGGCATTGAACAAGGCGATTCCACTTAGGTACCC<br>ACAAAAGATATACGAATCTATGAGATACTCTCTGCTTGCCGGTGGAAAGAGAGTTAGACCTGTTTTGTGCATAGCAGCCTGCGA<br>ATTGGTAGGAGGTACCGAGGAATTGGCAATTCCAACAGCATGTGCAATAGAAATGATTCATACAATGTCACTTATGCACGATGA<br>TTTGCCTTGCATCGATAATGACGATCTAAGACGAGGAAAGCCTACGAATCATAAGATCTTTGGTGAAGATACTGCTGTTACTGCG<br>GGAAACGCTCTTCACAGTTACGCCTTCGAACATATCGCAGTATCAACATCCAAAACCGTGGGTGCTGATAGAATACTACGTATG<br>GTTTCTGAATTAGGCAGAGCCACAGGCTCCGAGGGAGTTATGGGTGGGCAGATGGTCGATATTGCCAGCGAGGGGGACCCTTCA<br>ATAGACTTACAAACTCTTGAATGGATTCATATTCATAAGACGGCAATGTTATTGGAGTGTTCTGTTGTCTGTGGTGCTATCATCGG<br>TGGTGCCAGTGAAATCGTAATCGAAAGAGCCCGTAGATACGCGAGGTGTGTGGGGCTCCTCTTTCAAGTTGTTGATGATATCTTA<br>GATGTTACTAAGTCCTCCGATGAACTCGGTAAAACTGCTGGCAAGGATTTGATCAGTGATAAAGCTACTTATCCAAAGTTAATGG<br>GTTTAGAGAAGGCTAAAGAGTTCTCTGATGAATTGTTGAATAGAGCAAAAGGCGAGCTAAGCTGTTTTGACCCAGTTAAAGCTG<br>CCCCTCTGTTAGGATTAGCTGACTATGTCGCATTCAGACAAAACTAA |
| *CrCPR* | ATGGACTCATCTTCAGAAAAACTATCCCCATTCGAACTTATGTCAGCCATCTTGAAAGGTGCAAAGTTAGACGGCTCTAATTCAA<br>GTGACTCTGGTGTTGCCGTATCCCCTGCCGTGATGGCCATGCTTTTGGAGAATAAGGAATTGGTCATGATCTTAACAACATCCGT<br>GGCAGTTTTGATTGGCTGTGTTGTAGTTCTGATATGGCGTAGATCTTCAGGTTCTGGGAAAAAGGTTGTCGAACCTCCAAAACTT<br>ATCGTCCCAAAATCTGTTGTGGAACCAGAGGAAATTGATGAAGGCAAAAAGAAATTCACTATCTTTTTCGGTACTCAAACTGGA<br>ACTGCAGAAGGATTTGCAAAGGCATTAGCTGAAGAGGCCAAAGCCAGATATGAGAAAGCTGTTATCAAAGTTATTGACATTGAT<br>GATTATGCTGCTGACGATGAGGAATACGAGGAAAAGTTTAGAAAGGAAACCCTAGCCTTTTTCATCCTTGCTACTTACGGAGAT<br>GGGGAACCAACTGATAACGCAGCAAGATTCTACAAGTGGTTTGTTGAAGGCAACGACAGAGGAGACTGGTTGAAAAACTTACA<br>ATACGGCGTATTTGGACTTGGTAATAGACAATACGAACATTTCAACAAAATCGCTAAAGTCGTAGACGAAAAAGTTGCTGAACA<br>AGGTGGGAAGAGAATCGTGCCTCTAGTGTTGGGCGACGATGACCAATGTATTGAAGATGACTTTGCAGCATGGAGAGAAAATGT<br>ATGGCCTGAATTGGATAATCTGTTAAGAGATGAAGATGATACAACCGTCTCAACAACATATACAGCAGCTATTCCAGAGTATAG<br>AGTGGTTTTTCCAGATAAGTCTGATTCCTTAATCTCTGAGGCCAATGGGCATGCCAATGGTTACGCAAACGGTAACACAGTTTAC<br>GACGCACAGCACCCATGTAGATCTAATGTCGCCGTGAGAAAGGAACTACACACTCCAGCATCAGATCGTTCCTGCACACATCTT<br>GATTTTGACATAGCTGGCACAGGCCTATCTTACGGTACAGGTGATCACGTCGGAGTGTACTGTGACAACCTATCTGAAACTGTTG<br>AGGAAGCAGAAAGATTGTTGAATCTTCCACCAGAGACTTACTTTAGTTTGCATGCCGATAAGGAAGATGGCACACCATTGGCCG<br>GATCTTCCTTGCCTCCTCCATTTCCACCTTGTACCCTGAGAACTGCTTTAACTAGGTATGCAGATCTACTAAACACCCCTAAGAAA<br>TCTGCATTGTTAGCTCTGGCAGCTTATGCAAGTGATCCAAATGAGGCTGACAGATTGAAATACCTGGCATCTCCAGCTGGTAAAG<br>ATGAATATGCTCAATCACTGGTTGCTAATCAAAGATCATTGTTGGAAGTAATGGCCGAGTTCCCTTCTGCTAAACCACCATTGGG<br>TGTCTTTTTCGCCGCTATAGCTCCAAGGTTACAACCTAGATTCTACTCTATTAGTTCATCACCTAGAATGGCTCCTTCTAGAATAC<br>ATGTAACTTGCGCTTTAGTTTACGAGAAAACACCTGGCGGTAGAATCCATAAAGGAGTTTGTAGTACTTGGATGAAAAACGCAA<br>TACCATTAGAAGAGTCCAGAGATTGCTCATGGGCTCCAATCTTCGTTAGGCAATCAAACTTTAAGCTACCAGCTGATCCTAAAGT<br>ACCTGTCATTATGATCGGTCCAGGTACTGGTTTAGCTCCTTTTAGAGGATTTCTGCAAGAGAGACTTGCCTTGAAGGAAGAGGGT<br>GCTGAACTTGGTACTGCAGTATTCTTTTTCGGCTGCAGAAATAGAAAGATGGATTACATCTACGAAGATGAATTGAATCATTTTC<br>TGGAAATAGGAGCTCTATCTGAATTGTTAGTGGCTTTTAGTCGTGAAGGACCAACCAAACAATACGTGCAGCACAAAATGGCCG |

| | |
|---|---|
| | AAAAGGCTTCTGACATATGGAGAATGATCTCAGATGGGGCCTACGTGTACGTTTGTGGTGATGCAAAGGGTATGGCTAGAGATG<br>TCCATAGAACATTACACACCATCGCTCAGGAGCAAGGCAGTATGGATTCCACACAGGCCGAAGGATTCGTCAAAAACCTTCAAA<br>TGACAGGTCGTTACTTAAGGGATGTTTGGTAA |
| *CrG10H* | ATGGATTACTTAACTATCATATTGACTTTGTTGTTCGCCTTGACATTGTATGAAGCCTTTAGTTACTTGTCACGTAGAACCAAAAA<br>CCTTCCTCCAGGGCCTAGTCCACTGCCTTTCATCGGCTCTTTACACCTTTTAGGAGATCAACCACACAAATCCTTAGCTAAACTAT<br>CCAAAAAGCATGGACCAATCATGTCACTGAAATTGGGCCAGATAACTACTATTGTGATATCTTCATCTACCATGGCTAAAGAGGT<br>ACTTCAAAAGCAAGACTTGGCATTTTCATCTAGATCTGTTCCAAATGCTTTACATGCCCATAATCAGTTCAAATTCAGTGTAGTTT<br>GGCTACCAGTGGCTTCTAGATGGAGGTCTTTGAGAAAAGTATTGAATAGTAACATTTTCTCTGGAAACAGATTGGATGCAAATCA<br>ACACCTAAGGACCAGAAAGGTCCAGGAACTTATTGCTTACTGTAGGAAAAACTCCCAATCCGGTGAAGCTGTCGATGTCGGTAG<br>AGCTGCTTTTAGAACTTCATTGAATTTGTTATCAAACTTAATCTTTTCAAAGGACTTGACTGATCCATACTCTGATTCCGCCAAAG<br>AGTTTAAGGATCTTGTGTGGAACATTATGGTAGAAGCCGGTAAACCAAATCTTGTGGACTTTTTCCCACTGTTAGAAAAAGTTGA<br>CCCACAAGGGATCAGACATAGAATGACAATCCATTTTGGTGAAGTCCTAAAACTGTTTGGTGGACTAGTTAATGAAAGACTAGA<br>GCAGAGAAGATCAAAAGGCGAGAAAAACGATGTCCTTGATGTCTTGCTAACAACTTCTCAAGAGTCCCCAGAGGAAATTGACAG<br>AACACATATTGAGAGAATGTGCTTGGATCTTTTTGTCGCAGGAACTGATACTACATCATCAACACTTGAATGGGCCATGTCAGAA<br>ATGCTGAAAAACCCAGATAAGATGAAAAAGACACAAGATGAACTTGCACAAGTAATCGGCAGAGGTAAGACAATTGAAGAGAG<br>TGATATCAACAGATTGCCTTACCTAAGGTGCGTAATGAAGGAAACATTACGTATCCATCCACCTGTTCCTTTTCTAATCCCACGT<br>AAAGTTGAACAATCTGTTGAGGTTTGTGGGTATAACGTTCCAAAGGGTTCTCAAGTTTTAGTGAATGCATGGGCAATTGGGAGA<br>GACGAAACAGTTTGGGACGACGCCCTGGCATTCAAACCTGAAAGATTCATGGAATCTGAACTTGACATAAGAGGTAGAGATTTT<br>GAACTAATCCCTTTTGGTGCTGGACGTAGAATATGTCCTGGTTTACCATTAGCATTACGTACTGTTCCATTGATGTTGGGCTCTTT<br>GTTGAACTCCTTCAATTGGAAGCTGGAAGGTGGCATGGCTCCTAAGGACTTGGATATGGAAGAGAAGTTTGGTATTACATTACA<br>AAAGGCACACCCTTTAAGAGCTGTGCCTTCTACCCTGTAA |
| *BsG10H* | ATGGACATCTTAAGCTCTATGCCGTGGTTCCTGTTTACTTGGATCCTAGTTTTCGCTCTACAGTTTATAGTTAAAGGACGTAAACC<br>CACCTCAGGCAAGTTGCCCCCAGGGCCGACGGGATTTCCAGTCGTCGGGAATTTACTTGAGCTGGGAGAAAAACCGCACAAATC<br>TCTAGCTAAGTTAGCAAAAGTGCACGGTCCTATCCTAAAGCTTGGTCAGGTAACAGCAATCGTTATTTCTTCTGCTAGTATGGCT<br>AAGGAAGTGCTACAGACGCATGACTTATTTTTCAGCTCCTGCCGTGCGGTCCCTGATGCATTACTAGCCCACCAACATAATGATT<br>TTTCCATGGTATTCCTTGCCGTAAGTACAAGCTGGAGAAATCTAAGGAAGATATGTAATTCCCATATCTTCACCTCACAGAAGCT<br>TGATGCTTCCCAGGAACTGCGTCGTAAGAAAGTGCAAGAATTGCTGGCCTATGTTCAAGAGAGCTGCCACGCCGGTCGTGCGGT<br>GGACATTGGACAGGCGGCGTTTACAACAACGCTGAATCTGCTGAGTAACACCCTGTTTTCTGTGGATTTAGCAGACCCCAGTTCA<br>GAGACCGTAAGGGAATTTAAACAGCTTGTGTGGGGAATAATGGTTGAAGCTGGGAAACCAAATTTGGCTGATTACTTCAGCGTC<br>CTTAAAAAAATTGATCCACAGGGCATTAGGAGACGTATGGCGACGCACTTTCAGAAAATGTTGGATTTATTCGACCAGATGATA<br>GACCAGAGACTAAAACTGAGACAAACCAATGGTACCGTAGGCACTAAGGACGTTCTTGATATTCTACTTAACATTATAGAAGAC<br>AACTCAGGCGAGATCGATAGAAACTATGTAAAGCACCTTTTTTTGGATCTGTTCGCCGCCGGGACCGATACAACGTCTTCAACGC<br>TGGAATGGGCAGTAGCCGAGTTACTTCACCACCCTGCAGCATTAAGCAAGGCTAGGCTTGAGTTAGAGCAAACTATCGGCAAAG<br>GTAATCCCGTGGAAGAAAGCGACATAACTCGTTTGCCGTACCTTCAAGCAATTGTTAAGGAAACATTCCGTCTTCACCCTGCAGT<br>ACCATTTCTGATCCCAAGAAAGGCGAGCGCAGATAAAGAAGCATTAGGTGGATTCACCGTACCTAAGGGGGCCCAATTGCTTGT<br>TAACGCTTGGGCCATAGGTAGAGACCCATCAACGTGGGAAAATCCTAACAGTTTTTCCCGGAGCGTTTTCTTGGGAGTGATTTA |

| | |
|---|---|
| | GACGTTAAGGGGAGGAACTTTGAGCTTATCCCGTTTGGTGCCGGGCGTCGTATTTGCCCAGGACTGCCACTTGCTATAAGGATGC<br>TTCACCTGATGCTGGGGTCATTGTTACACTCTTTCGACTGGAAACTGGAAGACGGGTCTACGCCTGAGAAGATGGATATGGACG<br>ATAAATTCGGGATTACCCTGCAGAAGGCTAGGCCATTACGTGCCGTTCCGATCGCAATATAA |
| *EgG10H* | ATGGATGTTTGGGTGTCATTTTTATGTATATGTCTTGTCTGGTGTCTTTTTGAAGCGCTTAGCTTCATCCGTAAGAAGTCCAAGAC<br>TACCTTGTCCAACTTACCCCCCGGTCCTCCACCTCTTCCTGTTGTGGCTAATTTACTTAGCTTAGGTAGTCAACCGCATCGTTCATT<br>AGCAAGACTTAGTCATACATATGGCCCGATTATCAAGCTTCAACTGGGCTACGTCACGACGATCGTAGTTAGTAGTCCCCCCATA<br>GCAAGAGAAATCTTACAGACGCACGACGCTATTTTCAGTGACAGGACTATCCCTGACTCTATAACCGCGCTTAGACAAAACGAG<br>TTGGGTTTGCCCTGGATCCCAGTAAGCCCATTATGGAGAAATTTGCGTAAAATTTGTAATTTGCATATCTTTAGCCATAAGAAGT<br>TAGAGGCCAACCAACATATCAGGCAGGAGAAAATCCAAGAGTTGCTTTCTTACATCAACGGATCTCTATCTAGAGGCGACGCGG<br>TTGACATAGGAGAGGTCGCATTTAAGGCAGTGGTTAATCTACTGTCCAAAACAATGCTAAGCGAAGATTTGGCCAATCCTTCTGG<br>GTCCGCCAAGGATTTCAAAGATTTGGTGTGGAGAATAATGGTTGAGGCTGGGAAACCGAATATTGCTGACTATTTTCCGGCGCTA<br>AAAAAGATAGATCCACAAGGGAGTAGACGTAGAATGACCGTATATTTTACAAAAATTTTTGAGTTGTTCGACTCATTGATCGAA<br>AAGAGGTTAAGAGACCGTGCCGCTGTCGGGTCTATTAGAAAAAACGATGTTCTTGATACCCTTTTAGACGCTCGTGAGGACAAA<br>AGCGAGGAGGTTGACATATTCCTTATAAAACACTTCCTGTTAGATCTATTCGTTGCGGGCACAGACACGACCAGTTCAACCGTCG<br>AGTGGGCACTTAGCGAATTGATCCATAGCCCCGAGAAATTGAGTCGTGCTCAAGCCGAATTAGATAGAGTAATTGGAAAGGGAA<br>AGCCAATAGAAGAGTCCGAGATTGCGCGTTTGCCCTATTTGCAAGCTGTCATTAAAGAAACGTTCAGGTTACACCCACCAGTGCC<br>GCTTTACCTACCTAGGAAAAGCGGCTCCGAAATCGGTATAGCTGGATTTACTATTCCAAAGGGTGCTCAGGTGTTCGTCAACGTA<br>TGGGCAATAGGAAGAGATCCATCTATATGGAAGGACCCTGAGGTTTTATGCCAGAGCGTTTTCTGGGCTCTGAGATTGACGTAA<br>AGGGGCAGGATTTTGAATTAACGCCCTTTGGTGCTGGTAGGCGTATCTGCCCTGGTCTACCACTGGCTCTACGTATGCTACATTG<br>GATGCTAGGGTCCCTAATTAACTCCTTTAACTGGGAATTGGAAGGCAAAGTAAAGCCGGAAGACCTATCCATGGAAGAAAAGTT<br>CGGTATAACCCTTCAAAGAGCTCAGCCACTAAGGGTTATTCCTAAACCGTTGTAA |
| *GrG10H* | ATGAGAGAAATGGATCTGCTTGCAAGCTCCTTACTTGGTTTACTGCTTACCTGGTTCTTGTTCCAGGCTTTTTTGTCCATCAAAAA<br>CGGGAACAAAAGCTCACAACGTAAACTGCCACCAGGGCCGAGACGTATTCCGATATTCGGGAATTTATTCGATCTGGGCGATAA<br>ACCACATCGTAGCCTAGCGAAGTTCGCGCAGATTCATGGGCCTGTAATGAGTTTAAAACTTGGATCTTTAATCACAGTCGTCGTA<br>TCTAGCGAAACCACTGCGAAAGAGATATTGCAAAAACAAGATTTAATTTTCTGTAACCGTACCATAGTGGACGCAATACGTGCT<br>AGCCAGCACCACGAATTTGGCATGCCGTGGATTCCAGTCAGCCCGCTATGGAGAACCTTACGTAAAGTCGGTAATACACACATC<br>TTTTCATCTCTTAAACTGGACGCCAACAAATACCTGAGGAGACATAAAATTCAGCAACTGATAGCAAAGGTCGGCGAATCATGC<br>CTAAAATGTGAAGCAATCAACATAGGACAGGCGGCCTTCGATACGACAATAAACCTGCTAAGCAATACGATGTTTTCTGTTGAC<br>CTAGTAGACCCGAACTCTGCTAGAGCGCAGGTATTCAGGAAGACGGTATATAGCATCATGGTCGAAGCGGGGAAACCCAACTTA<br>GCCGACTACTTTCCGCTGCTGAGGAAGATGGACCCGCAAGGCGTCCGTAGAAGAATGACGGTCCATTCCGATAAACTTCTAAAA<br>CTTTTCGGAAACATGATGGACGAAAGGCAACAGAGTAGAAAATCTCCAGACTATACAGCGAGCAATGACGTTCTTGACACGTTA<br>CTAGACATTATTGAAGGCGACATAGAGGAACTTAATAAGGATCATATCAAACATCTGTTTTTAGTCCTATTTGTCGCCGGTACCG<br>ATACTACTTCTAACACACTAGAATGGGCCATGGCAGAAGTCCTACGTAATCCGCACGTACTGTTAAAGGTAAAAAAGGAATTGG<br>ATCAGGTGATCGGCAAAGGAAAACCAATAGAGGAGAGCGATATTAACAGCCTACCGTACCTGCAGGCGATAATCAAGGAGACG<br>TTTAGGATGCACCCTGCAGTGCCTCTGCTACTGCCACGTAGAGCCGGCTCTGATACCGACCTGTGTGGCTTCCACGTCCCAGAAG<br>GTTCACAAGTGCTAGTAAACGCCTGGGCGATTGGAAGGGACAGCTCCATCTGGGAGAACCCAAATTCCTTCATGCCAGAGAGGT |

| | |
|---|---|
| | TCCTAGGGTCTGAAATAGATGTCAAAGGGAGGGATTTTGGTTTAATTCCGTTTGGTGCGGGTAGACGTATTTGCCCTGGATTGCC<br>GCTAGCGAATAGGATGCTTTACCTGATGTTGGGTAGTCTAATTAACTCTTTTGACTGGAAGTTAGAGGGGGGGATCTCACCCCAG<br>GAGATGAACATGGAAGAGAAGTTTGGACTTACCGTACAAATGGCCGAACCATTACAGGCTATCCCAGTTGTGATATAA |
| *GriG10H* | ATGTTGGGCTCAAATAAGTCCTCTAGCCCCTCCACGTCTCACCATTCTATACACCCGTTTATAGCAATGGATTATCTAACCATTGC<br>CCTAGGTTTCTTGTTCGCATTAACACTGTATCAAGCTTTAATTTCCTTCAGTCGTAAAAGTAAGAACCTTCCCCCTGGGCCAGCCC<br>CTCTGCCCTTTATTGGCAATCTACACCTTTTAGGAGATCAGCCCCACAAGTCTTTAGCTAAGTTAGCCAAGAAACACGGCCCGAT<br>AATGGGCCTGCAATTCGGGCAGGTCACGACGATCGTAGTTACTTCTTCCGGGATGGCTAAGGAGGTGTTACAGAAGCAGGATTT<br>AGCGTTTAGTAGCAGGTCTGTACCAAATGCTCTACACGCCCATGACCAATACAAGTACTCTGTAATTTGGCTGCCTGTGGCGTCA<br>CAATGGAGGAGCCTTAGGAAGGTGTTAAACTCTAATATATTCTCTGGGAATTTAGATGCTAATCAACACTTACGTTGTCGTAAGG<br>TGCAGGAGCTTATTGCATACTGCAGGAAGAATTGTCAGACTGGGGAGGCAGTGGATCTAGGGAGGGCGGCTTTCAGGACTTCAC<br>TAAATTTGCTAAGCAACACGATATTTTCCAAGGATTTAACTGATCCTTATTCAGATTCTGCTAAAGAGTTCAAGGATCTGGTCTG<br>GAACGTTATGGTCGAAGCAGGTAAACCCAATTTAGTTGACTTCTTCCCGGTCTTAGACAAGGTCGACCCTCAAGGGATTAGGAA<br>GAGAATGACGTTCCACTTCGGCAAAATTCTGCAACTGTTTGGTGGATTGATCAATGAGAGGCTGCAGCAAAACAAAGCAAAGGG<br>CGCCCATAATGATGTGCTAGATGTTTTACTTAAAACGAGCCAAGAGACACCAGACGAATTAAATAGAACCCACATAGAAAGGAT<br>GTGTCTGGATTTATTTGTGGCCGGTACAGATACTACTAGCTCCACGTTGGAGTGGGCAATGGCTGAGATGCTAAAGTCACCTGAT<br>AAAATGAATAAGGCGAAGGAGGAACTAGCTCAGGTGATCGGCAAGGGTAAGGCAGTCGAGGAAGCCGATATCGCCTCATTGCC<br>TTATCTGAGATGCGCCATTAAGGAGACCTTAAGAATTCACCCTCCCGTACCGTTTCTTATCCCACGTCGTACAGAGCAGGAAGTG<br>GAAGTCTGTGGCTACACCGTCCCCAAGAACTCACAGGTGTTCGTCAATGTTTGGGCTATTGGTAGGGATGAAACCACATGGCCA<br>GATGCGTTGGAATTTAAGCCCGAAAGATTCCTAGAAAGCGAAATAGACATGAGAGGGAGAGATTTTGAACTATTACCGTTCGGC<br>GCTGGGAGAAGAATTTGTCCTGGTTTCCCTTTAGCGGTCAGAATGGTACCGGTGATGTTGGGTTCATTGTTAAATTCTTTCGACTG<br>GAAACTTGAAGGTGGGATCGCACCCAAAGACCTTGACATGGAGGAGAAGTTTGGGATTACGCTTCAGAAAGCCCACCCCCTGTG<br>CGCCATCCCAACATCCCTTTAA |
| *HaG10H* | ATGGGTTTTGTCATCGTTGTAGGGTTACTTTTGAGTTACATTCTGATTAGGGCTACATTCTTAGTCTTCGGGGTCGGACGTCCCAA<br>GAACTTGCCACCTGGACCTACATGGCTTCCTATTATTGGAAATTTGCACTTCCTTGGGGACCAGCCACACAAGAGTTTAGCCAAA<br>CTTGCTGAAACACATGGTCCTGTAATGTCTCTTAAGCTTGGGCACATAACTACCGTTGTTATATCTTCCGCCTCTGCCGCAAAGGA<br>AGTTCTGCAGAAGCAGGACCTAGCTTTTTCTGCTCGTCACGTACCCAATGCTGTCCACGCAAGAAACCACGCTGGACACTCCGTA<br>GTATGGCTACCCGTAGGAACTAGATGGCGTACCTTGAGAAGGATTCTTAACTCAAATATATTCTCCAGCAATAGTTTAGAAGCTA<br>ACGAGCACTTAAGAAGCAAAAAGATAGAGGAATTGATAGCATATTGTGGCAAGGCGAGTGTGTCAAATGATTACGTTGACATTG<br>GAAGAGCGGCATTTCGTACAAGCTTGAATCTGTTATCTAATACTATTTTTAGCAAGGATGTTACAGACCCATATGAGGAAGACTC<br>CGGCAAGGAATTTAAAGAGGCAATCACGAACATCATGGTAGAAGCAGGCAAGCCCAACTTAGTTGATTTCTTTCCCGTGCTGAA<br>GAAGATCGACCCTCAGGGAATACGTCGTAGACTTTCCGGCCATTTTGGTAAATTGCTAGAGATGTTTGAGGAGCTGATTGATGAA<br>AGACTGAGGATGGGGCGTTTAAAACATAATGATTTACTGGACGTGTGTCTGAAAATAATCGAGGACAATTCCAACGAGATAAAT<br>CATACGCACATAAAATCTCTTTTCCTTGATCTGTTCGGTGCTGGGACGGACACTACTTCCAACACATTGGAGTGGGCAATGGTAG<br>AGGTGCTGCGTAATCCAGACACAATGAGCAGAGCGAAAGAGGAGCTTGAAGAGGTAATCGGCAAAGCGAAAATCGTGAAGGA<br>GAACGACGTGCTAAGGTTGCCATACTTAAGCTGCATTGTTAAAGAGACGCTGAGACTACACCCCCCTGTACCACTTCTTATCCCC<br>AGAAAAGTTGTTAAAGAGGTACAATTGAACGGATACACGATCCCTAAAGGGACCCAGGTTTTAGTTAACGCATGGGCGATTGGA |

| | |
|---|---|
| | AGAGACCCGACGGTCTGGGACGACAGTCTTGAGTTTAAGCCGCAGAGATTTCTTAAGAGCGGGTTTGATTTTCGTGGCAAGGAT<br>TTCGACCTTATACCCTTTGGTGCAGGGAGGAGGATGTGTCCCGGCTTGCCACTTGCAGTCAGGATGATACCGATTATGCTTGGGT<br>CCCTTTTGAACAACTTCGATTGGATCCCAGATACCAAGAATCAAGCCGATACTCTGGATATGAATGAAAAATTTGGTATAACATT<br>AAGCAAAGCTAACCCACTGTGTGTCGTGCCGATACCCCTAAATTAA |
| *HrG10H* | ATGGACTTCTTGGTATTCGTGATTAGTTGCGTGTTTGCTTGGACTGTGTTCAGCACCTTGACGTCCCTTAGTAGAAGGAAGAAAA<br>GTAAGCTACCTCCGGGACCGACCCCTCTGCCGCTGATCGGCAATCTGCACCTTTTGGCGGGCGCACAACCCCACGATCTGGCAAA<br>TTTGGCCGCGAAGTACGGACCAATTATGTCCCTTAAGCTAGGCCAGGTTACCTCAGTTGTTGTGAGCTCCCCTGAGATGGCAAAA<br>GAGGTGCTTCAGAAGCAAGATTTAGCGTTCTCATCAAATCGTTTTTTTCCTGATGCTGCGACAGCCCTGGACTACCATAATCATTC<br>TCTAGGTTGGTTGCCGGTTGGCAGCAGATGGAGATCACTTAGAAAGATCGTGAACACGAACATATTTTCCGGCAGTATGTTAGA<br>AGCGACGCAAGATTTAAGATCAAGGAAAGTTGAGGAGCTAGTCGAATATTGTAGGAGGAAAAGCAACACCGGAGAATCAGTAA<br>ATATAGGTGAAGCCACCTGCGGGACAGTAGTTTCTATACTGTCAAATACGATTTTTTCCAAAGATATAACCGATCCCTATTCTGA<br>TTCTGCTCGTGAATTTAAGGAATTACTGCGTAATTTCGTCAGTGAGATGGGAAAGCCTAACCTAGTGGATTATTTCCCCGTCTTAC<br>GTAAATTTGACCCCCAGGGTGTCAGACGTAGAGAAACTACTAACATAACTAAGGTATTTGAGAGGTTCGACGACTTAATGAAAG<br>AACGTTTAGAGCAGCGTTCCAAGGGCATAACGAAGAACGATGACGTACTTGATGTATTACTGACCATCTGTGAAGAAAACTCTG<br>GCGAGTTCGTCAGACAGCAAATACACCATCTATGTCTTGATCTGTACGGCGCTGGCACAGACACTATCACAAGTTCAGTCGAGTG<br>GTCTATGGCCGAATTAGTTAAGAACCCCGAAGCCATGACGAAGACAAAGGAAGAGCTAGCCCATGTCGTTGGATCAGGTAAACC<br>GATAAAGGAGGTAGACCTGCCCAACCTGCCTTATTTACAATGTGTTGTAAAAGAAACGTTAAGGATGCACCCGCCCGCGCCCCT<br>GCTTTTGCCCAGAAGGGTTGAAAAGGATGTGAGGGTTTGTGAGTATATCGTCCCTGAGGGGTCTCAAGTTGTTGTCAATGCAATG<br>ATTGGTAGAGACGCTCGTCTGTGGGAGAATCCCCTGCGTTTTATACCTGAGCGTTTCTACAGAGTGGCGTCGATATACACGGAC<br>AAAACTTCGAGGTAATACCCTTTGGGGCCGGGAGGCGTATTTGTCCAGGCCTACCACTGGCGACGAGGATATTGCCGTATGTAA<br>TAGGCTCACTATTGAATTGCTTTGATTGGAAACTAGAGGGTGACATTACGCCTAACCAATTAAATATGGAGGAGACCGTAGATA<br>TTATACTACACAAAGCTGACCCACTATTAATAGTACCCGTTTAA |
| *InG10H* | ATGGATATGGACTACCAGAGGATCTTGATCGCGCTTTGTCTAGCTTGGACTCTAGTTCAATGCCTTCGTTTGTTATTGGCCAAGAG<br>GGGGAGAAAATTCCCCCCCGGCCCGTTCCAACTGCCTGTTGTAGGTAACTTGCACCTTCTAGGCGACTTACCACACAAAAGCTTA<br>GCACGTCTTGCTGGGAAATATGGCCCCGTTATGAACCTTAAGCTAGGTATGATCAACACTGTAGTTATTTCAAGCTCTACTATGG<br>CGAAAGAGGCTCTTCAAAAGCAGGACCTGACCTTCTCCACAAGATCTATTCCCGACGCGTTGAGAGCGAGAAACCACAGTCAAT<br>TCTCAGTTGTGTGGCTGCCGGTAGCTTCCAAATGGAGAACCCTAAGGAAAGCGATGAATAGTAATATTTTTTCAGGATCTAAGTT<br>GGACAATAATCAACATTTAAGAGTAACGAAAATACAGGAGCTGATAATATATTGCCAGAACAAGAGCCAAGCTGGAGAGGCAG<br>TCGACATAGGTCGTGCTGCCTTTAGAACTACTTTGAATTTACTTTCAAATACTATCTTTTCTAAAGACTTGACCGACCCATACTCT<br>GATTCTGCTAAGCAGTTCCGTGATCTAGTGTGGAATATGATGGTAGAAGCCGGTAAACCGAACCTTGTGGACTACTTTCCGTTAT<br>TAGAAAAATTCGACCCACAAGGTATCAGGTGTAGACTGACGGGGCATTTTACGAAAGTTCTGGAATTGTTTGAGGACTTAATAG<br>ATGAGAGACTAGAAGAGAGAAAAGTAATGGGTTGCAAAAACGTTGATGTGCTGGACAGTCTGCTAAATATATCTCAGGAAAGA<br>CCTGAGGAGATTGACCGTAACTATATACTGCATGTCTTCCTGGACCTATTCGCAGCGGGAACGGATACCAGCAGCAACACCCTA<br>GAATGGGCAATGACTGAGCTTTTGAAGAATCCGAATGCCCTTGCAAAGGCGCAGGCTGAGCTGGCGGACGTAATCGGGAAAGG<br>TAAATTGATTCAAGAAGCAGATGTTACAAGGTTGCCGTATCTTCAGTGCATAGTCAAGGAAACCTTCAGAATGCACCCCCCGGTT<br>CCGTTCCTTATTCCCAGGAAGGTAGAGCAGGAAGTCAATCTATGCGGTTATACTGTGCCAAAAGATTCTCAAATATTGGTCAATG |

| | |
|---|---|
| | TTTGGGCCATAGGCAGAGATTCCAGCATCTGGGAAAACCCACTTATCTTTAATCCGGAAAGATTCTGGAATTTCGGCATCAACGT<br>GAGGGGGCAGGATTTCGAGCTGATTCCGTTCGGTGCTGGTAGGAGGATCTGTCCTGGTTTACCTATGGCTATGAGAATGGTTCCT<br>GTTATGCTGGGAAGTCTATTAAATAGCTTTCAGTGGAAGCTTGAAGATGACATTGCTCCTAAAGACCTAGATATGGAAGAGAAA<br>TTCGGCCTGACCCTAGCCAAGGCTCACCCCCTAAGGGCTATACCGATTCCCTTCTAA |
| *JrG10H* | ATGGACTTTTTGGGGTTGCTGCTTTGTGCCATATTTACGTGCATATTAGTACACTCTCTTAATAGCTTACTGGGGTGGAAAAAAAG<br>ATCTGCAATCAAGTATCCCCCAGGTCCCAACGGTTTCCCAATTATCGGGAATTTATTTGAGCTGGGTGATAAACCGCATATGTCC<br>CTAGCTAAACTTAGCCAGATTCATGGGCCGATCATGTCCTTAAAACTGGGTTGTATTACGACAGTTGTAGTCTCATCACCTCTGA<br>TGGCCAAGGAGGTCCTACAGACACATGATATCTCCTTCAGCAACAGGACCGTCCCAGACGCTTTGAGAAGCTATAAGCATGATG<br>AGTTTGGACTACCTTGGATGCCAATGTCTTCCCGTTGGAGGTCCATCAGGAAGATCTGTAATTTACACATGTTTAGTGGGAAAAA<br>GCTTGATGGCAACCATCACTTACGTAGATCTAAGGTCCAGGAACTACTGTCAGAGGTCCAGAAATGCTGCGAGGCCGGTTGTGC<br>AGTAGCAATAGGAGAGGTAGCATTCGGGACATCACTTAATTTCTTGTCCTCTACTATATTTTCTATGGACCTTGACTCCCAACCTG<br>GGAGCGGGTCAGTACAGGATTTCAAAACACTTGTTCGTCATATGACGGAGGCCGGAGGAAAACCAAACCTAGCCGATTATTTCC<br>CTATCCTTAGAAAAATTGACCCGCAAGGTATTAGGTCCAGACATAAGATATTTACAGGTGAAATGATGCAGTTGTTCAACCAGA<br>TGATAGCCGTCAGGCTTCAACTACGTAAGGTGCCCGGATACGTCACCTCCAACGACATGCTGGATGCGTTACTTGATATATCAGA<br>GAACAACTCAGAAGAGATCGACACAAGCCGTATCGAGCACCTGCTGTTAGACCTGTTTGTGGCAGGAACTGACACCACGGCGTC<br>CACACTTGAATTTTCTATGGCGGAGCTATTAAGAAATCCTGAAACCTTGATGAAAGCAAGGGCTGAGCTTGAAAATACCATAGG<br>CAAGGGAAACCCAGTGCAAGAATCTGACATAAGTCGTTTGCCATACCTTCAGGCTATAGTCAAGGAAACGTTTAGGCTACATCC<br>TACGGCTCCGCTTTTAATTCCGAGAAAGGCTCCTGACGCTGATACAGCAATCGCAGGATTCACTGTGCCGAAAGGTGCTCAGGTA<br>CTTGTTAACGCTTGGGCGATTGGCCGTGACCCCAGTACTTGGGATAATCCCATATCATTTGAGCCTGAGAGATTTCTAAGAAGCG<br>ATATAGATGTCAGGGGACAAAATTTCGAATTGATTCCATTCGGCAGCGGGAGGCGTATATGCCCTGGCCTTCCGCTAGCGATAC<br>GTATGATCCATCTTATGTTAGGATCCTTAATACACTCATTCGACTGGAAATTGGAAGACGGCGTATCTACCGAGGACTTGAACAT<br>GGAGGAAAAGTTTGGGATCACTTTGCAGATGGCGCACCCTTTACGTGCTGTTCCAATTCTGAGATAA |
| *PkG10H* | ATGGATTTTTTAACTATAATTCTGGGTTTGCTATTTGCTGTAACCTTGATCCACGGTTATCAACTTCTTTCTTCCAAAGGAAAGCG<br>TTTGCCACCCGGCCCGACTCCTCTGCCATTGATCGGATCCCTACATCTATTGGGGGATCAGCCACACCAAAGCTTAGCGAAACTA<br>GCAAAGAAACACGGCGAATTGATGTGCTTAAGGCTAGGTTTTATCAACACAATCGTAATTTCCTCCGCCGCAATGGCGAAGGAA<br>GTTCTGCAAAAGCAGGACCTAGCATTCAGCAGTAGGATGTCACCCAACGCAGTACACGCCCACGACCAATTCAAATATAGCGTG<br>GTCTGGCTTCCGGTAGCCGCGCGTTGGAGGAGTTTACGTAAAGTACTAAATTCAAACATTTTCAGTGGGAACAGACTGGACGCC<br>AATCAGCACCTTAGATGCAGAAAAGTTCAGGAGCTGATAGCCTACTGCAGAAAAAGCTCCCAGACGGGAGCGGCAGTAGACAT<br>GGGGAGAGCCGCCTTCAGAACCTCTCTTAACTTGTTATCAAATACAATTTTTAGCAAGGATCTGACGGACCCCTTCTCAGACTCC<br>GCAAAGGAATTTAAGGAGTTAGTTTGGAATATCATGTTGGAGGCAGGTAAACCCAATCTAGTAGATTTTTATCCCGCCCTTGAGA<br>AGCTGGACCCCCAGGGTATCAGGAAACGTATGACGGTTCATTTCGGTAAGGTGATTGAACTATTTTCAGGATTGATTAACGAAA<br>GGACTGAGACTGGGAAAACCCAAATGTTAGGAACTACGGACGTGATAGACGTCCTATCCGATTCAAAGGAGACAAGAAGGATT<br>GACAGAACCCACATTGAGAGATTATGTCTTGACCTTTTCGTCGGCGGCCTAACTCATATCAACACGCTAGTGTGGGCAATGGCTG<br>ATCACGTAAGGACGAGGAAGTATTGCAAGAGCAGACTAACAAGGACAGTAATCGGGAAGGGCAAGTTTTCCACGAAGCAGAAC<br>ATCCAGCGTCTGCCATATCTGAGATGTATGGTAAAAGAGACACTGAGGATTCATCCTCCAGTCCCATTTCTGATACCCAGGAAGG<br>TCGAACAGGATGTGGACGTATGCGGTTATACGGTCCCTAAAAATTCACAAGTATTTGTCAATGCGTGGGCAATTGGAAGGGATC |

| | |
|---|---|
| | CTGAGACATGGCCCAATCCGTTGGAATTTAAACCGGAGCGTTTTATGGAAAGCGAGGTTGATATGCGTGGGAGGGATTTTGAAC<br>TTATTCCATTTGGCGCTGGAAGACGTATTTGCCCAGGAGTTACGCTGGCTGTCAGGATGGTCCCTGTTATGCTAGGTTCCCTATTG<br>AATTCATTTGACTGGAAATTGGAAGGGGGAGCCGGACCAAAAGATTTGGACATGGAGGAAAAGTTCGGGATAACATTGCAAAA<br>AGCGCTGCCTCTTATGGCTGTACCAATACCACTGTAA |
| *PpG10H* | ATGGACTTTTTGACCATCGTGGTTGTAGGTATTCTTTTTGCTATTACTTTGATTCAGGCCATCAGATCTCACTCTACTAGAGGGAA<br>AAATTTGCCACCCGGGCCTCATCGTTTACCGATCATCGGAAATCTGCATCTTCTGGGAGACCAGCCGCAAAAATCAATGGCAATG<br>TTAGCTAAAAAGCACGGGCCCGTTATGTCTTTACGTCTGGGACGTATAACTACGGTCCATATTGCCTCCGCAGCCATGGCGAAAG<br>AGGTGTTACAGACGCAAGATTTAGCATTTTGTAACAGGACGGTTCCCGACAGTCTTAAAGCTCACAATAACAACAATTACAACG<br>TAGTTTTCCTACCAGTCGGTTCTCGTTGGCGTAGTTTGAGGAAGGTGATCAACTCCTCTATATTCAGTTCTAACAGTTTGGATGCC<br>AACCAGGAGCTAAGGTCCTGTAAAGTAAGGGAACTTATCGCGTATTGCAGAAAGAAAAGCCAAGCGGGTGAGGCCGTCGACTT<br>AGGGAACTCAGCATTTAGGACGGCATTGAATCTTCTGTCCAACTCTATTCTATCCAAGGATCTAGCTGACCCCTTCTCAGATTCTG<br>CGGAGTTCAGAAACTTGGTCTGGAATATCATGGTCGAGGCGGGGAAGCCAAATTTAGTAGACTTTTTTCCTTTCCTTGCGAAGAT<br>TGACCCCCAGGGTATGAGGAGAAGGCTTACTATATATTTTGGGAAGTTGTTGAAACTTTTTTCCGACTTGATCGACGAACGTCTA<br>GAAAAAAAATCTTTCAGGGGCGGTAAGACAGACGCTCTTGATATATTACTTTCAATAACTCGTGAAAGTCCCGAAGAAATAGAT<br>AGGATCCATCTAGAGCACATGTTAATGGTCCTGTTCGTCGCAGGAACGGATACAACCTCTAGCACGGTTGAGTGGGCCATGACT<br>GAGACACTGAGAAACCCTGAGGTAATGAAAAAAGCGAAAGCTGAACTAGAACAGGTAATTGGGAAGGGAAAGATGGTAGAAG<br>AGGCGGACATATCTAGACTGCCCTACCTACGTTGCATGCTGAAGGAGACCTTCCGTATCCACCCCCCCGCTCCATTCCTAATACC<br>GCGTAAGGTTGATCAAGACGTGCAACTGTGTGGGTACACCGTCCCAAAGGATAGTCAAGTGTTAGTGAACGCTTGGGCAATCGG<br>TAGGGATTCTAGTACATGGGACTCCCCCTTAGAGTTCAAACCAGAGAGGTTCTTGGAATCCGAGATAGATGTTCGTGGCAGGGA<br>TTTTCAACTTATCCCATTCGGGGCCGGTAGAAGAATATGTCCCGGGCTGACTCTAGCCATGAGAGTCTTGCCCGTCGTCTTAGGG<br>TCACTGCTTAACCTATTCGATTGGGAGATCGAGGGAGGTTTGGCACCCGAGGAGTTGGATATGGAAGAAAAAGTTGGAATTACC<br>CTGCAGAAAGCCGTCCCATTAAGAGCCGTCCCAATACCAGTCTAA |
| *RcG10H* | ATGATGGACCTTCTTGTAGGCTGCGTTTTAAGTTTACTGTTTACTATAACTCTTGCGCAGGCCCTGACTAGCATTAGTAAACGTTC<br>CAAGACTGGACCAGGAAAACTACCTCCTGGACCCACACCCCTGCCTTTAGTTGGGAATCTGTTCGAGCTTGGGGATAAACCTCAT<br>CAGAGTCTGGCAAAGTTAGCTAAAATACATGGCCCGCTTATGTCCTTAAAATTGGGGCAAATCACCACGGTGGTTATCAGCAGC<br>GCAACCCTGGCGAAGGAAGTGCTGCAAACTCTTGATCTTTCTTTCGCTAATAGAATTTGCGTCCAAGCAGTTCACGCTCACGATC<br>ACCATGAAGCGTCAATGCCGTGGTTGCCCGTGGGAGCACCCTGGAGAAACCTAAGGAAAATATGCAACAGCTACTTATTCAGCA<br>ATCAGAAACTTGACGGCAATCAAGACATCAGACAAAAAAAAATCCAGGAACTGATTGCCGATGTTAAGGAAAGCTGTAGGCTG<br>GGAGCAGCTACTAACATCAGTCATGTCGCTTTCAAGACCGTGCTAAGTGTCCTGTCATCCAACGTTTTCAGTTTAGACCTTACTG<br>ATTCTAACAGTGACTCTGTGAGGGAGTTTAAGGAAGTCGCCAGATGTATTATGGATGAAGTAGGGAAGCCGAACTTGGCGGATT<br>ACTTTCCGGTATTAAGGAAAATTGATCCGCAGGGAGTGAGAAGAAGAACTGCGATTTATTTGGCAGGATGCTGGATCTATTTG<br>ATCCAATCATCGATCAGAGGTTGGAATTGAGAAAGGAGGAGGGGTATATATCCGCCAATGACATGCTTGACACGCTTTTAGCAC<br>TGATCGAGGAAAATAAGACTGAAATGGACATTAATTCCATGAAGCACCTGTTCCTTGATCTTTTTGCAGCAGGCACTGACACCAC<br>TTCTTCTACCCTAGAATGGGCAATGACTGAGCTGCTTAGGAATCCCAAAACACTGTCAAAGGCGCGTGCAGAGATTAAACAGAC<br>AATAGGAACGGGCAGTTTGCTGCAGGAGAGCGATATGGCCCGTCTGCCGTACCTAAAAGCGATCATTAAAGAAACCTTCAGGTT<br>GCACCCCGCAGTGCCTCTGCTACTTCCCAGAAAAGCAGGAGGGGATGTCGAGATGAATGGGTTTACAATTCCCAAAGACGCCCA |

| | |
|---|---|
| | GGTGCTTGTTAACGCCTGGGCGATTGGAAGGGACCCCTTTTTGTGGGAAGAACCGGAGCTGTTCCGTCCGGAACGTTTTTTGGAG<br>TCCAACATAGATGCACGTGGCCAGTATTTTGAGCTAATTCCTTTTGGTGCAGGACGTCGTATTTGCCCAGGATTACCCTTGGCGA<br>TCCGTATGTTGCACTTACTACTAGGGTCATTAATTTATTCATTTGATTGGAAGCTTGAAGACGGGGTGACACCCGAAAATATGGA<br>CATGGAAGACCGTTTCGGTATATCTCTTCAGAAGGCCAAGCCATTGATCGCCATACCCAATCAGGTCTAA |
| *SaG10H* | ATGGATTTCCTTGGTTTTATACTATATGTCCTGTTCGCCTGGGCTCTTATCAGAGCGCTACGTTCCCTGTCTAGAGGATCCAAGGC<br>AGCGGGAGGTAGGTTGCCACCTGGGCCTGTTCCACTACCTGTTGTCGGCAATTTGTTCCAATTAGGGAATAAGCCGCACAAATCT<br>CTGGCAAAGCTTGCCAAATCTTATGGGCCTATAATGAGTCTTAAGTTAGGGAGAATAACTACCATAGTGATAAGTGCATCTACTG<br>TTGCCAAAGAGATCTTGCAAAAGCAAGATCTGACATTTTCAAACCGTCACATCCCCGACGCAGCGCGTGCGCATCGTCACGATCT<br>ACATTCTATGGCGTGGCTGCCAGTGAGCACTAGGTGGCGTACGTTAAGAAAAATTTCTAATTCTCACATCTTCACAAGTCAACGT<br>CTAGATGAAAACCACCATTTAAGACGTCAAAAACTAGACGATCTTCTAGCTTGGGTAGCTGAAAGTTGTCAGGTAGGGGCCGCT<br>GTCGATATAGGAGCCGTCGCATTTATGACTAGTCTAAACTTGCTAAGTAACACCGTATTTTCTAAAGACTTGGTTGAACCTGGCT<br>TAGGTGCTGCGCAGGAAATGAAAGAGGTGGTTTGTGGCATTATGGAGGAGTTGGGCAGACCTAACCTGGTGGATTATTTTCCAG<br>TATTAAGAAGATTTGACCCACAGGGTATCAGGAGGAGACAAACCGGGAACTTTGGAAAGATGTTCGAAGTGCTTGGGGATATCA<br>TTGATCAGAGATTGCAGTGGAGGAAGCAGAGATCTGATGGAGACAGCCCCGCTGCTACAACGAAAGATGTGTTGGATGTTCTGC<br>TGAATATAATTGAAGATACCGAAGTCGAAGAAAAACCAAATAGAACGGAAGTCGAGCATCTGCTTGCAGACCTTTTCGTTGCTG<br>GGGGCGATACGACGAGCAGTACCGTTGAGTGGGCGATGGCGGAGTTACTTAGAAAACCCGAGACCCTTGAGAAGGCGCGTCAA<br>GAGCTGCATGAAACAATAGGACCCAGAAACCCTGTCCAAGAAGCCGATATACCGAGGTTGCCATATCTGCAGGCGGTCGTTAAA<br>GAAACTTTTCGTTTGCACCCGGCAGCCCCCCTATTGGTACCGAGAACTGTAGAGAATGACGCTGAACTATGCGGTTTCACTGTCC<br>CAGCCGGAGCGCAAATTATGGTGAATGCATGGGCGATAGGAAGGGACCCTGGAACATGGGAAGACCCAGAGTCATTCTTGCCC<br>GAGCGTTTCTTGTGGTCCGATGTGGATGTCAAGGGCAGGAATTTCGAACTTATTCCATTCGGCGGCGGTCGTAGGATCTGTCCTG<br>GCCTACCCCTTGCGATCAGAATGGTACATCTAATGTTGGGCAGCCTGATACACGGGTTTCGTTGGAAATTGGTAGATGATGGTAT<br>GGGGAGCCCCGAAACGGCAATGGACATGGACGACAAGTTTGGCATAACGCTGCAGAAAGCCAAACCATTATGTGCTGTACCCAT<br>TCGTTAA |
| *StG10H* | ATGGAGTATGTAAACATCCTTCTGGGACTGTTATTCGCCTGTTTCCTGGTTCGTGTCGTATTTATAAGCTTGCGTAGGTCCAAAAG<br>AGTCGCTCCGGGTCCTTTTCCCTTGCCCATCATTGGGAACCTACATCTACTGGGCGACAAGCCACATATATCATTAACGCAGCTT<br>GCGAAGAGATATGGTCCGATTATGAACCTAAAACTGGGGCAGATAAATACAATCGTGATCTCCAGTAGTGTGATGGCCAAACAA<br>GTAATGCAGAAGCAAGATTTAGCCTTCAGTAACCGTTTTGTGCCGGATACCCTGCACGCATGCAATCACTCCAATTACTCAGTTA<br>CATGGATCCCCGTAAACAACTCTCAGTGGAAGACGATACGTAAGATAATGAACACCCACATCTTTTCAGGCAACAAGCTAGATG<br>CAAATAAGCACATCAGAACAAAAAAAAATCCAAGAACTGATCGACTATTGTCACAAGAACGGGGAAAATGGTGGTGCCGTCAAT<br>ATTGGGGGTGCGACATTCAGGACATCTCTGAATCTACTTTCCAATTCCATCTTTAGCAAAGATTTAACAGACCCCTTCTCCAATA<br>GCTCCAAGGATTTCAAAGAATTGGTATGGAATATTATGGTCGAGGCAGGCAAGCCAAATTGGGTGGATTATTTCCCATTCCTTCA<br>AAAAATTGACCCTCAGGGCCGTAGGAGACGTATGACAGATTACTTTACTAAGGTTTTACATTTGATTTCAGGCTTTATTGATGAG<br>AGGATTAAGGAGAGAGAGATGGGTAATCACGCCAACGTCGACGTACTGGACGCGCTATTGAATATTAGCCCAGAAGAAATAGA<br>TAGAAACCATATAGAACACCTGTGCTTGGATTCTTTCGTAGCTGGCACTGATACATCTTCTAACACACTGGAATGGGCTATGGCA<br>GAACTACTAAAGAATCCGCATACATTAGAAAAAGTACAGGAGGAGTTAGCCCAGGTCATCGGACGTGGCAAGTTAATTAACGA<br>GGCTGATGTATCAAAGCTACCTTACCTGAAATGTATCGTTAAAGAGACTTTTAGGATTCACCCCCAGGTACCCTTCTTGATTCCCC |

| | |
|---|---|
| | GTAAGGCCGAAGAGGATGTCGAATTCTGTGGGTATATTATTCCTAAAGACAGTCAGGTTCTGGTAAACGTGTGGGCTATTGGCA<br>GGGATAGTAGTCTATGGGAGGATCCCTTAGACTTCAAGACAGAGCGTTTCTGGGAGTCAGAAATTGATGTGAGAGGGCAGGATT<br>TCGAGCTGATCCCTTTCGGGGCCGGAAGACGTATTTGTCCAGGCTTACCGTTGGCAATACGTATGATTTTAGTCACCTTAGGAAG<br>CCTGGTCAACACTTTCAATTGGAAATTGGATGGGGGCATTGCCCCCAAGGATTTGGATATGGAGGAAAAATTTGGTATTACTCTA<br>GCCAAAACCCAACCTCTACTAGCCATTCCAATACCCCTACAGCTTCTGGGATACTAA |
| *VvG10H* | ATGGATTACACTCCGCTTGTTCTATTGTTGTTGCTGCCTTGCTTTGTTTGGCTATGTTTTCATTTCCTGATCTTGGGATCAACACAC<br>AGAAAGTCATTCCAGGCGCGTCTACCGCCGGGCCCCAGACCTCTGCCTATTATCGGCAACTTGCTGGAGTTAGGGGATAAACCTC<br>ATCAATCCCTAACGACCCTTAGTAAGACTTACGGTCCTCTTATGTCCCTTAAGTTAGGTAGCACAACGACCATAGTGATCTCCAG<br>TCCGAAGACAGCACAAGAAGTGTTAAATAAGAAGGACCAAGCCTTTGCATCCAGAACAGTTTTAAACGCAATTCAAATACAGGA<br>TCATCACAAGTTCTCAATGGTGTTTTTGCCGGCGTCCGCGCATTGGCGTAACCTTAGGAAGATATGTTCTATGCAGATATTTTCAC<br>CCCAAAGGGTGGAAGCATCTCAAGACCTGAGAAGGAAGGTTGTTCAGCAACTTTTGGAACACGCCAGGGAAAGCTGCAACAGC<br>GGCAGGGCTGTGGACGTCGGACGTGCAGCTTTTACAACTACACTGAACTTATTATCTAATACTTTTTTTAGCGTTGATTTAGCTCA<br>CTACGATTCCAACTTAAGTCAAGAGTTTAAGGATCTTATTTGGTCAATAATGGTTGAGGCCGGAAAACCCAACCTAGCGGATTTT<br>TTTCCCGGCTTGAGGTTGGTAGACCCGCAGGGGATACAGAAACGTATGACCGTGTACTTTAACAAACTACTGGACGTATTTGACG<br>GATTTATTAATCAGCGTCTACCCCTGAAAGCATCAAGTCCTGACAACGATGTTTTGGATGCTCTGTTAAACTTGAACAAGCAGCA<br>TGACCATGAGCTTAGCTCCAATGACATAAGACACCTTTTAACCGATCTATTTTCCGCAGGCACCGACACCATATCTTCCACAATT<br>GAGTGGGCCATGGCCGAATTACTGAATAACCCGAAGGCAATGGCTAAAGCACAAGACGAATTGTCTCAGGTCGTCGGAAAAGA<br>TCGTATTGTAGAAGAATCTGATGTTACAAAATTACCGTATTTACAGGCGGTAGTCAAAGAAACCTTTAGACTTCACCCACCAGCA<br>CCCTTTCTGGTCCCGAGGAAGGCTGAAATGGACTCAGAAATCCTAGGGTATGCGGTTCCCAAAAACGCGCAGGTTTTGGTCAAT<br>GTTTGGGCCATCGGTAGGGATTCCAGGACGTGGAGCAATCCGAACTCTTTCGTGCCAGAGCGTTTTCTAGAGTGTCAAATTGACG<br>TTAAGGGTCGTGACTTTCAATTAATCCCGTTTGGCGCCGGTAGGAGGATCTGTCCAGGATTGCTTCTAGGACACCGTATGGTTCA<br>TCTGATGCTTGCAAGTTTACTTCACTCTTTCGACTGGAAATTAGAGGACTCCATGCGTCCCGAAGACATGGATATGTCCGAGAAA<br>TTCGGGTTTACGTTGCGTAAGGCTCAGCCACTGCGTGCCGTACCGACTAAACCCTAA |
| *PsCPR* | ATGGGGAGCAATAATCTTGCTAACTCTATAGAGTCTATGTTGGGTATATCCATCGGCTCTGAGTACATTTCTGACCCGATCTTTAT<br>TATGGTAACGACAGTAGCCTCAATGCTAATAGGATTCGGATTCTTCGCCTGCATGAAGTCATCTTCATCACAGTCCAAGCCTATA<br>GAGACGTATAAGCCGATAATAGACAAGGAGGAGGAAGAGATAGAGGTCGATCCTGGAAAGATAAAACTTACTATCTTTTTTGG<br>AACTCAAACGGGGACTGCCGAAGGCTTTGCCAAGGCGCTAGCGGAGGAAATCAAAGCCAAGTATAAGAAGGCGGTTGTCAAAG<br>TAGTTGACCTGGACGATTATGCCGCCGAGGATGATCAATATGAAGAGAAGCTTAAGAAGGAGAGTCTAGTCTTCTTTATGGTTG<br>CAACCTACGGCGACGGAGAGCCTACCGACAACGCCGCAAGATTCTATAAGTGGTTCACTCAAGAACACGAAAGGGGGGAGTGG<br>CTTCAGCAATTAACCTATGGAGTTTTTGGCTTAGGCAACCGTCAATACGAGCACTTTAACAAGATAGCCGTCGACGTCGACGAAC<br>AGCTAGGGAAGCAGGGGGCCAAGAGGATAGTGCAAGTAGGGTTGGGTGACGACGATCAATGCATTGAGGACGACTTTACGGCA<br>TGGCGTGAATTATTGTGGACAGAGCTTGATCAGCTGTTAAAAGATGAAGATGCGGCaCCGTCAGTAGCCACGCCGTACATAGCA<br>ACGGTCCCAGAGTATAGGGTTGTAATTCACGAAACAACCGTCGCAGCATTAGACGACAAACATATAAACACGGCTAACGGTGAT<br>GTTGCATTCGATATCCTGCACCCTTGTCGTACGATCGTCGCCCAACAAAGGGAGCTACACAAACCTAAGTCAGACAGATCCTGCA<br>TCCATCTTGAATTTGACATAAGTGGTAGCAGTCTTACGTATGAAACAGGCGATCACGTAGGAGTTTATGCCGAAAATTGCGACG<br>AGACAGTCGAAGAAGCTGGCAAGTTGCTTGGCCAGCCATTAGATCTTCTGTTCAGCATTCATACAGACAAGGAGGACGGAAGTC |

| | |
|---|---|
| | CACAAGGGTCTAGTCTACCGCCCCCCTTTCCTGGGCCCTGTACTTTGAGGTCTGCACTGGCCAGGTACGCCGATCTGCTTAATCC<br>GCCCCGTAAAGCGAGCCTGATTGCGCTTAGCGCCCATGCCTCTGTACCCTCTGAGGCTGAAAGGCTAAGGTTCTTAAGCTCACCT<br>TTGGGCAAGAACGAGTACTCTAAATGGGTTGTAGGTTCTCAACGTTCACTTCTAGAAATAATGGCTGAATTTCCCTCTGCAAAAC<br>CCCCGCTAGGGGTATTCTTCGCGGCGGTGGCCCCCAGATTACCCCCACGTTACTATTCAATATCTAGTAGTCCGAAGTTTGCTCCC<br>AGTAGAATACACGTGACGTGCGCCCTTGTATATGGACAGTCCCCGACGGGAAGGGTGCATCGTGGTGTATGCAGCACCTGGATG<br>AAGCATGCCGTACCGCAAGATAGTTGGGCTCCTATTTTCGTAAGAACAAGCAATTTCAAGCTGCCGGCCGACCCTAGTACCCCCA<br>TAATCATGGTCGGACCGGGGACCGGGCTTGCCCCCTTTAGAGGGTTTTTGCAGGAGAGGATGGCTTTGAAGGAGAACGGTGCGC<br>AATTGGGACCAGCGGTCTTGTTTTTTGGGTGCAGAAACAGGAATATGGACTTTATTTACGAGGACGAACTTAATAATTTCGTCGA<br>GCGTGGAGTCATAAGCGAACTGGTCATTGCCTTCTCTAGGGAGGGTGAGAAAAAGGAGTACGTTCAACACAAGATGATGGAGA<br>AGGCCACTGACGTGTGGAACGTAATATCCGGCGACGGTTACCTTTATGTCTGTGGCGATGCGAAAGGTATGGCCAGAGACGTAC<br>ATCGTACACTGCACACAATTGCGCAAGAGCAGGGACCGATGGAGTCTAGCGCCGCAGAAGCAGCTGTTAAGAAACTTCAAGTCG<br>AGGAGAGATACTTGAGAGATGTATGGGGCTGAAGAGC |
| *AtCPR* | ATGAGTTCATCCAGTTCTTCTTCAACCTCAATGATTGATCTAATGGCTGCCATAATTAAAGGCGAGCCGGTCATAGTGAGTGACC<br>CCGCTAATGCTAGTGCCTACGAATCCGTAGCGGCAGAATTATCCTCTATGTTAATTGAAAATAGACAGTTCGCGATGATCGTTAC<br>GACTTCTATCGCAGTACTTATAGGCTGCATAGTAATGTTAGTATGGAGGCGTAGCGGATCCGGGAACTCTAAGAGGGTCGAACC<br>TCTTAAACCGCTTGTGATAAAACCCAGAGAGGAGGAGATTGATGACGGGAGGAAGAAGGTCACGATTTTTTTCGGCACCCAGAC<br>AGGGACGGCTGAAGGTTTCGCGAAAGCCCTGGGGGAGGAAGCAAAAGCGAGATACGAAAAGACGAGATTTAAGATAGTTGACC<br>TTGATGATTATGCTGCCGATGACGATGAGTACGAAGAAAAACTTAAAAAGGAGGATGTTGCCTTCTTTTTCCTGGCAACTTACGG<br>CGATGGTGAGCCAACAGATAACGCCGCCCGTTTTTATAAGTGGTTTACCGAGGGTAATGATAGAGGCGAATGGTTGAAGAACTT<br>GAAATATGGGGTCTTTGGGCTGGGCAACAGACAGTATGAGCACTTTAACAAAGTAGCGAAGGTGGTTGATGATATCCTAGTAGA<br>GCAAGGAGCTCAAAGGCTTGTCCAAGTTGGCCTGGGGGATGATGACCAATGTATTGAAGACGACTTTACAGCATGGCGTGAAGC<br>TCTGTGGCCCGAATTAGATACCATATTGAGAGAAGAAGGTGACACGGCAGTGGCTACGCCTTATACGGCGGCCGTTCTGGAATA<br>CCGTGTATCCATCCACGATTCCGAAGATGCGAAATTCAACGATATTAATATGGCAAATGGGAATGGGTATACGGTGTTCGACGC<br>TCAGCACCCGTATAAGGCGAACGTCGCGGTAAAGCGTGAACTACATACTCCCGAGAGTGATAGGAGTTGCATCCACTTAGAATT<br>TGATATAGCGGGCTCCGGACTTACATACGAAACAGGGGACCACGTTGGTGTGTTATGCGATAACTTATCAGAGACTGTAGATGA<br>AGCGCTGCGTTTACTTGACATGTCCCCGGACACATACTTCTCTTTGCACGCCGAGAAAGAGGACGGGACTCCTATATCATCCAGC<br>TTACCGCCCCCGTTCCCTCCCTGTAATCTGAGAACTGCGCTTACTCGTTACGCGTGTCTATTATCCTCACCTAAAAAGTCCGCGCT<br>GGTCGCCTTAGCTGCTCATGCGTCAGACCCGACAGAGGCTGAACGTCTAAAACACCTGGCCTCACCaGCGGGTAAAGACGAGTA<br>CTCTAAGTGGGTGGTAGAAAGCCAACGTTCTCTGCTAGAAGTAATGGCGGAGTTCCCATCAGCCAAACCACCGCTAGGTGTATTT<br>TTCGCAGGGGTCGCCCCAAGACTGCAGCCTCGTTTCTACTCCATATCTAGCTCCCCTAAGATCGCGGAAACAAGGATACATGTTA<br>CATGTGCACTAGTATACGAAAAGATGCCTACGGGTAGGATACATAAGGGGGTCTGCTCTACATGGATGAAAAACGCGGTCCCAT<br>ATGAGAAGTCTGAAAATTGCTCAAGCGCACCAATCTTTGTGCGTCAATCCAATTTTAAATTACCCTCAGACAGTAAAGTACCTAT<br>TATTATGATAGGTCCAGGCACCGGACTGGCTCCATTCAGAGGCTTTTTACAGGAGAGGTTAGCCCTTGTAGAGTCTGGCGTCGAA<br>CTGGGACCAAGTGTCCTGTTCTTCGGCTGCAGAAATAGACGTATGGATTTTATATATGAAGAGGAGTTGCAAAGATTCGTAGAG<br>AGCGGGGCTCTAGCTGAACTATCCGTTGCATTCTCACGTGAAGGCCCAACGAAAGAATACGTGCAGCACAAGATGATGGATAAA<br>GCAAGTGACATATGGAATATGATATCCCAGGGTGCTTATTTGTACGTATGCGGGGATGCAAAGGGTATGGCCCGTGATGTACAT<br>AGGTCCCTGCACACGATTGCACAAGAGCAAGGTAGTATGGATTCCACTAAAGCTGAAGGGTTCGTGAAGAACTTACAGACATCA<br>GGACGTTACTTGAGGGACGTCTGGGGCTGAAGAGCTAG |

| | |
|---|---|
| *HcCPR* | ATGGAGAGCAGTAGCGTAAAGTTTAGCCCCTTGGACCTGATGTCCGCCATATTGAATGGCAAGAGTGATCCATTGGATGCTAGT<br>ATCCTAATTGAAAACCGTGAAGTCCTTATGATACTGACTACATCCATAGCAGTTCTAATAGGATGTGTAATACTGTTTCTTTGGA<br>AACGTTCCAGTGGCCAAAAATCAAGTAAGAGATTGGAACCACCGAAGCCGTTAATATTTAAAGAACCGGAGCCCGAGGTCGAC<br>GACGGCAAAAAAAAGGTCACGATCTTTTTCGGAACGCAGACTGGCACCGCTGAAGGTTTTGCTAAAGCCCTTGCGGAAGAGGCA<br>AAGGCCAGATACGAAAAGGCAATCTTTAAGGTTGTTGATTTGGATGATTACGCCGCAGACGATGATAAGTATGAAGAAAAGATG<br>AGCAAGGAAACCGTAGCCCTGTTCTTTATGGCGACATACGGGGATGGTGAACCAACGGATAACGCAGCGACTTTCTACAAGTGG<br>TTCACGGAGGGGAAAGAAAGAGGCATATGGCTACAGAATCTACAGTACGGGGTTTTTGGCTTGGGCAATAGGCAATATGAACAC<br>TTTAATAAGGTGGCCAAAGTTGTAGACGAGATTCTAGCCGAGCAGGGAGGAAAACGTCTAGTATCAGTCGGGCTTGGAGACGAT<br>GACCAATGCATAGAAGATGACTTTACTGCGTGGAGAGAATTAGTGTGGCCGGAGCTGGACAAGCTTCTGCTTGACGAGGACGCG<br>GCGACGTCTGTCAGTACGCCGTACACTGCCGCTGTCCCGGAATATCGTATAGTATTTCATGATCCAAGCGATGCCAGCTTACAGG<br>GAAAAAACCGTTCCAACGCCAATGGCCATACCATACCTGATATTCAACACCCCTTCCGTGTAAATGTCGCTGTAAAAAGGGAGC<br>TGCATACGCCCGCCAGCGATCGTAGTTGTACCCATCTTGAGTTTGATATTTCCAATACTGGTCTAACATACGAAACAGGAGATCA<br>CGTGGGTGTTTATAGCGAAAACTCCATCGAGTATGTGGAAGAGGCAGAGAGGTTGCTTGGTTATAGCGCTGATACTTTCTTTAGT<br>ATCCATGTTGAGGATGAAGACGGAACCCCCCTGGGAGGATCCTCACTAGCCCCACCATTCCCATCTCCCTGTACGCTGAGGACG<br>GCGCTTACAAGATATGCGGACTTACTGAACAGCCCGAAGAAAGCTGCGTTGATGGCCCTTGCCGCCCACGCGTCCGACCCCACG<br>GAGGCCGACAGACTACGTTTTTTGGCAAGTCCTGCCGGGAAGGACGAGTATGCTCAGTGGGTTGTAGCATCCCATCGTTCCTTAT<br>TGGAAATAATGGCAGAGTTTCCATCTGCCAAACCACCCTTGGGCGTATTCTTTGCCGCCATCGCCCCCAGGCTTCAACCGAGATT<br>TTATAGCATATCATCTTCTCCGCGTATCGCACCAACCAGGATCCACGTAACGTCCGCTATTGTATACGAAAAAACGCCTTCAGGG<br>AGAATCCACAAGGGAGTATGCAGCACGTGGATGAAGAACTCCGTTCCGCTAGAAGAAAGCCCGGATTGCAGCAGTGCGCCCAT<br>ATTCGTCAGGCAGAGTAACTTCAAATTACCAACTAACACGAGCCTACCAATCATTATGATTGGCCCTGGCACGGGGTTAGCTCCA<br>TTCCGTGGTTTTTTGCAAGAAAGGTTGGCGTTGAAGGAGGCGGGCACAGAGCTTGGGCACGCAATATTGTTCTTTGGCTGCAGGA<br>ACCGTAAGATGGACTACATTTATGAGGACGAGCTTAACGGGTTCGTGAAGGGGGGAGCGCTGACAGAGTTAGTAGTTGCCTTTA<br>GCAGGGAGGGACCTACCAAGGAATATGTGCAGCACAAGATGATGGAACGTGCTTCAGATATCTGGAACATGATTTCTCAGGGAG<br>GCTACATCTACGTTTGCGGAGACGCGAAAGGAATGGCTAAAGACGTTCACAGGACTCTTCACACGATAGTCCACGAAAAGGGCA<br>GTTTAGATAATTCTAAGACGGAGATGATGGTAAAGAACCTGCAAATGGACGGTAGGTATCTACGTGACGTATGGGGCTGAA |
| *NdCPR* | ATGCAAGAAGCGTCCAGCGTCAAGATTAGCCCCTTCGACTTGATGTCCGCCATCTTGCAAGGAAAATCCGACCCCCTGGATGTCA<br>CTGTATTGATCGAGAACCGTGAGGTCTTGATGATCCTTACGACTAGCATTGCGGTATTGATAGGGTGCGTGGTTCTGTTCGTCTG<br>GCGTAGATCCTCAGGGAGTAAGAGCAGCAAGAGCGTCGAAACTATCAGGCCACTTGTCCCGAAGGAACCGGAACCTGAGGCTG<br>ATGATGGCAAGAAGAAAGTGAGTATCTTCTTCGGGACACAGACTGGAACGGCGGAGGGATTCGCGAAGGCTTTAGCCGATGAA<br>GCCAAAGCTAGATATGACAAAGTGGCATTTAAGGTGATTGATCTGGACGACTACGCTGCTGATGACGCTGAGTACGAAGAAAAA<br>TTCAAGAAAGAATCTGTAGCTATATTCTTCCTGGCGACATATGGTGACGGGGAACCTACCGACAACTGTGCAAGGTTTTACAAGT<br>GGTTCACAGAGGAAGGAAAAGAGAAGGGCGCGTGGTTACAGAATCTGCAGTTTGGGGTTTTCGGACTTGGAAATAGGCAATAC<br>GAACATTTCAACAAGATAGCGAAGGTTGTCGATGAGCAGTTAGCGGAACAAGGGGCAAAACGTCTAGTACCTGTCGGCATCGGT<br>GATGATGATCAGTGTATCGAGGACGACTTTACCGCCTGGAGAGAGCTTGTGTGGCCTGAGCTAGATCAATTATTGCGTGACGAA<br>GACGATACAACTACGGTGAGCACTCCGTATACGGCAGCAGTCCTAGAATATCGTGTAGTATTCTACGACCCTGAAGATGCCCTTC<br>AGCAAGTGAAAAACTGGTCTAATGCCAACGGGCACGCAGTGCACGATATTAATCATCCATGCCGTGCAAATGTTGCTGTGAAGC |

| | |
|---|---|
| | GTGAGCTGCACACTCCAGCTAGCGACAGGTCATGTACTCATCTAGAGTTCGACATTAGCCATACTGGATTAACGTACGAAACGG<br>GGGATCATGTGGGCGTCTTCTGCGAGAATCTTATTGAGGACGTGGAAGAGGCTGAGAGGATACTTGGCTTAGCGCCGGACACCT<br>TTTTCAGCATACACACTGAAAATGAAGACGGCACCCCTGTAGGGAAGACTTCTTTGCCACCTCCATTCCCCTCCCCCTGCACTCTT<br>CGTGCTGCCCTTACGAGGTACGCAGATTTTCTAAACTCCCCAAAAAAGGCAGCaCTTCTTGCTTTAGCGGCGCACGCATCAGACC<br>CAAGCGAAGCCGATAGGCTAAGGTTTCTGGCATCACCGGCTGGGAAGGAGGAATACGCTCAGTGGATTGTCGCGTCACAACGTT<br>CACTTTTAGAGGTTATGGCTGCTTTCCCCAGCGTGAAGCCCCCCTTAGGCGTCTTCTTTGCAGCTATTGCACCCAGGTTACAACCC<br>CGTTACTACTCTATCTCATCAAGTCCGAGAATGGCtCCGAGTAGAATACACGTCACTTGCGCGCTTGTATATGAAAAAACTCCGA<br>CGGGTAGGATACATAAAGGTGTGTGTAGTACGTGGATGAAGAACGCTGTGCCACTAGAAGAGAGCAGGGACTGTTCCTGGGCTC<br>CTATTTTCGTCAGACAGAGCAACTTCAAACTTCCTATGGATACAACAACCCCGGTCATAATGATAGGCCCAGGGACCGGGCTGG<br>CACCCTTTCGTGGGTTCCTTCAGGAGAGGCTGGCCCTTAAGCAAGCAGGCGCTGAGCTGGGCCCTGCAATATTGTTTTTTGGGTG<br>CAGGAATAGGCGTACTGATTTCATTTACGAGGACGAACTTAATGGTTTCGTAGAGTCTGGTGCTCTTAGTGAGCTGATTGTTGCA<br>TTCTCCAGGGAGGGGCCAAATAAAGAATACGTACAACATAAAATGATGACGAGGGCGAGCGATGTATGGAACATGATCTCCCA<br>GGGCGGCTACCTGTATGTATGTGGAGATGCTAAGGGCATGGCGAAAGACGTGCACCGTACGTTGCATACGATAGCTCAAGAACA<br>AGGTAGTCTTGATAGTACCAAGGCGGAAGCGATGGTGAAAAATCTACAGACGGATGGGAGGTATCTGCGTGATGTTTGGGGCTG<br>AA |
| *NsCPR* | ATGGCAAGTAATCTTGTAAATTCTTTAGAGTCCGCGCTTGGAATTTCTTTGGGCTCAGAGAGCACCGACCCGCAGACCATTTTGG<br>TCGTATTAACAACCTCTGTAGCGGTCGTGATTGGACTGGTCGTTTTCTTCTGGAAAAGAAGTTCCCAGGATAGATCCAGTAAGCC<br>GATCCAAAGCTTAAAACCCATAAGTTTTAAAGCCGAGGAAGAGGATGAAATCGAACCTGGCAAGACTAAAGTAACAATTTTCTT<br>CGGTACTCAAACAGGGACTGCCGAGGGATTTGCCAAGGCCTTAAGCGAGGAAATAAAGGCGAGGTACGAAAAGGCTGCAGTCA<br>AGGTTGTCGATCTGGACGACTACGCGGCAGAAGACGATCAATATGAAGAGAAACTTAAGAAAGAAACATTGAGCTTCTTCATGT<br>TAGCCACCTATGGCGACGGGGAGCCGACGGACAACGCTGCCCGTTTTTATAAATGGTTCACTCAGGAGCATGAGAGAGGTGCAT<br>GGTTGCAACATATGTCTTATGCTGTGTTTGGACTAGGAAACAGACAGTACGAGCACTTCAATAAAATCGCAAAAGTCGTCGACG<br>AACAACTTATTGAGCAGGGTGCGAAGAGGCTGGCGGAAGTAGGCTTAGGTGACGATGACCAGTGCATTGAGGATGACTTCACCG<br>CATGGCGTGAAAGCCTATGGCCTGAATTGGACCAGTTATTAAGAGATGAGGACGATATCAATACCGTCACCACTCCCTATACAG<br>CAGCGGTGCCAGAGTATAGGGTTATGATACATGACCCCACAGTGACTAGCTGGGAAGATAAGTACCTGTCCATGTCAAATGGCA<br>ATGGTTATGCTAACGGCCATTCCGCATTCGATATCCACCACCCTTGTCGTGCTAACGTTGCGGTCACGAAGGAACTTCACAAACC<br>AGATTCCGATAGATCTTGCATCCATCTGGAATTTGATATAAGTGGGACCGGACTTATCTACGAGACtGGAGATCATGTAGGTGTT<br>TACGCGGAGAACTGCGATGAGACAGTAGAGGAGGCTGGCAAACTGCTTGGCCAGTCCCTAGACCTGTTGTTCTCACTTCATACC<br>GATAAAGAGGATGGCACCCCTCAAGGATCCTCCCTGCCGCCCCCGTTCCCAGGACCATGTACGCTTAGGGTCGCTCTTGCTCGTT<br>ACGCTGATTTGCTGAATCCTCCCAAGAAAGCCGCGCTGACCGCATTGGCTGCCCACGCATCTGAACCCAGTGAGGCTGACAAGC<br>TAAAATATTTGTCCTCACCTGAAGGTAAGGACGAGTATTCAAAGTGGATCGTTGGATCTCAAAGGGGACTTTTGGAGGTGATGG<br>CAGAGTTTCCGTCAGCCAAACCTCCAGTCGGAGTATTTTTCGCGGCCATAGCGCCACGTCTACAGCCCAGGTACTACTCAATTTC<br>TTCCAGCCCCAGGTTTTCTCCTTTTAGAATACACGTCACtTGCGCACTTGTGTACGGACCCAGTCCAACTGGCAGAATCCATAAGG<br>GGGTCTGTTCCACTTGGATGAAAAACGCTGTTCCGCTAGAAAAAAATCACGATTCTTCATGGGCtCCCATCTTCGTCAGGACAAG<br>TAACTTTAAACTTCCTGCAGACCCATCTATTCCTATCGTGATGGTGGGCCCTGGGACAGGCCTGGCTCCCTTCAGAGGCTTCCTTC<br>AAGAGAGGATGGCACTTAAGGAGGACGGTGTTCAGCTTGGACCTGCTGTACTATTTTTTGGCTGTAGGAATAGACGTATGGACTT<br>CATATACGAGGACGAACTTAACAGTTTCGTCGAACGTGGAGCCTTAAGCGAATTGGTAATCGCCTTTAGTAGGGAGGGGCAACA<br>AAAAGAGTACGTACAACACAAAATGATGGAAAAAGCCGCGTATATCTGGAGCTTAATAAGTAATGGCGGCTACCTATACGTTTG |

| | |
|---|---|
| | TGGGGATGCGAAAGGCATGGCCAGAGATGTACACAGGACACTTCACACTATAGTGCAGGAACAAGATCACGTAGACTCAACGA<br>AAGCTGAATCATCAGTGAAGAAGTTGCAGATGGATGGCAGGTACCTTAGGGACGTGTGGGGCTGAA |
| *PkCPR* | ATGCAATCCACGCCGGAAAAATTATCTCCATTAGACCTTATGACTGCCATATTGAAAGGGGTCAATCTGGACACCTTGAATATAA<br>GCTCCGAGTCAGCCCCGGCTCCGGTGGTTGCGATGTTATTAGAATACCGTGAGCTTCTGATGATGGTGACTACGTCAATCGCAGT<br>ACTGATTGGGTGCGCCGTCTTCTTAGCGTTGCGTGGTGGTAAGAAAAAGAAAAAGGCAGTAGAACCCCCGAAATTGGTGGTACC<br>CAAGGGACCTGTGGAGCCTGAGGAAGTTGATGACGGGAAGAAAAAAGTGACCATCTTTTTCGGAACGCAGACCGGCACGGCAG<br>AAGGTTTTGCGAAGGCCCTGGCGGAAGAGGCTAAGGCGCGTTATCAACAGGCGAAATTCAAAGTAGTGGACTTAGATGATTATG<br>CCGCGGACGATGAGGAATACGAGGAGAAAATGAAAAAGGAAACCTTGGCCTTTTTCTTCCTGGCGACCTACGGAGACGGTGAG<br>CCAACGGATAATGCCGCCAGATTCTATAAGTGGTTCACTGAGGGAAAGGAGAAAGGGGACTGGCTTAAAAATCTGCAATATGGC<br>ATTTTCGGGCTGGGGAATAGGCAGTACGAGCACTTTAATAAGATTGCCCTAGTGGTGGATGACATTATCACTGGGCAAGGAGGA<br>AAGAGGTTAGTCCCTGTAGGCCTAGGTGACGACGACCAGTGTATCGAAGATGACTTCACTGCGTGGAGGGAATTAGTGTGGCCA<br>GAGCTTGACAAACTTTTGTTAAACGAGGATGAAGCCTCAGCCGCAACCCCGTACACCGCTGCAGTGCTGGAATACAGAGTAGTG<br>TTCCATGATCAAGTCAACGAGTCCTCTCTAGAAAACGGTTTAGCCAATGGCCACGCGAATGGTCATGCAGTTTATGATGCCCAGC<br>ACCCCTGCAGAGCCAATGTGGCAGTGAGGAAAGAGCTGCACGCCCCTGCCAGTGACAGATCATGCACCCACCTAGAGTTCGACA<br>TCGCTGCTACAGGGCTTATGTACGAGACTGGTGACCATGTTGGTGTTTACTGCGAAAATCTAATCGAGAACGTTGAGGAGGCTG<br>AGAGGCTTTTGAACATGCCTCCGCAGACGTATTTTTCCATTCATACCGACAAAGAGGATGGAACCCCACTATCAGGATCAAGCCT<br>ACAACCACCATTCCCTCCCTGTACATTAAGAACAGCCTTAACAAGATACGCGGACCTACTAAGCGCTCCAAAAAAGAGTACTCT<br>AGTTGCCTTGGCGGCATATGCTAGTGACTTAAATGAAGCTGATAGGCTGAGACATTTGGCATCTCCCGTCGGCAAGGAAGAGTA<br>TACTCAATATATACTAACGAATATGAAATCCTTACTGGAAGTCATGGCAGACTTTCCTTCCGCCAAACCGCCTCTTGGGGTATTC<br>TTTGCCGGTGTTGCGCCCAGGCTTCAACCCCGTTTTTGTAGCATCTCCTCTTCACCGAAGATAGCTCCGAACAGAATTCACGTGAC<br>TTGCGCGCTTGTTTACGAAAAGACCCCAACGGGCCGTATACATAAAGGAATTTGTAGCACATGGATGAAAAACGCGATCCCAAG<br>TGAAGAGAGTTTGGACTGCTCAAGTGCTCCAATCTTTGTCAGAACGTCAAATTTCAGACTTCCCGCAGACCCGAAGGTACCGATA<br>ATAATGATTGGCCCAGGGACTGGTTTGGCACCTTTCAGAGGCTTTTTGCAAGAAAGATTAGCATTGAAAGAAAGTGGAGCTGAA<br>TTGGGGCCAGCGGTGCTTTTTTTTGGGTGTAGAAATCGTCAGATGGACTTTATATACGAAGACGAGCTAAATAACTTTGTTAAAG<br>CAGGGGTGATCAGCGAGTTGGTTTTAGCATTCTCCAGGCAAGGACCTACTAAGGAATACGTGCAACATAAAATGGCTCAAAAAG<br>CCAGTGATATATGGAACATGATCTCAGAGGGCGGTTACGTGTACGTGTGTGGTGACGCGAAAGGTATGGCGAGAGATGTACACC<br>GTACACTGCACACCATCGTACAGGAACAGGGGAGCTTAGATAGCTCCAAAACTGAAGCTCTTGTTAAGAACCTACAAATGACAG<br>GAAGGTATTTAAGAGACGTTTGGTAA |
| *Sr1CPR* | ATGCAATCAGATTCAGTCAAAGTCTCTCCATTTGATTTGGTTTCCGCTGCTATGAATGGCAAGGCAATGGAAAAGTTGAACGCTA<br>GTGAATCTGAAGATCCAACAACATTGCCTGCACTAAAGATGCTAGTTGAAAATAGAGAATTGTTGACACTGTTCACAACTTCCTT<br>CGCAGTTCTTATTGGGTGTCTTGTATTTCTAATGTGGAGACGTTCATCCTCTAAAAAGCTGGTACAAGATCCAGTTCCACAAGTTA<br>TCGTTGTAAAGAAGAAAGAGAAGGAGTCAGAGGTTGATGACGGGAAAAAGAAAGTTTCTATTTTCTACGGCACACAAACAGGA<br>ACTGCCGAAGGTTTTGCTAAAGCATTAGTCGAGGAAGCAAAAGTGAGATATGAAAAGACCTCTTTCAAGGTTATCGATCTAGAT<br>GACTACGCTGCAGATGATGATGAATATGAGGAAAAACTGAAAAAGGAATCCTTAGCCTTCTTCTTCTTGGCCACATACGGTGAT<br>GGTGAACCTACTGATAATGCTGCTAACTTCTACAAGTGGTTCACAGAAGGCGACGATAAAGGTGAATGGCTGAAAAAGTTACAA<br>TACGGAGTATTTGGTTTAGGTAACAGACAATATGAACATTTCAACAAGATCGCTATTGTAGTTGATGATAAACTTACTGAAATGG |

| | |
|---|---|
| | GAGCCAAAAGATTAGTACCAGTAGGATTAGGGGATGATGATCAGTGTATAGAAGATGACTTCACCGCCTGGAAGGAATTGGTAT<br>GGCCAGAATTGGATCAACTTTTAAGGGACGAAGATGATACTTCTGTGACTACCCCATACACTGCAGCCGTATTGGAGTACAGAG<br>TGGTTTACCATGATAAACCAGCAGACTCATATGCTGAAGATCAAACCCATACAAACGGTCATGTTGTTCATGATGCACAGCATCC<br>TTCAAGATCTAATGTGGCTTTCAAAAAGGAACTACACACCTCTCAATCAGATAGGTCTTGTACTCACTTAGAATTCGATATTTCTC<br>ACACAGGACTGTCTTACGAAACTGGCGATCACGTTGGCGTTTATTCCGAGAACTTGTCCGAAGTTGTCGATGAAGCACTAAAACT<br>GTTAGGGTTATCACCAGACACATACTTCTCAGTCCATGCTGATAAGGAGGATGGGACACCTATCGGTGGTGCTTCACTACCACCA<br>CCTTTTCCTCCTTGCACATTGAGAGACGCTCTAACCAGATACGCAGATGTCTTATCCTCACCTAAAAAGGTAGCTTTGCTGGCATT<br>GGCTGCTCATGCTAGTGATCCTAGTGAAGCCGATAGGTTAAAGTTCCTGGCTTCACCAGCCGGAAAAGATGAATATGCACAATG<br>GATCGTCGCCAACCAACGTTCTTTGCTAGAAGTGATGCAAAGTTTTCCATCTGCCAAGCCTCCATTAGGTGTGTTCTTCGCAGCA<br>GTAGCTCCACGTTTACAACCAAGATACTACTCTATCAGTTCATCTCCTAAGATGTCTCCTAACAGAATACATGTTACATGTGCTTT<br>GGTGTACGAGACTACTCCAGCAGGCAGAATTCACAGAGGATTGTGTTCAACCTGGATGAAAAATGCTGTCCCTTTAACAGAGTC<br>ACCTGATTGCTCTCAAGCATCCATTTTCGTTAGAACATCAAATTTCAGACTTCCAGTGGATCCAAAAGTTCCAGTCATTATGATA<br>GGACCAGGCACTGGTCTTGCCCCATTCAGGGGCTTTCTTCAAGAGAGATTGGCCTTGAAGGAATCTGGTACAGAATTGGGTTCTT<br>CTATCTTTTTCTTTGGTTGCCGTAATAGAAAAGTTGACTTTATCTACGAGGACGAGCTTAACAATTTTGTTGAGACAGGAGCATTG<br>TCAGAATTGATCGTCGCATTTTCAAGAGAAGGGACTGCCAAAGAGTACGTTCAGCACAAGATGAGTCAAAAAGCCTCCGATATA<br>TGGAAACTTCTAAGTGAAGGTGCCTATCTTTATGTCTGTGGCGATGCAAAGGGCATGGCCAAGGATGTCCATAGAACTCTGCATA<br>CAATTGTTCAGGAACAAGGGAGTCTGGATTCTTCCAAGGCTGAATTGTACGTCAAAAACTTACAGATGTCTGGAAGATACTTAA<br>GAGATGTTTGGTAG |
| *Sr8CPR* | ATGCAATCTAACTCCGTGAAGATTTCGCCGCTTGATCTGGTAACTGCGCTGTTTAGCGGCAAGGTTTTGGACACATCGAACGCAT<br>CGGAATCGGGAGAATCTGCTATGCTGCCGACTATAGCGATGATTATGGAGAATCGTGAGCTGTTGATGATACTCACAACGTCGG<br>TTGCTGTATTGATCGGATGCGTTGTCGTTTTGGTGTGGCGGAGATCGTCTACGAAGAAGTCGGCGTTGGAGCCACCGGTGATTGT<br>GGTTCCGAAGAGAGTGCAAGAGGAGGAAGTTGATGATGGTAAGAAGAAAGTTACGGTTTTCTTCGGCACCCAAACTGGAACAG<br>CTGAAGGCTTCGCTAAGGCACTTGTTGAGGAAGCTAAAGCTCGATATGAAAAGGCTGTCTTTAAAGTAATTGATTTGGATGATTA<br>TGCTGCTGATGACGATGAGTATGAGGAGAAACTAAAGAAAGAATCTTTGGCCTTTTTCTTTTTGGCTACGTATGGAGATGGTGAG<br>CCAACAGATAATGCTGCCAGATTTTATAAATGGTTTACTGAGGGAGATGCGAAAGGAGAATGGCTTAATAAGCTTCAATATGGA<br>GTATTTGGTTTGGGTAACAGACAATATGAACATTTTAACAAGATCGCAAAAGTGGTTGATGATGGTCTTGTAGAACAGGGTGCA<br>AAGCGTCTTGTTCCTGTTGGACTTGGAGATGATGATCAATGTATTGAAGATGACTTCACCGCATGGAAAGAGTTAGTATGGCCGG<br>AGTTGGATCAATTACTTCGTGATGAGGATGACACAACTGTTGCTACTCCATACACAGCTGCTGTTGCAGAATATCGCGTTGTTTTT<br>CATGAAAAACCAGACGCGCTTTCTGAAGATTATAGTTATACAAATGGCCATGCTGTTCATGATGCTCAACATCCATGCAGATCCA<br>ACGTGGCTGTCAAAAAGGAACTTCATAGTCCTGAATCTGACCGGTCTTGCACTCATCTTGAATTTGACATCTCGAACACCGGACT<br>ATCATATGAAACTGGGGACCATGTTGGAGTTTACTGTGAAAACTTGAGTGAAGTTGTGAATGATGCTGAAAGATTAGTAGGATT<br>ACCACCAGACACTTACTCCTCCATCCACACTGATAGTGAAGACGGGTCGCCACTTGGCGGAGCCTCATTGCCGCCTCCTTTCCCG<br>CCATGCACTTTAAGGAAAGCATTGACGTGTTATGCTGATGTTTTGAGTTCTCCCAAGAAGTCGGCTTTGCTTGCACTAGCTGCTCA<br>TGCCACCGATCCCAGTGAAGCTGATAGATTGAAATTTCTTGCATCCCCCGCCGGAAAGGATGAATATTCTCAATGGATAGTTGCA<br>AGCCAAAGAAGTCTCCTTGAAGTCATGGAAGCATTCCCGTCAGCTAAGCCTTCACTTGGTGTTTTCTTTGCATCTGTTGCCCCGCG<br>CTTACAACCAAGATACTACTCTATTTCTTCCTCACCCAAGATGGCACCGGATAGGATTCATGTTACATGTGCATTAGTCTATGAG<br>AAAACACCTGCAGGCCGCATCCACAAAGGAGTTTGTTCAACTTGGATGAAGAACGCAGTGCCTATGACCGAGAGTCAAGATTGC<br>AGTTGGGCCCCAATATACGTCCGAACATCCAATTTCAGACTACCATCTGACCCTAAGGTCCCGGTTATCATGATTGGACCTGGCA |

| | |
|---|---|
| | CTGGTTTGGCTCCTTTTAGAGGTTTCCTTCAAGAGCGGTTAGCTTTAAAGGAAGCCGGAACTGACCTCGGTTTATCCATTTTATTC<br>TTCGGATGTAGGAATCGCAAAGTGGATTTCATATATGAAAACGAGCTTAACAACTTTGTGGAGACTGGTGCTCTTTCTGAGCTTA<br>TTGTTGCTTTCTCCCGTGAAGGCCCGACTAAGGAATATGTGCAACACAAGATGAGTGAGAAGGCTTCGGATATCTGGAACTTGCT<br>TTCTGAAGGAGCATATTTATACGTATGTGGTGATGCCAAAGGCATGGCCAAAGATGTACATCGAACCCTCCACACAATTGTGCA<br>AGAACAGGGATCTCTTGACTCGTCAAAGGCAGAACTCTACGTGAAGAATCTACAAATGTCAGGAAGATACCTCCGTGACGTTTG<br>GTAA |
| *TcCPR* | ATGCAGACTTCCGAAGTCAAAATATCCCCTTTTGACCTGATGTCTGCTATCCTAAAAGGTTCCGTGGATCCTCTAGATTCCTCTGT<br>GTTAGTAGAAAACAGAGGCGTGTTGATGATGTTGACAACATCAATCGCTGTTCTTATTGGATGCGTTTTTATTTTTGTGTGGAGG<br>AGGAGCAGTGGGCAAAAAACTACGAAATCTGTGGAACCACCAAAACCTCTGATCGTTAAGGATGTTGAACCCGATGTGGACGAT<br>GGTAAAAAAAAAGTAACAATACTGTTTGGAACTCAGACGGGGACGGCGGAGGGTTTTGCGAAAGCGTTAGCCGAGGAAGCTAA<br>AGCCCGTTACGACAAAGCTACGTTTAAGGTAATAGACCTTGATGATTACGGGGCAGACGATGATGACTACGAGGAGAAACTGAA<br>AAAGGAGACAATAGTTTTTTcTTTCTTGCAACATACGGTGACGGTGAACCGACTGATAATGCGGCCAGGTTTTACAAGTGGTTC<br>ACTGAGGGCAAAGAAAGGGGTATTTGGCTGGACAAGGTACAGTTCGGAGTGTTCGGTCTAGGCAACAGACAGTACGAGCACTTT<br>AATAAGGTGGCAAAGGTCGTCGATGAGATTCTAGCCGACCAGGGTGGGAAGAGGTTAGTTCCGGTTGGCCTTGGCGACGATGAT<br>CAATGTATAGAAGACGACTTTACAGCTTGGCGTGAACTAGTATGGCCGGAATTAGATCAGCTTCTACAGGATGAGGATGGAATC<br>GCACATGTATCCACTCCGTACACCGCCGTTGTGCCCGAGTACAGGGTCGTGTTTCACGAAATAAGTAACGGACAGGTATACAAC<br>AAAGATCTTTCCAGTGCAAATGGTCATGCAGTTCACGACATTCAACATCCCTGTAGAGCGAACGTCGCCGTACGTAGAGAACTA<br>CACACACCGGCATCTGACAGATCCTGTACACACCTAGAGTTCGATATAAGCGGGACAGGACTTGTCTACGAAACCGGGGATCAC<br>GTAGGTGTATACTGCGAGAACTGCATAGAGGTAGTTGAGGAGGCCGAAAGGCTTTTAGGATATAGCGCCGACACCTTCTTCTCT<br>ATCCACGCGGATAACGAAGATGGTACGCCTCTATCTGGTTCTTCACTGCCTCCACCCTTCCCGTCCCCTTGCACACTGAGGATGG<br>CCCTTACGAGATACGCAGATCTATTGAATTTCCCTAAGAAAGCCGCATTGTTAGCATTAGCCGCGCATGCCTCTGACTCTTCAGA<br>AGCGGACCGTTTGAAGTTCTTGGCATCACCaGCCGGGAAGGACGAATACGCACAATGGGTGGTTGCGTCTCAGAGAAGCCTTTTA<br>GAAGTTATGGCAGAGTTTCCTAGTGTAAAACCGCCCCTAGGGGTGTTCTTCGCCGCGATCGCACCGCGTCTACAGCCTAGATACT<br>ATTCAATTAGTAGCAGCCCAAGAATGGCACCGAGTAGGATTCACGTTACATGTGCATTGGTGTATGAGAAAACGCCGGCGGGAA<br>GAATTCACAAGGGCGTCTGTTCAACTTGGATGAAGAACGCCGTCTCCCTTGAAGAATCTACGGACTGCTCCTGGGCTCCGATTTA<br>TGTACGTCAATCTAACTTTAAACTACCTGCGGATACTAGTGTTCCAGTCATCATGATAGGGCCTGGGACCGGCCTTGCACCCTTT<br>AGAGGCTTCTTACAGGAGAGGTTAGCCCTAAAAGAAGCGGGGGCAAATTTGGGTCCTGCCATCTTGTTCTTCGGCTGTAGAAAC<br>AGGAGGATGGATTATATCTATGAGGATGAGTTGAACGGCTTCGTAAAAGCAGGGGCTCTAACCGAACTAATAGTTGCTTTCTCTC<br>GTGAGGGGCCGACGAAGGAGTATGTTCAGCATAAGATGACCGAGAGAGCGTCCGACATCTGGTCCATGATATCTAAGGGTGCAT<br>ATCTTTATGTCTGTGGTGATGCAAAAGGAATGGCGAAGGACGTCCACCGTACTCTGCATACTATTGTGCAGGAACAGGGGTCaCT<br>AGATTCCTCAAAAGCCGAATCAATGGTGAAGTCCCTACAGATGGACGGACGTTACCTTAGAGACGTATGGGGCTGAA |
| *VvCPR* | ATGCAATCCTCCTCTGTTAAGGTGAGCCCATTTGACCTAATGAGCGCGATTATTAAGGGTTCCATGGACCAGTCAAACGTTTCCA<br>GTGAAAGCGGAGGTGCCGCAGCAATGGTCCTTGAGAACCGTGAATTTATTATGATCTTAACAACAAGCATTGCCGTTCTGATCG<br>GATGCGTTGTAGTGCTAATATGGAGGCGTTCCGGGCAAAAGCAGAGTAAAACTCCTGAGCCTCCCAAGCCCCTAATCGTTAAAG<br>ATCTTGAGGTGGAGGTCGACGATGGTAAACAGAAAGTCACAATCTTTTTTGGTACGCAGACGGGTACCGCAGAGGGATTCGCGA<br>AAGCCTTGGCGGAAGAAGCCAAGGCGAGGTATGAGAAGGCTATATTTAAGGTGGTTGACCTTGATGACTACGCTGGAGATGATG<br>ATGAGTATGAAGAGAAATTAAAAAAGGAGACTTTAGCGTTTTTTTTTCTTGCGACGTACGGAGATGGGGAGCCAACCGATAACG |

| | |
|---|---|
| | CGGCCCGTTTTTACAAATGGTTTGCAGAGGGTAAGGAGAGAGGAGAGTGGCTGCAAAATCTGAAATACGGTGTATTCGGCCTTG<br>GCAATAGACAATACGAACACTTTAACAAAGTAGCAAAGGTGGTGGACGACATTATCACTGAGCAAGGTGGTAAACGTATCGTAC<br>CGGTGGGACTGGGCGACGACGATCAGTGCATCGAGGACGACTTTGCCGCATGGAGGGAACTACTGTGGCCTGAACTAGACCAGT<br>TGTTGCGTGATGAGGACGACGCGACCACCGTATCTACTCCCTATACCGCAGCTGTATTGGAATATCGTGTAGTCTTCCATGACCC<br>TGAAGGCGCTAGCTTGCAAGACAAAAGTTGGGGGTCCGCTAATGGCCATACAGTGCACGACGCGCAACACCCGTGCAGGGCCA<br>ATGTTGCGGTACGTAAGGAGCTACATACCCCAGCTTCCGACAGGAGTTGTACCCACCTGGAATTCGACATCAGCGGAACTGGCC<br>TGACGTATGAAACAGGAGATCATGTGGGAGTTTACTGTGAGAACCTACCAGAGACAGTTGAAGAGGCCGAAAGACTTCTAGGAT<br>TCAGCCCGGATGTATATTTCTCAATACACACCGAGAGGGAAGACGGGACCCCGCTAAGTGGATCTAGTTTGAGCCCGCCTTTTCC<br>ACCGTGTACCTTGAGGACCGCGCTAACGAGATATGCTGATGTTTTATCTTCCCCCAAGAAGAGCGCCTTAGTTGCGTTAGCAGCA<br>CACGCATCCGACCCATCAGAAGCGGACCGTCTGAAGTACTTAGCGTCCCCAAGTGGGAAAGATGAATATGCACAATGGGTTGTC<br>GCCTCCCAAAGGTCACTACTGGAAATAATGGCGGAATTCCCTTCTGCCAAGCCACCGCTAGGGGTGTTTTTTGCAGCTGTGGCAC<br>CGAGATTACAGCCAAGATATTATAGTATATCCTCATCCCCTAAGATGGTGCCCTCCAGAATTCACGTAACATGCGCCCTAGTGTG<br>TGATAAGATGCCCACAGGCAGGATACATAAAGGAATTTGCAGCACTTGGATGAAATATGCGGTGCCTCTGGAGGAGTCCCAAGA<br>CTGTAGCTGGGCCCCTATTTTCGTAAGGCAGAGCAACTTCAAGCTTCCTGCTGACACAAGCGTCCCCATTATAATGATCGGACCC<br>GGTACAGGGCTGGCCCCCTTTAGAGGCTTCCTTCAAGAAAGGTTTGCGTTGAAGGAAGCGGGCGCAGAGCTGGGTTCTTCAATA<br>CTTTTCTTTGGATGCCGTAATAGAAAGATGGATTACATATATGAGGATGAACTGAATGGCTTTGTCGAGTCAGGCGCATTGAGTG<br>AGCTGATCGTCGCATTCAGTCGTGAAGGTCCCACCAAAGAATATGTTCAGCACAAAATGATGGAAAAGGCTAGTGACATCTGGA<br>ATGTTATCTCCCAGGGCGGATACATTTACGTCTGCGGCGACGCCAAAGGAATGGCAAGGGATGTCCACAGAACTCTTCATACAA<br>TTCTTCAGGAGCAAGGATCCTTAGATTCTAGCAAGGCGGAAAGCATGGTTAAGAACCTTCAGATGACCGGGAGATACTTGCGTG<br>ATGTATGGTAA |
| 10HGO | ATGACAAAGACAAACTCCCCCGCTCCCTCCGTAATCACTTGCAAAGCCGCAGTTGTCTGGAAGAGTGGCGAACCTCCAAAAGTT<br>GAAGAGATTCAGGTCGATCCACCCAAGGCTTCTGAGGTAAGAATCAAGATGTTGTGTGCCTCATTGTGCCACACCGATTTCTTGG<br>CGTGTAATGGATTGCCCGTGCCATTATTTCCCAGAATACCGGGTCATGAAGGAGTCGGGATGATAGAATCAGTGGGCGAGAATG<br>TGACAAACCTGAAGGAGGGCGATATTGTAATGCCGTTGTACCTTGGCGAGTGTGGTGAATGCCTAAACTGTAAAAGTGGTAGGA<br>CGAACCTATGCCACAAGTATCCGTTAGGCTTTAGTGGACTGCTGTTAGATGGAACGTCCCGTATGAGTATCGGAGAACAAAAAG<br>TATACCACCACTTCTCTTGTTCAACATGGAGCGAATATATCGTCATTGAGGCAGCTTACGCCGTCAAGGTAGACCCGAGGGTTTC<br>TCTTCCACACGCAAGCTTTCTATGCTGCGGGTTTACCACCGGTTTTGGCGCAACTTGGAGAGATGTGAACGTCGTCAAGGGAAGC<br>ACTGTTGCGGTATTGGGTCTTGGAGCAGTTGGACTTGGGGCAGTTCAGGGGGCAAAGTCACAGGGTGCAAGTAGGATCATAGGC<br>TTGGATATTAATGACAAGAAGAGGGAAAAGGGCGAAGCATTCGGTATGACGGAATTTATTAACCCTAAAGGGAGCAACAAAAG<br>CATCTCAGAATTAATAAACGAAGCTACGGGTGGGTTGGGCCTTGACTATGTGTACGAGTGCACTGGTGTACCGGCATTACTGAA<br>CGAGGCTATTGAGAGTAGCAAGGTGGGATTAGGGACCGCAGTGCTAATCGGAGCAGGGCTGGAGACTTCAGGAGAAATTAAGT<br>TCATACCTCTGTTATGCGGCAGGACGGTGAAAGGAAGTATTTACGGGGGAGTGAGGCCTAAAAGTGACCTACCAACATTGATCG<br>AAAAGTGTATCAACAAAGAAATTCCAATGGATGAGCTTATGACCCATGAAGTGTCTCTGTCTGAAATCAACAAGGGCTTTGAAT<br>ACCTTAAGCACCCGGACTGTGTGAAGGTCGTGATAAAGTTCTAG |
| ISY | ATGTCTTGGTGGTGGAAACGTAGTATCGGGGCGGGGAAGAATCTGCCTAATCAGAACAAAGAAAATGGCGTGTGTAAGTCTTAT<br>AAGTCCGTTGCACTTGTTGTTGGTGTTACCGGTATCGTTGGGAGCAGTCTAGCGGAGGTGCTGAAATTACCAGATACACCGGGGG |

| | |
|---|---|
| | GACCGTGGAAGGTGTACGGCGTAGCGAGAAGGCCGTGCCCCGTTTGGCTTGCCAAAAAACCTGTGGAGTATATCCAATGTGACG<br>TTAGCGACAATCAAGAAACGATTTCCAAACTGTCACCTTTAAAGGACATAACGCACATTTTTTACGTATCCTGGATCGGAAGTGA<br>AGATTGCCAGACCAACGCAACTATGTTCAAGAACATCCTGAATTCAGTAATCCCGAATGCGAGTAACCTGCAGCACGTATGCCT<br>ACAAACAGGAATCAAACACTACTTTGGAATATTTGAGGAGGGTAGTAAAGTTGTCCCACATGATAGCCCATTTACTGAGGACTT<br>GCCAAGACTGAATGTACCCAATTTCTATCACGACCTGGAGGACATATTATATGAAGAAACCGGTAAAAATAATCTAACATGGAG<br>TGTCCATCGTCCCGCACTTGTCTTTGGGTTCTCACCATGCTCTATGATGAACATTGTATCCACACTTTGCGTTTATGCGACGATCT<br>GCAAACACGAAAATAAGGCATTGGTTTATCCTGGGTCAAAAAACTCCTGGAACTGCTATGCAGATGCCGTTGACGCCGATCTAG<br>TAGCCGAACACGAGATATGGGCAGCTGTTGACCCTAAGGCGAAAAATCAGGTGTTGAATTGTAATAATGGCGATGTGTTTAAAT<br>GGAAGCATATCTGGAAGAAGTTAGCTGAGGAATTCGGAATTGAAATGGTTGGATATGTCGAAGGTAAAGAGCAGGTGTCTCTAG<br>CCGAATTAATGAAAGATAAGGATCAGGTTTGGGACGAAATCGTCAAGAAGAATAACTTGGTGCCCACAAAGCTTAAGGAAATA<br>GCCGCCTTTTGGTTCGCTGACATCGCCTTTTGTTCTGAGAATCTGATTTCATCCATGAATAAGTCCAAAGAATTGGGTTTCCTGGG<br>TTTTAGAAATAGCATGAAGTCTTTCGTATCTTGTATTGACAAAATGCGTGACTATAGGTTCATACCCTAA |
| NEPS1 | ATGGCGTCCACGGCAAATCCTATGCAGGTTATGAAAAAGAAGCTGGAGGGGAAAGTTGTCATCGTTACCGGGGGCGCCTCTGGT<br>ATTGGGCAAACGGCTGCTAGGGTGTTCGCTCAACACGGTGCTAGAGCAGTGGTGATCGCGGATATACAGAGTGAAGTTGGAAAA<br>TCAGTAGCTAAGAGCATTGGTGATCCGTGCTGTTACGTCCAATGCGACGTATCAGATGAGGAGGAAGTGAAGTCCATGATCGAG<br>TGGACCGCAAGCGCCTATGGAGGTCTGGACATGATGTTCTCTAACGTGGGGATTATGTCAAAGAGTGCTCAAACGGTGATGGAT<br>CTTGATTTACTTGAATTTGACAAAGTCATGAGGGTCAACGCCCGTGGGATGGCAGCCTGCCTAAAACACGCCGCTAGGAAGATG<br>GTCGAACTGGGGACCAGGGGCACCATAATTTGCACCACAACACCCCTTAGTAGCCGTGGTGGGCAAAGTATGACTGACTACGCC<br>ATGAGTAAGCACGCAGTCATGGGATTAGTACGTTCTGCTTCCATACAACTTGGTGCCCATGGCATTCGTGTTAACTGCGTAACTC<br>CTTCAGTGGTGCTTACGCCACTTGCCCAGAGAATGGGCTTAGCAACTCCGGACGATTTTCACACACATTTCGGTAACTTTACTAG<br>CCTAAAAGGCGTATACCTGACGCCGGAACAAGTTGCAGAGGCCGTGGTGTACCTAGCAAGCGACGATGCTGCCTTTATAACGGG<br>TCACGACTTGGTATTGGACGGTGGGCTATTATGCTTACCCTTTTTCGCTCCATCTTAG |
| NEPS2 | ATGGGAAATAAGAAAACGCTAGAGGGGAAGGTCGCCATTGTCACAGGTGGAGCATCTGGGATTGGGGAAACGGCGGCTAGAGT<br>ATTCGCCAACCTTGGGGCTAGGGCTGTTGTAATTGCCGATATTCAGAGTGAACTAGGCAGGGAGGTGGCTGAATCTATCGGCGC<br>CAAGAGATGCTCTTACGTTCAATGTGATATAGGGGACGAGGAACAGGTGAAATCCATGGTGGAATGGACCGCAACAACGTATG<br>GGGCATTAGACGTGATGTTCTGTAACGCCGGTATCATGTCTAAAGCCGAGAGTGCGCAGACCGTGCTAGAACTAGATATGTCAA<br>AGTTCGACGAGGTGATGAGGGTAAACACGCGTGGAACTTCTGCGTGTGTGAAGCAGGCCGCGAGAAAAATGGTAGAACTTGGG<br>ACTAAGGGTGGCGCGATAGTCTGTACTTCCTCTCCACTTGCCAGTCGTGGGGGCTATATAGATACAGATTACGTAATGTCCAAGC<br>ACGCGGTGATGGGTCTAGTAAGAAGTGCTTCCATGCAGCTTGGGGCACACGGCATCAGAGTAAATAGCGTTAGCCCAATGGCCG<br>TCTTAACTCCGCTGACGAGGAGAATGGGCCTGGCTACTCCCGCTGACGTTGAGAATGCATTTGGTAGGTTCACTTCCTTGAAAGG<br>TGTGGCCTTGACGGCTGAGCATGTTGCCGAGGCGGCTGCGTTCCTGGCGAGTGATGAGGCTGCTTTTATTACTGGACATGATCTG<br>ATGGTAGACGGGGGTTTGCTGTGTCTACCTTTCTTTGCCCCCACATCTTAA |
| LP2.T10 | TGCTCCAAGTGTGTGACTCCTTCATCTGACAACGTGCAACCCCTATCGCCATCGATTGTTTCTGCGGACGGTGTTGTCCTCATAGT<br>TTGGGCATGTTTCCCTTGTAGGTGTGAAACCACTTAGCTTCGCGCCGTAGTCCTAAAGGAAAACCTATGGACTTTGTTTCGGGTA |

| | |
|---|---|
| | GCACCAGGAATCTGAACCATGTGAATGTGGACGTGGCGCGCGTACACCTTAATCTCCGGTTCATGCTAGGGATGTGGCTGCATG<br>CTACGTTGACACACCTACACTGCTCTCCCCAATCGTGGAGTGAAGCGGGAAGTAGCGAGATGAAGTGTACGACCTGGCCGGAGC<br>CGTTCCGCATCGTCACGTGTTCGTTTACTGTTAATTGGTGGCACATAAGCAATATCGTAGTCCGTCAAATTCAGCCCTGTTATCCC<br>CGGCGTTATGTGTCAAATGGCGTAGAACTGGATTGACTGTTTGACGGTACCTGCTGATCGGTACGGTGACCGAGAATCTGTCGGG<br>CTATGTCACTAATACTTTCCAAACGCCCCGTATCGATGCTGAACGAATCGATGCACACTCACGTCTTTGAAGC |
| LV3 | AGGAATACTCTGAATAAAACAACTTATATAATAAAAATGC |
| LV5 | CCTCTTTATATTACATCAAAATAAGAAAATAATTATAACA |

**Table S2. Actual titers and standard deviation of 8HG accumulation for CYP/CPR combinatorial assessment**

| | | ***CPR*** | | | | | | | | | |
|---|---|---|---|---|---|---|---|---|---|---|---|
| | | ***Cr*** | ***At*** | ***Sr1*** | ***Sr8*** | ***Pk*** | ***Ns*** | ***Hc*** | ***Nd*** | ***Tc*** | ***Vv*** |
| ***G8H*** | **(-)** | 0.04 ± 0.05 | - | - | - | - | - | - | - | - | - |
| | ***Cr*** | 2.22 ± 0.24 | 1.19 ± 0.03 | 5.01 ± 0.34 | 3.30 ± 0.31 | 4.50 ± 0.64 | 7.36 ± 0.97 | 1.38 ± 0.03 | 1.86 ± 0.73 | 6.09 ± 0.59 | 5.57 ± 0.85 |
| | ***Gri*** | 0.00 ± 0.00 | - | - | 0.00 ± 0.00 | 0.00 ± 0.00 | 0.44 ± 0.01 | 0.03 ± 0.06 | 0.37 ± 0.05 | - | 0.00 ± 0.00 |
| | ***Pk*** | 0.00 ± 0.00 | 0.27 ± 0.46 | 0.00 ± 0.00 | 0.00 ± 0.00 | 0.11 ± 0.18 | 0.00 ± 0.00 | 0.04 ± 0.07 | 0.00 ± 0.00 | 0.11 ± 0.19 | 0.15 ± 0.26 |
| | ***Pp*** | 0.00 ± 0.00 | 0.00 ± 0.00 | 0.06 ± 0.10 | 0.00 ± 0.00 | 0.04 ± 0.06 | 0.01 ± 0.01 | 0.04 ± 0.08 | 0.04 ± 0.08 | 0.11 ± 0.16 | 0.05 ± 0.09 |
| | ***Ha*** | 0.03 ± 0.01 | 0.06 ± 0.05 | 0.02 ± 0.02 | 0.00 ± 0.00 | 0.50 ± 0.50 | 0.39 ± 0.40 | 0.00 ± 0.00 | 0.00 ± 0.00 | 0.00 ± 0.00 | 0.00 ± 0.00 |
| | ***St*** | 0.00 ± 0.00 | 0.00 ± 0.00 | 0.03 ± 0.05 | - | 0.03 ± 0.05 | 0.29 ± 0.18 | 0.02 ± 0.03 | 0.00 ± 0.00 | 0.19 ± 0.16 | 0.00 ± 0.00 |
| | ***Hr*** | 0.02 ± 0.04 | 0.00 ± 0.00 | 0.00 ± 0.00 | 0.18 ± 0.28 | 0.00 ± 0.00 | 0.17 ± 0.29 | 0.00 ± 0.00 | 0.25 ± 0.42 | 0.00 ± 0.00 | 0.01 ±0.02 |
| | ***Gr*** | 0.18 ± 0.05 | 0.45 ± 0.03 | 0.17 ± 0.29 | 0.22 ± 0.18 | - | 0.20 ± 0.35 | 0.00 ± 0.00 | 0.18 ± 0.16 | 0.15 ± 0.13 | 0.60 ± 0.13 |
| | ***Sa*** | 0.00 ± 0.00 | 0.08 ± 0.07 | 0.00 ± 0.00 | 0.01 ± 0.02 | 0.03 ± 0.05 | 0.04 ± 0.05 | 0.01 ± 0.07 | 0.00 ± 0.00 | 0.08 ± 0.00 | 0.09 ± 0.13 |
| | ***Jr*** | 0.00 ± 0.00 | - | 0.02 ± 0.04 | 0.00 ± 0.00 | 0.06 ± 0.11 | 0.33 ± 0.07 | 0.10 ± 0.09 | 0.00 ± 0.00 | 0.00 ± 0.00 | 0.11 ± 0.13 |
| | ***Rc*** | 0.00 ± 0.00 | 0.06 ± 0.10 | 0.09 ± 0.15 | 0.00 ± 0.00 | 0.00 ± 0.00 | 0.05 ± 0.06 | 0.03 ± 0.05 | 0.10 ± 0.03 | 0.24 ± 0.18 | 0.18 ± 0.31 |

| | | | | | | | | | | |
|---|---|---|---|---|---|---|---|---|---|---|
| ***Bs*** | 0.01 ± 0.01 | - | 0.13 ± 0.15 | 0.06 ± 0.07 | 0.34 ± 0.03 | 0.02 ± 0.03 | 0.00 ± 0.00 | 0.36 ± 0.07 | 0.00 ± 0.00 | 0.03 ± 0.03 |
| ***Vv*** | 0.01 ± 0.01 | 0.05 ± 0.07 | - | 0.03 ± 0.03 | 0.25 ± 0.08 | 0.00 ± 0.00 | - | 0.04 ±0.06 | 0.13 ± 0.23 | 0.00 ± 0.00 |

**Table S3. Molar conversion of geraniol to 8HG for various *Cr*G10H – *x*CPR combinations**

| G10 H | CPR | Geraniol [uM] | 10-hydroxygeraniol [uM] | Total count [uM] | % molar conversion |
|---|---|---|---|---|---|
| **(-)** | **(-)** | 47.4 | 0.2 | 47.6 | 0.4 |
| ***Cr*** | ***Cr*** | 36.6 | 13.1 | 49.7 | 26.3 |
| ***Cr*** | ***At*** | 20.4 | 7.0 | 27.4 | 25.6 |
| ***Cr*** | ***Sr1*** | 38.2 | 29.4 | 67.6 | 43.5 |
| ***Cr*** | ***Sr8*** | 20.1 | 19.4 | 39.5 | 49.1 |
| ***Cr*** | ***Pk*** | 9.1 | 26.4 | 35.5 | 74.5 |
| ***Cr*** | ***Ns*** | 14.6 | 43.3 | 57.8 | 74.8 |
| ***Cr*** | ***Hc*** | 20.5 | 8.1 | 28.6 | 28.4 |
| ***Cr*** | ***Nd*** | 41.0 | 10.9 | 52.0 | 21.0 |
| ***Cr*** | ***Tc*** | 66.1 | 35.8 | 101.9 | 35.1 |
| ***Cr*** | ***Vv*** | 71.7 | 32.7 | 104.4 | 31.3 |

**Table S4. Actual titers and standard deviation of geraniol accumulation for CYP/CPR combinatorial assessment**

| | | ***CPR*** | | | | | | | | | |
|---|---|---|---|---|---|---|---|---|---|---|---|
| | | ***Cr*** | ***At*** | ***Sr1*** | ***Sr8*** | ***Pk*** | ***Ns*** | ***Hc*** | ***Nd*** | ***Tc*** | ***Vv*** |
| ***G8H*** | **(-)** | 7.31 ± 0.80 | - | - | - | - | - | - | - | - | - |
| | ***Cr*** | 5.65 ± 0.63 | 3.15 ±0.68 | 5.89 ±0.99 | 3.10 ±0.30 | 1.40 ± 0.35 | 2.25 ± 0.24 | 3.16 ± 0.58 | 6.33 ± 0.96 | 10.20 ± 0.26 | 11.06 ± 1.87 |
| | ***Gri*** | 5.35 ± 0.20 | - | - | 4.12 ±0.29 | 0.28 ± 0.01 | 8.73 ± 0.09 | 0.27 ±0.06 | 8.90 ± 0.44 | - | 2.90 ± 0.75 |
| | ***Pk*** | 2.12 ± 0.30 | 1.38 ± 0.19 | 1.25 ± 0.18 | 1.73 ±0.17 | 2.14 ± 0.56 | 2.38 ± 0.48 | 2.34 ± 0.19 | 0.39 ± 0.02 | 1.18 ± 0.39 | 0.34 ± 0.29 |
| | | 0.25 ± 0.02 | 0.47 ± 0.16 | 0.29 ± 0.05 | 0.28 ± 0.04 | 0.37 ± 0.04 | 0.38 ± 0.09 | 0.32 ± 0.11 | 0.27 ± 0.01 | 0.33 ± 0.04 | 7.96 ± 0.58 |
| | ***Ha*** | 9.98 ± 0.52 | 6.74 ± 5.45 | 12.69 ± 1.28 | 14.92 ± 0.93 | 8.67 ± 0.55 | 9.96 ± 0.70 | 4.79 ± 0.14 | 4.64 ± 0.18 | 4.90 ± 0.05 | 17.38 ± 1.80 |
| | ***St*** | 0.09 ± 0.08 | 0.27 ± 0.09 | 0.35 ±0.04 | - | 0.16 ± 0.15 | 0.32 ± 0.20 | 0.34 ± 0.05 | 0.26 ± 0.01 | 0.48 ± 0.18 | 0.36 ± 0.07 |
| | ***Hr*** | 6.37 ± 0.43 | 3.43 ± 0.14 | 3.16 ± 0.18 | 8.42 ± 0.75 | 3.52 ± 0.15 | 3.20 ± 0.13 | 2.69 ± 0.15 | 3.97 ± 0.30 | 3.85 ± 0.09 | 3.10 ± |
| | ***Gr*** | 7.76 ± 0.19 | 8.38 ± 0.09 | 1.83 ± 0.98 | 8.25 ± 0.85 | - | 8.79 ± 0.37 | 3.42 ±0.12 | 1.80 ± 0.97 | 4.00 ± 1.19 | 13.59 ± 0.64 |
| | ***Sa*** | 8.50 ± 0.47 | 8.34 ± 0.48 | 12.83 ±1.15 | 9.28 ± 2.59 | 9.90 ± 0.90 | 9.60 ± 0.67 | 9.05 ± 0.76 | 9.28 ± 0.75 | 11.25 ± 1.82 | 8.75 ± 0.35 |
| | ***Jr*** | 5.15 ± 0.47 | 14.46 ± 1.47 | 15.71 ± 1.97 | 16.71 ± 1.85 | 12.94 ± 0.41 | 6.94 ± 0.44 | 5.44 ± 0.13 | 7.10 ± 0.67 | 16.40 ± 1.00 | 8.03 ± 1.37 |
| | ***Rc*** | 6.90 ± 0.74 | 5.39 ± 1.26 | 6.75 ± 0.35 | 4.38 ± 0.15 | 7.08 ± 0.63 | 9.14 ± 0.89 | 5.55 ± 0.43 | 5.51 ± 0.77 | 6.60 ± 0.50 | 7.50 ± 0.19 |
| | ***Bs*** | 5.86 ± 0.17 | - | 13.54 ± 0.71 | 12.05 ± 1.19 | 7.98 ± 1.04 | 12.88 ± 1.11 | 0.50 ± 0.15 | 7.47 ± 0.85 | 3.48 ± 0.23 | 4.35 ± 0.27 |

| | | | | | | | | | | |
|---|---|---|---|---|---|---|---|---|---|---|
| ***Vv*** | 3.85 ± 0.15 | 0.23 ± 0.07 | 2.83 ± 0.31 | - | - | 0.21 ± 0.02 | - | 0.31 ± 0.04 | 0.19 ± 0.04 | 0.30 ± 0.02 |

**Table S5. Molar conversion of geraniol to 8HG for multi-copy *CrG10H* assessment**

| **G10H** | **CPR** | **Geraniol [uM]** | **10-hydroxygeraniol [uM]** | **Total count [uM]** | **% molar conversion** |
|---|---|---|---|---|---|
| *Cr*G10H | *Cr* | 28.5 | 19.3 | 47.8 | 40.3 |
| 2x *Cr*G10H | *Cr* | 24.0 | 53.4 | 77.4 | 69.0 |
| 3 x *Cr*G10H | *Cr* | 9.3 | 67.8 | 77.1 | 88.0 |

### Table S6. Strains utilized in this study

| Description | Identifier | Integration: Relevant Genotype | Integration: Locus | Parent | Reference |
|---|---|---|---|---|---|
| **WT** | CEN.PK113 | - | - | *MATα; leu2-3, 112; ura3-52; MAL2-8C; SUC2* | EUROPHINS |
| **Geraniol optimization** | MD1 | $P_{TPI1}$-***erg20***$^{K197E}$*-loxP-kanMX* | ERG20 | CEN.PK113 | [61] |
| | MD2 | $P_{PGK}$-***tHMGR***-$T_{TDH2}$-$P_{TEF1}$-***IDI***-$T_{CYC1}$ | USERXII-2 | MD1 | This study |
| | MD3 | $P_{TEF2}$-***ObGES***-$T_{TDH2}$ | FgF20 | MD2 | This study |
| | MD4 | $P_{TEF2}$-***t63ObGES***-$T_{TDH2}$ | FgF20 | MD2 | This study |
| | MD5 | $P_{PYK1}$-***AgGPPS***-$T_{ENO2}$-$P_{PDC1}$-***CrCPR***-$T_{ADH2}$ | USERXII-3 | MD4 | This study |
| **G10H variants** | MD6 | $P_{TEF1}$-***CrG10H***-$T_{ADH1}$ | USERXII-5 | MD5 | This study |
| | MD7 | $P_{TEF1}$-***BsG10H***- $T_{ADH1}$ | USERXII-5 | MD5 | This study |
| | MD8 | $P_{TEF1}$-***EgG10H***- $T_{ADH1}$ | USERXII-5 | MD5 | This study |
| | MD9 | $P_{TEF1}$-***GrG10H***- $T_{ADH1}$ | USERXII-5 | MD5 | This study |
| | MD10 | $P_{TEF1}$-***GriG10H***- $T_{ADH1}$ | USERXII-5 | MD5 | This study |
| | MD11 | $P_{TEF1}$-***HaG10H***- $T_{ADH1}$ | USERXII-5 | MD5 | This study |
| | MD12 | $P_{TEF1}$-***HrG10H***- $T_{ADH1}$ | USERXII-5 | MD5 | This study |
| | MD13 | $P_{TEF1}$-***InG10H***- $T_{ADH1}$ | USERXII-5 | MD5 | This study |
| | MD14 | $P_{TEF1}$-***JrG10H***- $T_{ADH1}$ | USERXII-5 | MD5 | This study |
| | MD15 | $P_{TEF1}$-***PkG10H***- $T_{ADH1}$ | USERXII-5 | MD5 | This study |
| | MD16 | $P_{TEF1}$-***PpG10H***- $T_{ADH1}$ | USERXII-5 | MD5 | This study |
| | MD17 | $P_{TEF1}$-***RcG10H***- $T_{ADH1}$ | USERXII-5 | MD5 | This study |
| | MD18 | $P_{TEF1}$-***SaG10H***- $T_{ADH1}$ | USERXII-5 | MD5 | This study |
| | MD19 | $P_{TEF1}$-***StG10H***- $T_{ADH1}$ | USERXII-5 | MD5 | This study |
| | MD20 | $P_{TEF1}$-***VvG10H***- $T_{ADH1}$ | USERXII-5 | MD5 | This study |
| **CrG10H optimization** | MD36 | $P_{TEF1}$-***CrG10H***- $T_{ADH1}$ | Fgf16 | MD6 | This study |
| | MD37 | $P_{TEF1}$-***CrG10H***- $T_{ADH1}$ | Fgf24 | MD36 | This study |
| | MD38 | $P_{PGK}$-***CrG10H-CrCPR***- $T_{ADH1}$ | 106a | MD37 | This study |
| **CrG10H - CPR swap** | MD39 | *AtCPR* | *CrCPR* | MD6 | This study |
| | MD40 | *HcCPR* | *CrCPR* | MD6 | This study |
| | MD41 | *NdCPR* | *CrCPR* | MD6 | This study |
| | MD42 | *NsCPR* | *CrCPR* | MD6 | This study |
| | MD43 | *PkCPR* | *CrCPR* | MD6 | This study |
| | MD44 | *Sr1CPR* | *CrCPR* | MD6 | This study |
| | MD45 | *Sr8CPR* | *CrCPR* | MD6 | This study |
| | MD46 | *TcCPR* | *CrCPR* | MD6 | This study |
| | MD47 | *VvCPR* | *CrCPR* | MD6 | This study |

| | | | | | |
|---|---|---|---|---|---|
| **BsG10H - CPR swap** | MD48 | *HcCPR* | *CrCPR* | MD7 | This study |
| | MD49 | *NdCPR* | *CrCPR* | MD7 | This study |
| | MD50 | *NsCPR* | *CrCPR* | MD7 | This study |
| | MD51 | *PkCPR* | *CrCPR* | MD7 | This study |
| | MD52 | *Sr1CPR* | *CrCPR* | MD7 | This study |
| | MD53 | *Sr8CPR* | *CrCPR* | MD7 | This study |
| | MD54 | *TcCPR* | *CrCPR* | MD7 | This study |
| | MD55 | *VvCPR* | *CrCPR* | MD7 | This study |
| **GrG10H - CPR swap** | MD56 | *AtCPR* | *CrCPR* | MD9 | This study |
| | MD57 | *HcCPR* | *CrCPR* | MD9 | This study |
| | MD58 | *NdCPR* | *CrCPR* | MD9 | This study |
| | MD59 | *NsCPR* | *CrCPR* | MD9 | This study |
| | MD60 | *Sr1CPR* | *CrCPR* | MD9 | This study |
| | MD61 | *Sr8CPR* | *CrCPR* | MD9 | This study |
| | MD62 | *TcCPR* | *CrCPR* | MD9 | This study |
| | MD63 | *VvCPR* | *CrCPR* | MD9 | This study |
| **GriG10H - CPR swap** | MD64 | *HcCPR* | *CrCPR* | MD10 | This study |
| | MD65 | *NdCPR* | *CrCPR* | MD10 | This study |
| | MD66 | *NsCPR* | *CrCPR* | MD10 | This study |
| | MD67 | *PkCPR* | *CrCPR* | MD10 | This study |
| | MD68 | *Sr8CPR* | *CrCPR* | MD10 | This study |
| | MD69 | *VvCPR* | *CrCPR* | MD10 | This study |
| **HaG10H - CPR swap** | MD70 | *AtCPR* | *CrCPR* | MD11 | This study |
| | MD71 | *HcCPR* | *CrCPR* | MD11 | This study |
| | MD72 | *NdCPR* | *CrCPR* | MD11 | This study |
| | MD73 | *NsCPR* | *CrCPR* | MD11 | This study |
| | MD74 | *PkCPR* | *CrCPR* | MD11 | This study |
| | MD75 | *Sr1CPR* | *CrCPR* | MD11 | This study |
| | MD76 | *Sr8CPR* | *CrCPR* | MD11 | This study |
| | MD77 | *TcCPR* | *CrCPR* | MD11 | This study |
| | MD78 | *VvCPR* | *CrCPR* | MD11 | This study |
| **HrG10H- CPR swap** | MD79 | *AtCPR* | *CrCPR* | MD12 | This study |
| | MD80 | *HcCPR* | *CrCPR* | MD12 | This study |
| | MD81 | *NdCPR* | *CrCPR* | MD12 | This study |
| | MD82 | *NsCPR* | *CrCPR* | MD12 | This study |
| | MD83 | *PkCPR* | *CrCPR* | MD12 | This study |
| | MD84 | *Sr1CPR* | *CrCPR* | MD12 | This study |
| | MD85 | *Sr8CPR* | *CrCPR* | MD12 | This study |

| | | | | | |
|---|---|---|---|---|---|
| | MD86 | *TcCPR* | *CrCPR* | MD12 | This study |
| | MD87 | *VvCPR* | *CrCPR* | MD12 | This study |
| **JrG10H-CPR swap** | MD88 | *AtCPR* | *CrCPR* | MD14 | This study |
| | MD89 | *HcCPR* | *CrCPR* | MD14 | This study |
| | MD90 | *NdCPR* | *CrCPR* | MD14 | This study |
| | MD91 | *NsCPR* | *CrCPR* | MD14 | This study |
| | MD92 | *PkCPR* | *CrCPR* | MD14 | This study |
| | MD93 | *Sr1CPR* | *CrCPR* | MD14 | This study |
| | MD94 | *Sr8CPR* | *CrCPR* | MD14 | This study |
| | MD95 | *TcCPR* | *CrCPR* | MD14 | This study |
| | MD96 | *VvCPR* | *CrCPR* | MD14 | This study |
| **PkG10H-CPR swap** | MD97 | *AtCPR* | *CrCPR* | MD15 | This study |
| | MD98 | *HcCPR* | *CrCPR* | MD15 | This study |
| | MD99 | *NdCPR* | *CrCPR* | MD15 | This study |
| | MD100 | *NsCPR* | *CrCPR* | MD15 | This study |
| | MD101 | *PkCPR* | *CrCPR* | MD15 | This study |
| | MD102 | *Sr1CPR* | *CrCPR* | MD15 | This study |
| | MD103 | *Sr8CPR* | *CrCPR* | MD15 | This study |
| | MD104 | *TcCPR* | *CrCPR* | MD15 | This study |
| | MD105 | *VvCPR* | *CrCPR* | MD15 | This study |
| **PpG10H-CPR swap** | MD106 | *AtCPR* | *CrCPR* | MD16 | This study |
| | MD107 | *HcCPR* | *CrCPR* | MD16 | This study |
| | MD108 | *NdCPR* | *CrCPR* | MD16 | This study |
| | MD109 | *NsCPR* | *CrCPR* | MD16 | This study |
| | MD110 | *PkCPR* | *CrCPR* | MD16 | This study |
| | MD111 | *Sr1CPR* | *CrCPR* | MD16 | This study |
| | MD112 | *Sr8CPR* | *CrCPR* | MD16 | This study |
| | MD113 | *TcCPR* | *CrCPR* | MD16 | This study |
| | MD114 | *VvCPR* | *CrCPR* | MD16 | This study |
| **RcG10H-CPR swap** | MD115 | *AtCPR* | *CrCPR* | MD17 | This study |
| | MD116 | *HcCPR* | *CrCPR* | MD17 | This study |
| | MD117 | *NdCPR* | *CrCPR* | MD17 | This study |
| | MD118 | *NsCPR* | *CrCPR* | MD17 | This study |
| | MD119 | *PkCPR* | *CrCPR* | MD17 | This study |
| | MD120 | *Sr1CPR* | *CrCPR* | MD17 | This study |
| | MD121 | *Sr8CPR* | *CrCPR* | MD17 | This study |
| | MD122 | *TcCPR* | *CrCPR* | MD17 | This study |
| | MD123 | *VvCPR* | *CrCPR* | MD17 | This study |

| | | | | | |
|---|---|---|---|---|---|
| **SaG10H-CPR swap** | MD124 | *AtCPR* | *CrCPR* | MD18 | This study |
| | MD125 | *HcCPR* | *CrCPR* | MD18 | This study |
| | MD126 | *NdCPR* | *CrCPR* | MD18 | This study |
| | MD127 | *NsCPR* | *CrCPR* | MD18 | This study |
| | MD128 | *PkCPR* | *CrCPR* | MD18 | This study |
| | MD129 | *Sr1CPR* | *CrCPR* | MD18 | This study |
| | MD130 | *Sr8CPR* | *CrCPR* | MD18 | This study |
| | MD131 | *TcCPR* | *CrCPR* | MD18 | This study |
| | MD132 | *VvCPR* | *CrCPR* | MD18 | This study |
| **StG10H-CPR swap** | MD133 | *AtCPR* | *CrCPR* | MD19 | This study |
| | MD134 | *HcCPR* | *CrCPR* | MD19 | This study |
| | MD135 | *NdCPR* | *CrCPR* | MD19 | This study |
| | MD136 | *NsCPR* | *CrCPR* | MD19 | This study |
| | MD137 | *PkCPR* | *CrCPR* | MD19 | This study |
| | MD138 | *Sr1CPR* | *CrCPR* | MD19 | This study |
| | MD139 | *TcCPR* | *CrCPR* | MD19 | This study |
| | MD140 | *VvCPR* | *CrCPR* | MD19 | This study |
| **VvG10H-CPR swap** | MD141 | *AtCPR* | *CrCPR* | MD20 | This study |
| | MD142 | *NdCPR* | *CrCPR* | MD20 | This study |
| | MD143 | *NsCPR* | *CrCPR* | MD20 | This study |
| | MD144 | *Sr1CPR* | *CrCPR* | MD20 | This study |
| | MD145 | *TcCPR* | *CrCPR* | MD20 | This study |
| | MD146 | *VvCPR* | *CrCPR* | MD20 | This study |
| **3x-*Cr*G10H - CPR swap** | MD147 | *HcCPR* | *CrCPR* | MD37 | This study |
| | MD148 | *NsCPR* | *CrCPR* | MD37 | This study |
| | MD149 | *PkCPR* | *CrCPR* | MD37 | This study |
| | MD150 | *Sr1CPR* | *CrCPR* | MD37 | This study |
| | MD151 | *TcCPR* | *CrCPR* | MD37 | This study |
| **Nepetalactone production** | MD155 | $P_{TDH3}$-***10HGO***-$T_{Eno1}$ | *Ura3* | MD37 | This study |
| | MD156 | $P_{TDH3}$-***10HGO***-$T_{Eno1}$ – $P_{CCW12}$-***CrISY***-$T_{SSA1}$ | *Ura3* | MD37 | This study |
| | MD157 | $P_{TDH3}$-***10HGO***-$T_{Eno1}$ – $P_{CCW12}$-***CrISY***-$T_{SSA1}$ – $P_{PGK1}$- ***NEPS1***-$T_{ADH1}$ | *Ura3* | MD37 | This study |
| | MD158 | $P_{TDH3}$-***10HGO***-$T_{Eno1}$ – $P_{CCW12}$-***CrISY***-$T_{SSA1}$ – $P_{PGK1}$-***NEPS2***-$T_{ADH1}$ | *Ura3* | MD37 | This study |
| | MD159 | $P_{TDH3}$-***10HGO***-$T_{Eno1}$ – $P_{CCW12}$-***CrISY***-$T_{SSA1}$ – $P_{PGK1}$-***NEPS1***-$T_{ADH1}$ – $P_{HHF2}$– ***NEPS2***-$T_{PGK1}$ | *Ura3* | MD37 | This study |
| **OYE deletions** | MD152 | LP2.T10 | *Oye3* | MD6 | This study |
| | MD154 | LP2.T10 | *Oye2* | MD37 | This study |
| | MD160 | LP2.T10 | *Oye2* | MD155 | This study |

| | | | | | |
|---|---|---|---|---|---|
| | MD161 | LP2.T10 | *Oye3* | MD155 | This study |
| | MD162 | LP2.T10 | *Oye3* | MD160 | This study |
| | MD163 | LP2.T10 | *Oye2* | MD156 | This study |
| | MD164 | LP2.T10 | *Oye3* | MD156 | This study |
| | MD165 | LP2.T10 | *Oye3* | MD163 | This study |
| | MD166 | LP2.T10 | *Oye2* | MD157 | This study |
| | MD167 | LP2.T10 | *Oye3* | MD157 | This study |
| | MD168 | LP2.T10 | *Oye3* | MD166 | This study |
| | MD169 | LP2.T10 | *Oye2* | MD158 | This study |
| | MD170 | LP2.T10 | *Oye3* | MD158 | This study |
| | MD171 | LP2.T10 | *Oye3* | MD169 | This study |
| | MD172 | LP2.T10 | *Oye2* | MD159 | This study |
| | MD173 | LP2.T10 | *Oye3* | MD159 | This study |
| | MD174 | LP2.T10 | *Oye3* | MD172 | This study |

**Table S7. Oligonucleotides utilized in this study**

| **Amplicon** | **Primer ID** | **Sequence 5' -> 3'** | **Description** |
|---|---|---|---|
| tHMGR - IDI cassette | MD139 | AACGAAAAAGAAAAGAAAGACCATGTCATGTACGGGCAATCAGAATCTGTAACAAGCGCCACGCACAGATATTATAACATCTGCATAATAG | tHMGR cassette_F with 5' overhang |
| | MD168 | GTAAGGATTCGCGGTCCTCGAAAATTAAAAGTCCAACGCGCCTGTTGCTTGCGAAAAGCCAATTAGTGTG | tHMGR cassette_R with 3' homology to ISI cassette |
| | MD171 | ATCACGGATTTTCGATAAAGCACTTAGTATCACACTAATTGGCTTTTCGCAAGCAACAGGCGCGTTG | IDI cassette_F with 5' homology to tHMGR cassette |
| | MD140 | CTATTTCCTATAATAGAAATCCAAGTGGCAAAAGCGTTAGACGCAGTACAAGGACGCGTTAAGGCAAATTAAAGCCTTCGAGCG | IDI cassette_R with 3' overhang |
| ObGES/ tObGES | payge122 | AGGAATACTCTGAATAAAACAACTTATATAATAAAAATGCCTATATGGGGCCGTATACT | full length ObGES cassette_F with 5' homology to LV3 |

| | | | |
|---|---|---|---|
| | payge130 | TGTTATAATTATTTTCTTATTTTGATGTAATATAAAGAGGGCGAAAAGCCAATTAGT | full length ObGES cassette_R with 5' homology to LV5 |
| | MD146 | GTACTTGTTTTTAGAATATACGGTCAACGAACTATAATTAAAAAACAATGCAACACATGGAGGAGAGCAGC | Truncation of ObGES at position 63_F with 5' homology to TEF2p |
| AgGPPS - CrCPR cassette | MD158 | AGGAATACTCTGAATAAAACAACTTATATAATAAAAATGCAATGCTACTATTTTGGAGATTAATCTCAG | AgGPPS cassette_F with 5' homology to LV3 |
| | MD169 | TTTATCTTGCACATCACATCAGCGGAACATATGCTCACCCAGTCGCATGTAGGTATCATCTCCATCTCCCATATG | AgGPPS cassette_R with 3' homology to CrCPR cassette |
| | MD170 | TGCGGGCCACGACCACAGTGATATGCATATGGGAGATGGAGATGATACCTACATGCGACTGGGTGAG | CrCPR cassette_F with 5' homology to GPPS cassette |
| | payge126 | TGTTATAATTATTTTCTTATTTTGATGTAATATAAAGAGGTAGAATTATATAACTTGATGAGATGAG | CrCPR cassette_R with 3' homology to LV5 |
| BsG10H | MD178 | AACTTTTTTTACTTCTTGCTCATTAGAAAGAAAGCATAGCAATCTAATCTAAGTTTTAATAAAACAATGGACATCTTAAGCTC | BsG10H_F with 5' homology to TEF1p |
| | MD180 | AGGTAGACAAGCCGACAACCTTGATTGGAGACTTGACCAAACCTCTGGCGAAGAAGTCCATTATATTGCGATCGGAACGG | BsG10H_R with 3' homology to ADH1t |
| CrG10H | MD187 | AACTTTTTTTACTTCTTGCTCATTAGAAAGAAAGCATAGCAATCTAATCTAAGTTTTAATAAAACAATGGATTACTTAACTATCATATTGAC | CrG10H_F with 5' homology to TEF1p |
| | MD189 | AGGTAGACAAGCCGACAACCTTGATTGGAGACTTGACCAAACCTCTGGCGAAGAAGTCCATCACAGGGTAGAAGGCAC | CrG10H_R with 3' homology to ADH1t |
| GrG10H | MD195 | AACTTTTTTTACTTCTTGCTCATTAGAAAGAAAGCATAGCAATCTAATCTAAGTTTTAATAAAACAATGAGAGAAATGGATCTGC | GrG10H_F with 5' homology to TEF1p |
| | MD197 | AGGTAGACAAGCCGACAACCTTGATTGGAGACTTGACCAAACCTCTGGCGAAGAAGTCCATTATATCACAACTGGGATAGCCTG | GrG10H_R with 3' homology to ADH1t |
| GriG10H | MD199 | AACTTTTTTTACTTCTTGCTCATTAGAAAGAAAGCATAGCAATCTAATCTAAGTTTTAATAAAACAATGTTGGGCTCAAATAAGTC | GriG10H_F with 5' homology to TEF1p |
| | MD201 | AGGTAGACAAGCCGACAACCTTGATTGGAGACTTGACCAAACCTCTGGCGAAGAAGTCCATTAAAGGGATGTTGGGATGGC | GriG10H_R with 3' homology to ADH1t |
| HaG10H | MD203 | AACTTTTTTTACTTCTTGCTCATTAGAAAGAAAGCATAGCAATCTAATCTAAGTTTTAATAAAACAATGGGTTTTGTCATCGTTG | HaG10H_F with 5' homology to TEF1p |
| | MD205 | AGGTAGACAAGCCGACAACCTTGATTGGAGACTTGACCAAACCTCTGGCGAAGAAGTCCATTAATTTAGGGGTATCGGCACG | HaG10H_R with 3' homology to ADH1t |
| HrG10H | MD207 | AACTTTTTTTACTTCTTGCTCATTAGAAAGAAAGCATAGCAATCTAATCTAAGTTTTAATAAAACAATGGACTTCTTGGTATTCG | HrG10H_F with 5' homology to TEF1p |
| | MD209 | AGGTAGACAAGCCGACAACCTTGATTGGAGACTTGACCAAACCTCTGGCGAAGAAGTCCATTAAACGGGTACTATTAATAGTGGGTCA | HrG10H_R with 3' homology to ADH1t |
| JrG10H | MD215 | AACTTTTTTTACTTCTTGCTCATTAGAAAGAAAGCATAGCAATCTAATCTAAGTTTTAATAAAACAATGGACTTTTTGGGGTTG | JrG10H_F with 5' homology to TEF1p |

| | | | |
|---|---|---|---|
| | MD217 | AGGTAGACAAGCCGACAACCTTGATTGGAGACTTGACCAAACCTCTGGCGAAG AAGTCCATTATCTCAGAATTGGAACAGCACG | JrG10H_R with 3' homology to ADH1t |
| PkG10H | MD219 | AACTTTTTTTACTTCTTGCTCATTAGAAAGAAAGCATAGCAATCTAATCTAAGTT TTAATAAAACAATGGGCTTTTTGTCTGC | PkG10H_F with 5' homology to TEF1p |
| | MD221 | AGGTAGACAAGCCGACAACCTTGATTGGAGACTTGACCAAACCTCTGGCGAAG AAGTCCATTATACATGAAAACTTGGTATCGCTAG | PkG10H_R with 3' homology to ADH1t |
| PpG10H | MD223 | AACTTTTTTTACTTCTTGCTCATTAGAAAGAAAGCATAGCAATCTAATCTAAGTT TTAATAAAACAATGGATTTTTTAACTATAATTCTGGG | PpG10H_F with 5' homology to TEF1p |
| | MD225 | AGGTAGACAAGCCGACAACCTTGATTGGAGACTTGACCAAACCTCTGGCGAAG AAGTCCATTACAGTGGTATTGGTACAGCC | PpG10H_R with 3' homology to ADH1t |
| RcG10H | MD227 | AACTTTTTTTACTTCTTGCTCATTAGAAAGAAAGCATAGCAATCTAATCTAAGTT TTAATAAAACAATGATGGACCTTCTTGTAG | RcG10H_F with 5' homology to TEF1p |
| | MD229 | AGGTAGACAAGCCGACAACCTTGATTGGAGACTTGACCAAACCTCTGGCGAAG AAGTCCATTAGACCTGATTGGGTATGGC | RcG10H_R with 3' homology to ADH1t |
| SaG10H | MD231 | AACTTTTTTTACTTCTTGCTCATTAGAAAGAAAGCATAGCAATCTAATCTAAGTT TTAATAAAACAATGGATTTCCTTGGTTTTATAC | SaG10H_F with 5' homology to TEF1p |
| | MD233 | AGGTAGACAAGCCGACAACCTTGATTGGAGACTTGACCAAACCTCTGGCGAAG AAGTCCATTAACGAATGGGTACAGCACATAATG | SaG10H_R with 3' homology to ADH1t |
| StG10H | MD235 | AACTTTTTTTACTTCTTGCTCATTAGAAAGAAAGCATAGCAATCTAATCTAAGTT TTAATAAAACAATGGAGTATGTAAACATCCTTC | StG10H_F with 5' homology to TEF1p |
| | MD237 | AGGTAGACAAGCCGACAACCTTGATTGGAGACTTGACCAAACCTCTGGCGAAG AAGTCCATTAGTATCCCAGAAGCTGTAGGG | StG10H_R with 3' homology to ADH1t |
| VvG10H | MD239 | AACTTTTTTTACTTCTTGCTCATTAGAAAGAAAGCATAGCAATCTAATCTAAGTT TTAATAAAACAATGGATTACACTCCGC | VvG10H_F with 5' homology to TEF1p |
| | MD241 | AGGTAGACAAGCCGACAACCTTGATTGGAGACTTGACCAAACCTCTGGCGAAG AAGTCCATTAGGGTTTAGTCGGTACGG | VvG10H_R with 3' homology to ADH1t |
| Tef1p | Payge121 | AGGAATACTCTGAATAAAACAACTTATATAATAAAAATGCATAGCTTCAAAATG TTTCTACTC | Tef1 cassette_F with 5' homology to LV3 |
| | MD157 | GCATAGCAATCTAATCTAAGTTTTAAT | Tef1 cassette_R |
| Adh1t | MD175 | TGGACTTCTTCGCCAGAG | ADH1 cassette_F |
| | Payge124 | TGTTATAATTATTTTCTTATTTTGATGTAATATAAAGAGGGCATGCCGGTAGAG | AtCPR_F with 5' homology to PDC1p |
| AtCPR | MD335 | TATCTTCTACTCATAACCTCACGCAAAATAACACAGTCAAATCAAAAACAATGA GTTCATCCAGTTCTTCTTC | AtCPR_F with 5' homology to PDC1p |
| | MD336 | ATGCTTGATAATGAAAACTATAAATCGTAAAGACATAAGATCCGCCTAGCTCTT CAGCCCCAGAC | AtCPR_R with 3' homology to ADH2t |
| HcCPR | MD339 | TATCTTCTACTCATAACCTCACGCAAAATAACACAGTCAAATCAAAAACAATGG AGAGCAGTAGCGTAAAG | HcCPR_F with 5' homology to PDC1p |

| | | | |
|---|---|---|---|
| | MD340 | GCATGCTTGATAATGAAAACTATAAATCGTAAAGACATAAGATCCGCCTAGCTCTTCAGCCCCATACG | HcCPR_R with 3' homology to ADH2t |
| NdCPR | MD343 | TATCTTCTACTCATAACCTCACGCAAAATAACACAGTCAAATCAAAAACAATGCAAGAAGCGTCCAGC | NdCPR_F with 5' homology to PDC1p |
| | MD344 | GCATGCTTGATAATGAAAACTATAAATCGTAAAGACATAAGATCCGCCTAGCTCTTCAGCCCCAAACATC | NdCPR_R with 3' homology to ADH2t |
| NsCPR | MD345 | TATCTTCTACTCATAACCTCACGCAAAATAACACAGTCAAATCAAAAACAATGGCAAGTAATCTTGTAAATTCTTTAG | NsCPR_F with 5' homology to PDC1p |
| | MD346 | TAGGCATGCTTGATAATGAAAACTATAAATCGTAAAGACATAAGATCCGCCTAGCTCTTCAGCCCCACAC | NsCPR_R with 3' homology to ADH2t |
| Sr1CPR | MD349 | TATCTTCTACTCATAACCTCACGCAAAATAACACAGTCAAATCAAAAACAATGCAATCAGATTCAGTCAAAGTC | Sr1CPR_F with 5' homology to PDC1p |
| | MD350 | TAGGCATGCTTGATAATGAAAACTATAAATCGTAAAGACATAAGATCCGCCTACCAAACATCTCTTAAGTATCTTCCAG | Sr1CPR_R with 3' homology to ADH2t |
| Sr8CPR | MD351 | TATCTTCTACTCATAACCTCACGCAAAATAACACAGTCAAATCAAAAACAATGCAATCTAACTCCGTGAAG | Sr8CPR_F with 5' homology to PDC1p |
| | MD352 | TAGGCATGCTTGATAATGAAAACTATAAATCGTAAAGACATAAGATCCGCCTATTACCAAACGTCACGGAGG | Sr8CPR_R with 3' homology to ADH2t |
| TcCPR | MD353 | TATCTTCTACTCATAACCTCACGCAAAATAACACAGTCAAATCAAAAACAATGCAGACTTCCGAAGTCAAAATATC | TcCPR_F with 5' homology to PDC1p |
| | MD354 | TAGGCATGCTTGATAATGAAAACTATAAATCGTAAAGACATAAGATCCGCCTAGCTCTTCAGCCCCATACG | TcCPR_R with 3' homology to ADH2t |
| PkCPR | MD375 | AATTATTATCTTCTACTCATAACCTCACGCAAAATAACACAGTCAAATCAAAAACAATGCAATCCACGC | PkCPR_F with 5' homology to PDC1p |
| | MD376 | TAGGCATGCTTGATAATGAAAACTATAAATCGTAAAGACATAAGATCCGCTTACCAAACGTCTCTTAAATACCTTC | PkCPR_R with 3' homology to ADH2t |
| VvCPR | MD377 | AATTATTATCTTCTACTCATAACCTCACGCAAAATAACACAGTCAAATCAAAAACAATGCAATCCTCCTCTG | VvCPR_F with 5' homology to PDC1p |
| | MD378 | TAGGCATGCTTGATAATGAAAACTATAAATCGTAAAGACATAAGATCCGCTTACCATACATCACGCAAGTATC | VvCPR_R with 3' homology to ADH2t |
| 10HGO | DT180 | GCATCGTCTCATCGGTCTCATATGGCCAAGTCTCCAGAAGTTG | Amplify 10HGO_F |
| | DT181 | ATGCCGTCTCAGGTCTCAGGATTTAAGCAGACTTTAAGGTATTAGCGAC | Amplify 10HGO_R |
| ISY | MD520 | ATGTCTTGGTGGTGGAAAC | Amplify ISY_F |
| | MD521 | TTAGGGTATGAACCTATAGTCACG | Amplify ISY_R |
| NEPS1 | MD522 | ATGGCGTCCACGGCAAATC | Amplify NEPS1_F |

| | | | |
|---|---|---|---|
| | MD523 | CTAAGATGGAGCGAAAAAGGGTAAG | Amplify NEPS1_R |
| NEPS2 | MD524 | ATGGGAAATAAGAAAACGCTAG | Amplify NEPS2_F |
| | MD525 | TTAAGATGTGGGGGCAAAG | Amplify NEPS2_R |
| OYE2 | MD295 | TCCAGATATAGAATAAATCATCATATTAAGCTAAATATAGACGATAATATAGTATCGATATGCTCCAAGTGTGTGACTC | LP2.T10_F with 5' homology to OYE2 locus |
| | MD296 | ATATATTCATTAATTATATAAATTAGAAGAAAAAGAAATGGTGCTACAAAGTACGGTTAAGCTTCAAAGACGTGAGTGTG | LP2.T10_R with 3' homology to OYE2 locus |
| OYE3 | MD299 | TAATTAAAAATATGGCAGGAATATGAAAAATACATAACATCAATGTCTTTATTCATGATTTGCTCCAAGTGTGTGACTC | LP2.T10_F with 5' homology to OYE23locus |
| | MD300 | TTCAGAGATTCTACTCTTGACCACTGTTTCGTGTAGCCGCTCAAGGTTTATTTCTTTCTTGCTTCAAAGACGTGAGTGTG | LP2.T10_R with 3' homology to OYE3 locus |
| USERXII-3 | MD5 | GAAACTAACCCGATGGGACAATTAC | UP region_F |
| | MD6 | GCATTTTTATTATATAAGTTGTTTTATTCAGAGTATTCCTTACCCCTTATTATAATGATTAATACTTACATCATAG | UP region_R with 3' homology to LV3 |
| | MD7 | CCTCTTTATATTACATCAAAATAAGAAAATAATTATAACAGGAAGTTTTGCAGATGAAGTGC | DOWN region_F with 5' homology to LV5 |
| | MD8 | CCAACGCATTTACAAACCACG | DOWN region_R |
| USERXII-5 | MD159 | CAATCTGGCGGCTTGAGTTC | UP region_F |
| | MD160 | GCATTTTTATTATATAAGTTGTTTTATTCAGAGTATTCCTTACCGGTTCTGCCACCTC | UP region_R with 3' homology to LV3 |
| | MD161 | CCTCTTTATATTACATCAAAATAAGAAAATAATTATAACACTCAGAAGTTTGACAGCAAGC | DOWN region_F with 5' homology to LV5 |
| | MD162 | ATACTAGAGTTAACTGATGGTCTTAAACAG | DOWN region_R |
| FgF16 | MD249 | GCATTTTTATTATATAAGTTGTTTTATTCAGAGTATTCCTTCCGTTAATTCGGGTTTCAATC | UP region_R with 3' homology to LV3 |
| | MD250 | TTCGTGAAACACGTGGGATA | UP region_F |
| | MD251 | TTGTTGGGATTCCATTGTGATTAAG | DOWN region_R |
| | MD252 | CCTCTTTATATTACATCAAAATAAGAAAATAATTATAACATGCCTACGCAACACTTTAGC | DOWN region_F with 5' homology to LV5 |
| FgF24 | MD257 | GCATTTTTATTATATAAGTTGTTTTATTCAGAGTATTCCTGGATCACCTCGCCCTG | UP region_R with 3' homology to LV3 |
| | MD258 | GGCTGAACAACAGTCTCTCC | UP region_F |

| | | | |
|---|---|---|---|
| | MD259 | GGCGGTAGTGATAACCATTCTC | DOWN region_R |
| | MD260 | CCTCTTTATATTACATCAAAATAAGAAAATAATTATAACAGTTGAAGTCGCCTGGTAGC | DOWN region_F with 5' homology to LV5 |
| 106a | LB1543 | CCCTGTTCATGGCAACGTCACC | UP region_F |
| | LB1544 | GCATTTTTATTATATAAGTTGTTTTATTCAGAGTATTCCTCGTATCACAACCGACGATCCG | UP region_R with 3' homology to LV3 |
| | LB1545 | CCTCTTTATATTACATCAAAATAAGAAAATAATTATAACACCTGGTCAAACTTCAGAACTAA | DOWN region_F with 5' homology to LV5 |
| | LB1546 | GGATTAAGTAACAGATACAGACATCAC | DOWN region_R |

### Table S8. Integration sites utilized in this study

| ***S. cerevisiae locus*** | **Cas9 target site** |
|---|---|
| USERXII_2 | TCGAGAGAGTCGCCGATAGT |
| FgF20 | GTTAGAGCTGTTACAAGTTA |
| USERXII_5 | TTGTCACAGTGTCACATCAG |
| FgF16 | TGTACCAAAAGTTATCCTGT |
| FgF24 | CCTATTGGACAAGATTTACG |
| 106a | ATACGGTCAGGGTAGCGCCC |
| *Cr*CPR | TTTGTTGAAGGCAACGACAG |
| OYE3 | CTCATCAGACCAAATCCCAG |
| OYE2 | GTAACCCCCAGATTGTGGAG |
